# YdbL directly modulates YdbH-YnbE bridge formation to maintain *Escherichia coli* outer membrane homeostasis

**DOI:** 10.64898/2026.03.16.712125

**Authors:** Clare De’Ath, Sujeet Kumar, Sachi Chhibber, Benjamin F. Cooper, Emilia K. Taylor, Emily Jones, Thomas Lanyon-Hogg, Jani R. Bolla, Christina Redfield, Natividad Ruiz, Georgia L. Isom

## Abstract

Gram-negative bacteria pose a threat to global healthcare mainly because their outer membrane (OM) provides an intrinsic barrier to many antimicrobials. Key to this barrier function is the asymmetric structure of the OM, with phospholipids constituting the inner leaflet and lipopolysaccharides the outer leaflet. Although the mechanism of phospholipid transport between the inner membrane (IM) and OM remains poorly understood, recent studies implicate TamB, YhdP, and YdbH as functionally redundant proteins mediating this process in *Escherichia coli*. Accordingly, collective loss of these three paralogs is lethal and any one of them is sufficient for growth. YdbH is anchored to the IM and its periplasmic repeating β-sheet groove domain interacts with the OM lipoprotein YnbE via β-strand augmentation to form an intermembrane bridge. Additionally, YnbE multimerizes, and the periplasmic protein YdbL is proposed to modulate YnbE multimerization to facilitate its stacking on the C-terminus of YdbH. Here, we demonstrate that excess YdbL specifically inhibits the function of the YdbH-YnbE complex since overexpression of *ydbL* causes lethality in the Δ*yhdP* Δ*tamB* double mutant but the presence of both ydbH and ynbE in trans abrogates this lethality. We resolve high-resolution structural data for YdbL and ascertain its interaction site with the YnbE C-terminal *α*-helix, with residues mediating this interface highly conserved and critical for YdbL function. Finally, we show that YdbL is protected from degradation by the protease DegP when complexed with YnbE. Overall, our data supports a model in which YdbL ensures proper YdbH-YnbE intermembrane bridge formation by directly interacting with YnbE.

## Introduction

The Gram-negative bacterial cell envelope consists of an inner membrane (IM), a thin peptidoglycan cell wall housed within the periplasm, and an outer membrane (OM). The OM is a permeability barrier that confers intrinsic levels of resistance to antibiotics and detergents (1), making it an attractive target for novel therapeutic strategies. The OM is asymmetric, consisting of glycerophospholipids at the inner leaflet and lipopolysaccharide (LPS) dominating the outer leaflet. Tight electrostatic interactions between adjacent LPS molecules are crucial to the OM barrier function (2), protecting bacteria from a wide range of antimicrobials (3). OM proteins (OMPs) form a near-static network of stable β-barrels contributing to membrane integrity in addition to facilitating the import and export of hydrophilic substrates (4–6). Most cellular lipoproteins are tethered to the inner or outer leaflet of the OM resulting in exposure to the periplasm or cell surface, respectively.

To build the OM, components must be transported from the cytoplasm and IM across the aqueous periplasm (2). Bulk phospholipid transport between the IM and OM remained elusive until the recent discovery that members of the AsmA-like protein family play a role in this process (7–9). The bacterial AsmA-like proteins are ancestors of the repeating β-groove (RBG) superfamily, which includes eukaryotic bridge-like lipid-transfer proteins (BLTPs) that form bridges between organelles at membrane contact sites to transport phospholipids (10, 11). In *Escherichia coli*, there are six AsmA-like paralogues with a similar predicted architecture comprising a single-pass IM transmembrane helix followed by a periplasmic domain of varying length. Crucially, this periplasmic domain is composed of a stacked, β-sheet groove lined with hydrophobic residues that can accommodate lipophilic substrates, such as the acyl tails of glycerophospholipids, to shield them from the aqueous periplasm. The three largest paralogues by molecular weight in *E. coli*, YhdP, TamB and YdbH, are functionally redundant in maintaining OM lipid homeostasis, with a triple knockout being synthetically lethal to cells (8, 9). The absence of both YhdP and TamB causes pleiotropic defects related to the OM including sensitivity to MacConkey agar (bile salts), as well as formation of mucoid colonies due to the activation of the Rcs stress response and upregulation of capsule formation (8, 9). Additionally, depletion of YhdP in a Δ*tamB* Δ*ydbH* double mutant results in an accumulation of phospholipids at the IM, suggesting a role for all three proteins in anterograde phospholipid transport (9).

Bridge-like protein structures between the IM and OM must extend around 250 Å to span the full periplasmic width (12–14). YhdP is of sufficient length to do so (15), and TamB interacts with the Omp85-family β-barrel protein TamA, forming an envelope-spanning complex (16–18). YdbH is significantly shorter than YhdP and TamB, with a predicted periplasmic domain length of ∼180 Å, and thus requires additional components to span the cell envelope.

YdbH is encoded in an operon with the OM lipoprotein YnbE and the soluble periplasmic protein YdbL. Introducing multiple copies of YdbH-YnbE-YdbL rescues the mucoidy phenotype of *E. coli* lacking TamB and YhdP (19), further demonstrating overlap in function of these systems. Genetic and biochemical evidence has revealed YdbH forms a functional complex with YnbE via a β-strand augmentation at YdbH’s C-terminus, as predicted by AlphaFold-Multimer (19, 20). Additionally, *in vivo* photocrosslinking has shown that YnbE also self-multimerizes to form higher molecular weight YnbE-YnbE adducts (19). Together, these data suggest that IM-tethered YdbH can interact directly with OM-tethered YnbE to form a continuous bridge to the OM, possibly with YnbE multimerization forming a stack of β-strands to sufficiently span the periplasm in order to transport phospholipids (Fig. 1(a)). In support of this model, *Pseudomonas aeruginosa* YdbH fused to a long C-terminal region of YhdP can function in the absence of YnbE (20).

**Figure 1.**
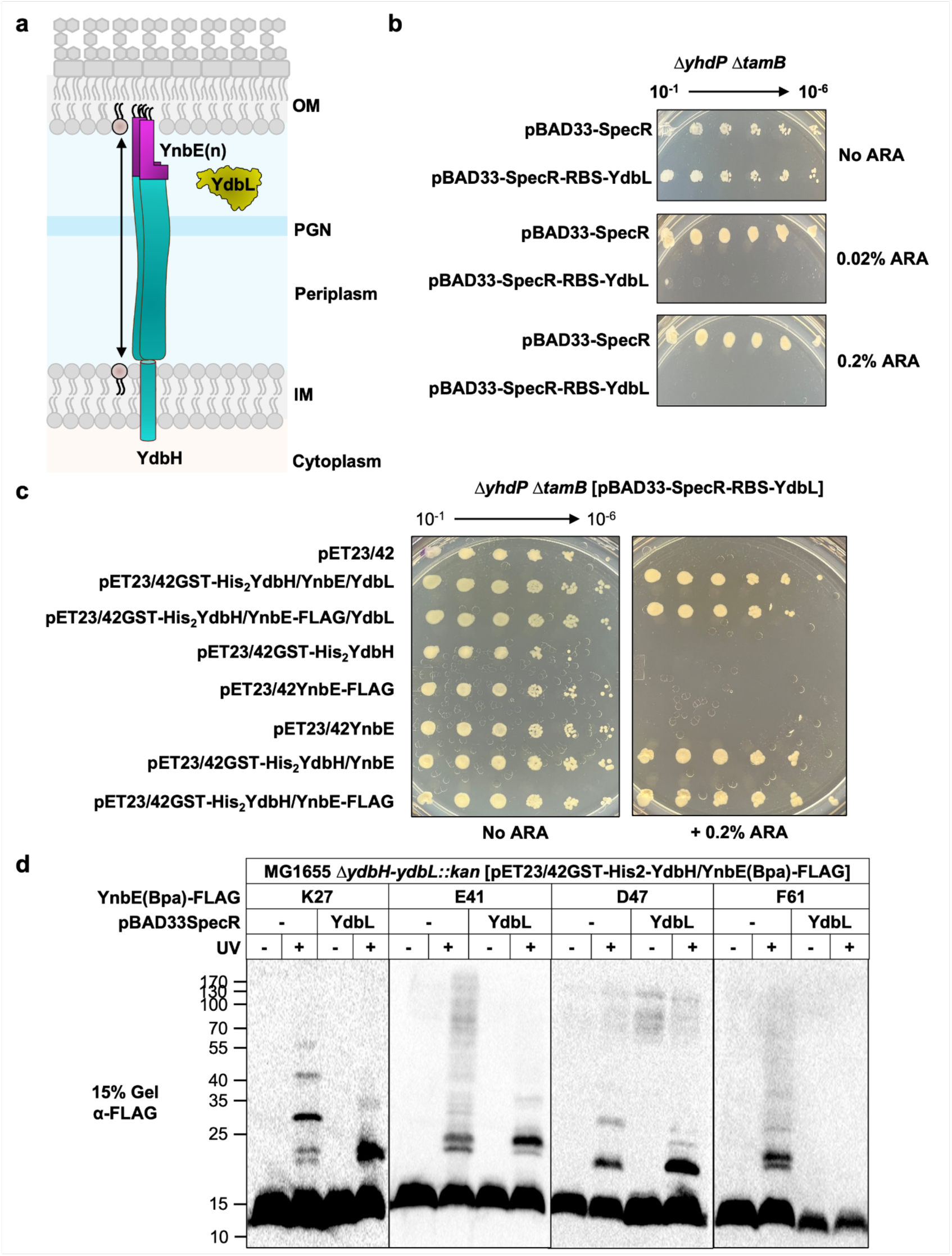
Overexpression of YdbL from pBAD33 causes lethality in Δ*yhdP* Δ*tamB* double mutant and affects the conformation and/or complex formation of YnbE. **a.** The *ydbH* gene is co-transcribed with *ynbE* and *ydbL*, encoding an OM lipoprotein and a periplasmic protein, respectively. YdbH forms an intermembrane bridge through its interaction with a YnbE monomer or oligomer (YnbEn) of unknown stoichiometry, while YdbL is proposed to facilitate complex formation by restricting YnbE multimerization**. b.** The effect of YdbL overexpression on the growth of the Δ*yhdP* Δ*tamB* double mutant was tested by spotting serially diluted cultures of *ΔyhdP ΔtamB* double mutant strains carrying either the pBAD33-SpecR vector or pBAD33-SpecR-RBS-YdbL plasmid on LB agar containing spectinomycin, with or without 0.02% or 0.2% arabinose. In the presence of 0.2% arabinose, Δ*yhdP* Δ*tamB* mutant carrying pBAD33RBS-YdbL plasmid showed no visible growth. **c.** The rescue of lethality caused by YdbL overexpression in the Δ*yhdP* Δ*tamB* double mutant was tested by spotting serially diluted cultures of these strains carrying both the pBAD33-SpecR-RBS-YdbL and pET23/42-derived plasmids encoding *ydbH*, *ynbE* and *ydbL* alleles on LB agar containing spectinomycin and ampicillin, with or without 0.2% arabinose. Co-overexpression of *ydbH* and *ynbE*, but not their individual overexpression from pET23/42-derived plasmids, fully rescued the lethality caused by pBAD33-SpecR-RBS-YdbL in Δ*yhdP* Δ*tamB* double mutant. **d.** Immunoblot analysis showed that the presence of YdbL affects the conformation and/or complex formation of YnbE. Exponentially growing Δ*ydbH-ydbL::kan* cells producing plasmid-encoded YnbE(pBPA)-FLAG variants with substitutions at K27, E41, D47 and F61, and carrying either pBAD33SpecR vector or pBAD33SpecR-RBS-YdbL, were treated (or not) with UV light for 30 minutes at room temperature. Whole-cell lysate samples were prepared as described in the Material and Methods section, and proteins were separated on 15% SDS-polyacrylamide gel and probed with α-FLAG antibodies by immunoblotting. Molecular mass marker (in kDa) are shown on the left. All data are representative of three independent experiments.

In *E. coli*, YnbE and YdbH are essential in the absence of TamB and YhdP, but YdbL is not (19). However, multicopy *ydbL* alongside *ydbH* and *ynbE* is required to suppress mucoidy (resulting from activation of the Rcs stress response) of the Δ*yhdP* Δ*tamB* double mutant (19). Furthermore, overproduction of YdbL results in a dominant-negative effect, leading to killing of the Δ*yhdP* Δ*tamB* double mutant. In the absence of YdbL, YdbH levels decrease whereas YnbE multimerization significantly increases (19). Based on these observations, YdbL has been proposed to possess a chaperone-like function to control runaway multimerization of YnbE, thereby influencing the efficiency of complex formation with YdbH (Fig. 1(a)). The dominant-negative phenotype conferred by increased levels of YdbL also suggests that relative levels of these proteins are important, with YdbL possibly sequestering YnbE and preventing its interaction with YdbH. However, this model relies on YdbL’s dominant-negative phenotype being specific to YdbH-YnbE function and assumes a direct interaction between YdbL and YnbE, which has yet to be determined.

In this study, using a combination of genetic, biochemical, and structural methods, we elucidate a direct interaction of YdbL with YnbE. Genetic analyses reveal YdbL specifically affects the function and complex formation of YdbH-YnbE. Through our extensive structural and functional characterization, we identify a conserved binding interface for YdbL with YnbE which is vital for YdbL function. We also consider the cellular mechanisms by which relative protein levels are controlled, defining YdbL as a substrate for periplasmic protease DegP *in vitro*. Our work contributes detailed molecular insights into the YdbH-YnbE-YdbL complex to more comprehensively understand how these proteins interact to function as one of the putative phospholipid transport systems vital for OM biogenesis and homeostasis in *E. coli*.

## Results

### Overproduction of YdbL negatively impacts YdbH-YnbE function in Δ*yhdP* Δ*tamB* cells

Since viability of the Δ*yhdP* Δ*tamB* double mutant depends on a functional YdbH/YnbE complex, we hypothesized that excess YdbL would negatively affect YdbH/YnbE function (19). Because previous studies showing the negative effect of YdbL relied on cells with higher gene dose of *ydbL* than *ydbH-ynbE* (19), we constructed the pET23/42GST-His2YdbH/YnbE/RBS-FLAG-YdbL plasmid in which we introduced a strong RBS directly upstream of *ydbL* to increase only the relative translation of *ydbL* alongside an N-terminal FLAG-tag after the signal sequence to allow immunodetection (SI Fig. 1(a)). The presence of an N-terminal FLAG tag on mature YdbL did not affect its function since FLAG-YdbL behaved like untagged YdbL (SI Fig. 1(b)). We found that the pET23/42GST-His2YdbH/YnbE/RBS-FLAG-YdbL plasmid was only able to partially suppress the envelope defects of the Δ*yhdP* Δ*tamB* double mutant suggesting that excess YdbL negatively affects YdbH/YnbE function (SI Fig. 1(b)). To determine whether this was indeed due to excess YdbL, we monitored production of FLAG-YdbL from pET23/42-derived plasmids using immunoblotting. Unfortunately, we could not detect FLAG-YdbL produced from pET23/42-derived plasmids under conditions in which we could easily detect GST-His2YdbH and YnbE-FLAG from pET23/42GST-His2YdbH/YnbE-FLAG/YdbL (19). However, we were able to specifically detect FLAG-YdbL produced from pET23/42-derived plasmids in rhamnose-induced whole-cell lysate samples from KRX strains, which express an inducible T7 RNA polymerase under the control of a tightly controlled rhamnose promoter. As expected, higher levels of FLAG-YdbL were detected when produced from pET23/42GST-His2YdbH/YnbE/RBS-FLAG-YdbL plasmid as compared with the pET23/42GST-His2YdbH/YnbE/FLAG-YdbL plasmid (SI Fig. 1(c)). These findings are consistent with the idea that elevated YdbL levels negatively impact the function of YdbH/YnbE.

### Increased levels of both YdbH and YnbE suppresses the lethality caused by excess YdbL in Δ*yhdP* Δ*tamB* cells

To further probe how YdbL affects the function of YdbH/YnbE, we co-transformed the various pET23/42-containing strains with an arabinose inducible pBAD33-SpecR-RBS-YdbL plasmid (21). This enabled us to control production of YdbL in the cells by varying arabinose concentration, while overcoming the complication of transcriptional and translational coupling in the *ydbH-ydbL* operon. As expected, pBAD33-SpecR-RBS-YdbL caused arabinose-dependent lethality in the Δ*yhdP* Δ*tamB* double mutant (Fig. 1(b), SI Fig. 2) (19). We next tested whether pET23/42-encoded *ydbH* and *ynbE* could individually or collectively rescue the lethality caused by pBAD33-SpecR-RBS-YdbL in the Δ*yhdP* Δ*tamB* double mutant in the presence of arabinose. We found that the presence of both *ydbH* and *ynbE* was required to rescue growth (Fig. 1(c)). Together these results show that YdbL specifically affects the function of the YdbH/YnbE complex.

We next investigated whether levels of functional FLAG-YdbL were affected by the presence of YdbH and YnbE. Thus, we sought to detect FLAG-YdbL produced from pET23/42 plasmid, while ydbH and ynbE were expressed in trans either individually or together from pBAD33-SpecR. Immunoblot analysis of whole-cell lysates from KRX Δ(*ydbH-ynbE-ydbL*) cells revealed that FLAG-YdbL was detected at higher levels when YdbH or YnbE were produced in trans either alone or together from pBAD33-SpecR (SI Fig. 3). This result suggests that YdbH and/or YnbE may stabilize YdbL, overcoming a negative effect that YdbL exerts on the function and/or assembly of the YdbH-YnbE complex in Δ*yhdP* Δ*tamB* mutants. Thus, it is likely that YdbL physically interacts with YdbH and YnbE. Notably, AlphaFold-Multimer predicted that YdbH, YnbE and YdbL form a complex in which the C-terminus of YdbH interacts with both YnbE and YdbL (19). In this study, we focused on probing the functional and physical interaction between YnbE and YdbL.

### Overproduction of YdbL decreases multimerization of YnbE

YnbE can multimerize and interact with YdbH in cells forming a complex of unknown stoichiometry (19). The multimerization of YnbE and formation of YdbH-YnbE complexes can occur in the absence of YdbL, although its absence can alter the pattern of YnbE multimerization observed at specific YnbE sites using an *in vivo* site-specific photocrosslinking analysis (19). Here we monitored the effect of producing higher amounts of YdbL using the same crosslinking approach, by incorporating the unnatural amino acid *p*-benzoyl-L-phenylalanine (*p*BPA) at sites of interest using amber codon suppression, followed by UV-dependent crosslinking with nearby C-H bonds (22). We performed these experiments in cells producing four YnbE(*p*BPA)-FLAG variants (with *p*BPA substitutions at residues K27, E41, D47, or F61) from pET23/42GST-His2-YdbH/YnbE(Bpa)-FLAG and carrying an arabinose-inducible *ydbL* allele in pBAD33-SpecR. Immunoblot analysis from whole-cell lysate samples showed that YdbL decreased the detection of higher molecular weight (HMW) species of YnbE(*p*BPA)-FLAG variants (Fig. 1(d)). We also observed an increase in the signal of a potential YnbE dimer (∼20kDa band) in the YnbE(*p*BPA)-FLAG variants (K27, E41 and D47) when YdbL was present (Fig. 1(d)). Interestingly, the YnbE(F61*p*BPA)-FLAG variant showed an overall decrease in signal of both crosslinked and uncrosslinked species in the presence of YdbL (Fig. 1(d)). Collectively, these data show that the presence of YdbL influences the conformation and/or formation of multimeric YnbE complexes and that, when in excess, it decreases YnbE multimerization.

### The X-ray crystal structure of YdbL resolved to 2.12 Å at room temperature

We resolved the structure of YdbL by X-ray crystallography (XRC). In short, we produced YdbL lacking its signal peptide (residues 22-108) carrying an N-terminal 6xHis-tag and TEV cleavage site in the cytoplasm. We purified this protein to produce untagged, mature YdbL which eluted as a monodisperse peak by SEC (SI Fig. 4(a)). Purified protein was submitted to XRC trials via the VMXi beamline (23–25) to produce crystals of exemplar quality for *in situ* room temperature diffraction experiments (Table S6). The structure of YdbL was determined at 2.12 Å, permitting the modelling of all residues including their side-chain densities (Fig. 2(a)). Alignment with the AlphaFold2-predicted structure revealed near identical models, with a very low root mean square deviation (RMSD) below 1.0 Å across all atom pairs (SI Fig. 4(b)). Differences were primarily observed at the C-terminal loop and side-chain orientations. The PDBePISA server (26, 27) was used to assess crystal contact interfaces to investigate whether the C-terminal loop shift had biological relevance or was a crystal-packing artefact (SI Fig. 4(c)). With an exemplary fit to the electron density as shown in SI Fig. 4(d), this C-terminal region adopts an alternative conformation to pack in the crystal lattice and is thus reflective of a genuine mobile region.

**Figure 2.**
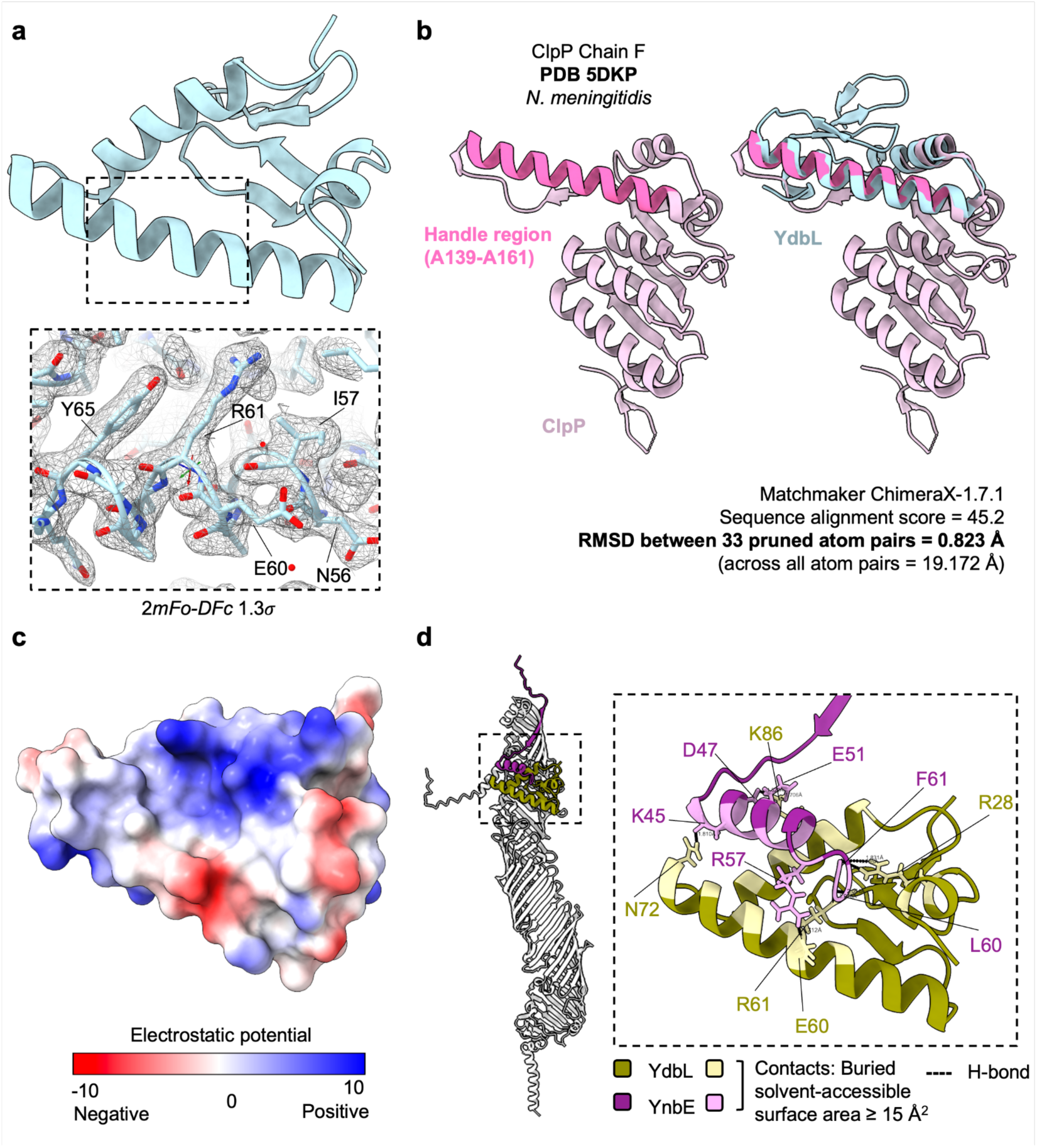
X-ray crystal structure of YdbL resolved to 2.1 Å at room temperature. **a.** XRC structure of YdbL model presented as ribbon secondary structure. The boxed region depicts an example (residues N56 - Y65) of the fit of side chains to the electron density map. **b.** Structural homology screening via the Dali server reveals YdbL shares a homologous fold to the handle region (A139 - A161 (31)) in the ATP-dependent proteolytic subunit of ClpP protease. The left panel shows the top hit structure from Table S8, ATP-dependent proteolytic subunit of ClpP protease from *N. meningitidis* (pastel pink) (PDB 5KDP, chain F) with the handle region highlighted (deep pink). The right panel shows the overlay of YdbL to ClpP using MatchMaker from ChimeraX-1.7.1. YdbL (light blue) overlaps directly with the conserved handle region of ClpP essential for establishing functional protease conformation. Sequence alignment scores and RMSD between atom pairs are given below. **c.** Coulombic electrostatic potential reported from ChimeraX v1.7.1 reveals a strong patch of positive charge in YdbL to form a putative binding pocket for an interaction partner. **d.** MultiFold2 prediction of YdbH (white), YnbE (purple) and YdbL (gold) in 1:1:1 stoichiometry. All three proteins form a complex at the C-terminus of YdbH, with YnbE aligning via a β-strand to the β-sheet groove of YdbH and a C-terminal α-helix aligning to the putative binding pocket of YdbL. This YdbL-YnbE interaction defined in the dotted box is shown with the contact region (buried solvent-accessible surface area ≥15 Å2) shaded in pastel yellow and pastel pink for YdbL and YnbE respectively and hydrogen bonds with a black dotted line. All other side chains not forming hydrogen bonds are hidden for clarity.

To evaluate whether our crystal structure accurately represented the conformation of YdbL in solution, SEC-SAXS experiments were conducted (SI Fig. 5(a-e)) with data reduction statistics given in Table S7. Given the structure was determined from cytoplasmically-expressed protein, experiments were conducted in parallel with periplasmically-expressed protein to ensure the structure matched protein expressed in its native subcellular location. Solution-state structures from both cytoplasmic and periplasmic proteins aligned harmoniously with the solid-state and thus we concluded our crystal structure to be a physiologically relevant model SI Fig. 5(f)).

### YdbL possesses a conserved binding interface with the C-terminus of YnbE

To assess how YdbL may interact with YnbE, we investigated the structural characteristics of our model in further detail. First, we assessed YdbL’s structural homology using the Dali server (28), which revealed statistically significant homology to the handle region of ClpP (ATP-dependent proteolytic subunit, PDB 5DKP (29)) (Table S8). ClpP is a highly-conserved serine protease in the bacterial cytoplasm responsible for degrading mainly misfolded proteins (30, 31). YdbL aligns via its two longest *α*-helices (residues A48-N72 and V76-R90) with an RMSD of 0.823 Å across 33 atom pairs that encompass ClpP’s handle region (A139 - A161) (Fig. 2(b)). The handle region is vital to mediating *α*-helical interactions that produce a functional oligomeric assembly of ClpP subunits with its catalytic triad at a suitable geometry (31). Importantly, the long *α*-helical portion directly associates with its counterpart from other subunits. Although we do not have evidence of YdbL oligomerization (as determined by SEC-SAXS *in vitro*), this homology suggests that this region of YdbL may be prone to mediating *α*-helical interactions with other proteins. We also evaluated the coulombic electrostatic potential which highlighted a putative binding pocket with a distinct region of strongly positive charge that could favor binding of a complementary negatively charged partner (Fig. 2(c)).

Next, we wanted to determine if YdbH or YnbE could occupy this putative binding site in YdbL. As both YdbH and YnbE remain structurally elusive, we used *in silico* methods to understand how they may interact with YdbL. Having validated the AlphaFold2 YdbL structure experimentally, MultiFold2 (32, 33) was used to simulate YdbL with YdbH and YnbE as conducted previously in AlphaFold-Multimer by Kumar *et al.* (19) (Fig. 2(d), SI Fig. 6(a(i)). Results between AlphaFold-Multimer and MultiFold2 were consistent, with all three proteins predicted to form a complex via the C-terminus of YdbH interacting with both YnbE and YdbL. YnbE forms a β-strand (residues I30-I38) which aligns to the β-sheet groove of YdbH to promote a direct interaction between IM-tethered YdbH and OM-tethered YnbE for YdbH’s putative transport function (19) (SI Fig. 6(a(i))). The other structured region of YnbE, a C-terminal *α*-helical portion (residues K45-T56), occupied the putative binding pocket of YdbL (SI Fig. 6(a(i))). Although simulating YnbE and YdbL together in the absence of YdbH resulted in loss of YnbE’s β-strand secondary structure to a disordered region and lengthening of its C-terminal *α*-helix (SI Fig. 6(a(ii)), (b)), the interacting region with YdbL remained the same (K45-F61), suggesting it may be key for YnbE-YdbL complex formation (SI Fig. 6(b)).

To experimentally validate the MultiFold2 prediction, we sought to use purified YdbL and YnbE to probe their interaction. The intrinsic self-multimerization of YnbE makes it a challenging target to isolate (19). Therefore, a C-terminal YnbE peptide (residues E39-F61) was designed to avoid any β-strand mediated self-multimerization as shown in Fig. 3(a) and SI Fig. 6(a(iii)). We employed solution NMR to experimentally probe this predicted interface which revealed significant amide backbone chemical shift changes indicative of a high-affinity interaction across multiple residues (Fig. 3(b))

**Figure 3.**
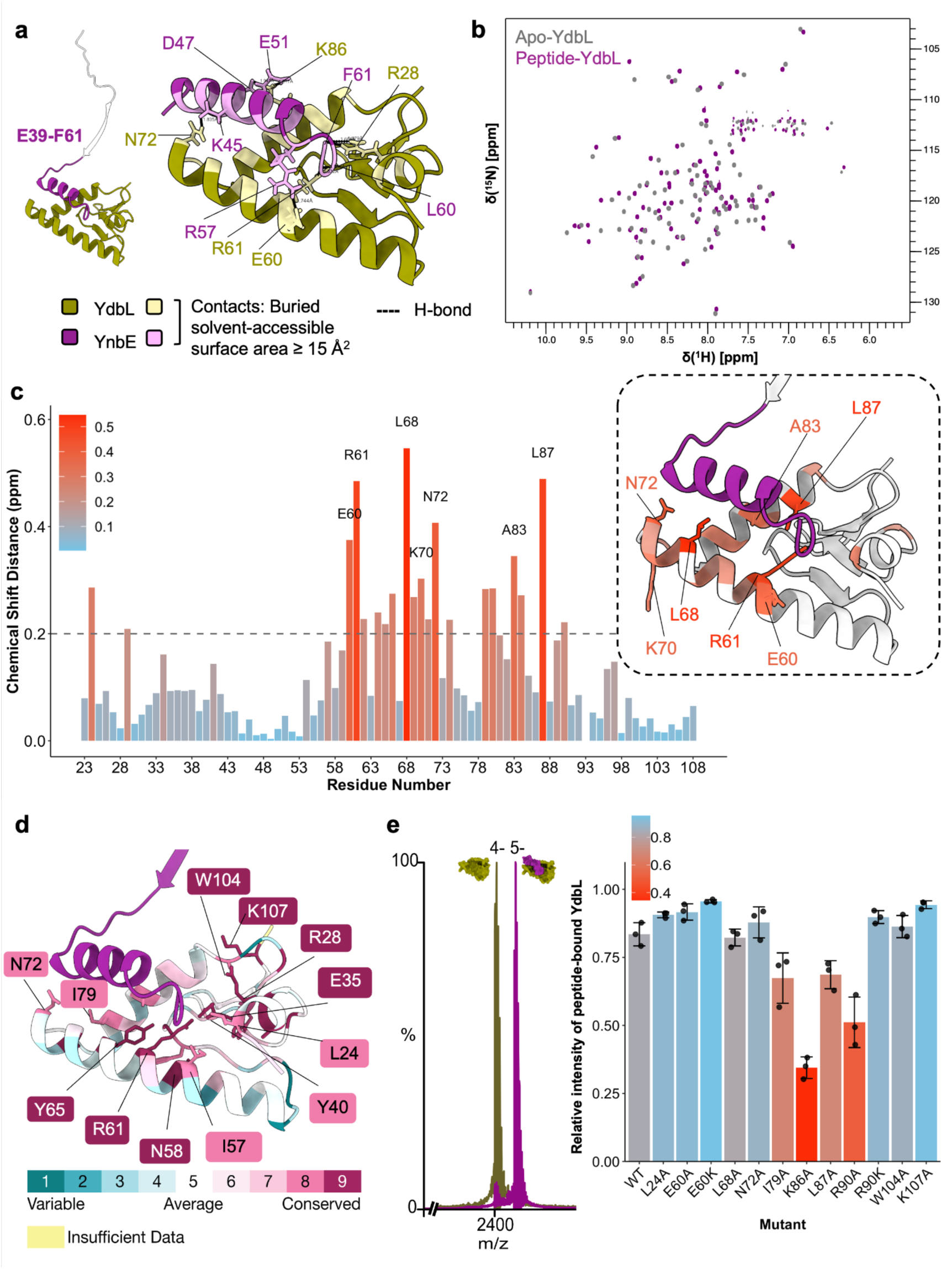
YdbL interacts with the C-terminal *α*-helix of YnbE. **a.** MultiFold2 prediction of YdbL with YnbE peptide (E39-F61) matches well with predictions for full length YnbE in complex with YdbL. **b.** ^1^H-^15^N HSQC NMR spectra showing amide backbone peaks of apo-YdbL (grey) and peptide-bound YdbL (purple) reveal that the C-terminal YnbE peptide interacts with YdbL. **c.** Amide backbone chemical shift changes, in ppm, between apo- and peptide-bound YdbL. Experimental shifts above 0.2 ppm are colored according to the plot and the largest experimental chemical shifts (ppm > 0.3) are labelled with side chains shown. These concur with the predicted interface when mapped to the MultiFold2 model. Table S9 lists the amide chemical shift changes for all assigned peaks. Assigned chemical shift data can be found in the BMRB under deposition codes 53587 (apo-YdbL) and 53588 (peptide-bound YdbL)). **d.** Consurf analysis of YdbL XRC structure reveals that evolutionarily conserved residues line up with the binding interface. All residues with a score above 8 have side chains shown and are labelled with the exception of conserved alanine and glycine residues. **e.** Native mass spectrum of wild type YdbL in the presence of YnbE peptide. Main charge states for wild-type series and peptide-bound series are labelled in gold and purple, respectively. The accompanying bar chart reveals the relative intensity of peptide-bound YdbL for wild-type and available mutants, shown in SI Fig. 9 (N=3).

The combined ^1^H-^15^N amide chemical shift changes revealed that the C-terminal YnbE peptide interacted with YdbL in a manner that is consistent with the MultiFold2-predicted interface of full-length YnbE in complex with YdbL (Fig. 3(c), SI Fig. 6(a(ii-iii), b)). Residues E60, R61, L68, K70, N72, A83 and L87 showed significant amide shift changes above 0.3 ppm. Although residue K70 was not predicted to directly interact with YnbE by the MultiFold2 complex prediction (Fig. 2(d)), it is flanked by interacting residues L68 and N72 which likely conferred significant amide backbone shift changes in this region upon peptide binding. Additionally, residues E60, R61 and N72 were also predicted by MultiFold2 to mediate direct hydrogen bonds with YnbE as shown in Fig. 2(d).

Next, we probed the relative secondary structure upon peptide binding to identify possible conformational changes not reflected by *in silico* predictions. We employed the Talos-N server (34) to assign the secondary structure of YdbL in its apo- and peptide-bound forms. This revealed a predominantly *α*-helical secondary structure with two β-hairpin loop regions reflective of the resolved XRC structure (Fig. 2(a), SI Fig. 7(a)). Further analysis comparing chemical shift changes between apo- and peptide-bound YdbL as per SI Fig. 7(b-c) did not reveal significant structural rearrangements of YdbL upon binding, with the observed results indicative of minor or localized rearrangements mediated by local environment changes and side chain rearrangements (SI Fig. 7(a, b, c)).

One notable discrepancy between the experimental data and *in silico* prediction pertained to R28, a residue situated on the short *α*-helical loop at the N-terminus of YdbL. By MultiFold2, R28 was predicted to form a hydrogen bond with F61 from YnbE; however, R28 did not exhibit a significant chemical shift change of the amide proton by NMR. We postulated this lack of chemical shift change may have been masked by the combination of its localization in a more mobile region and its long side chain mediating an interaction distal to its backbone. Therefore, whilst there was strong overlap between experimentally determined and predicted interacting YdbL residues, we used both analyses to inform our functional studies.

### Structure-function analysis reveals residues mediating the YdbL-YnbE interface are functionally important

We next probed the functional relevance of YdbL residues by testing the ability of pET23/42 plasmid-encoded *flag-ydbL* mutant alleles to cause lethality in the Δ*tamB* Δ*yhdP* double mutant. Given that wild-type YdbL causes a synthetically lethal phenotype in this genetic background, we postulated that any mutants that formed colonies produced non-functional YdbL variants. As shown in Table 1, YdbL residues mediating the YnbE-YdbL interface were chosen for mutagenesis based on structural predictions and experimental data. Namely, L24, R28, E60, R61, Y65, L68, N72, I79, K86, L87 and R90 were changed to alanine. Glutamic acid and arginine residues were also changed to aspartic acid and lysine respectively to maintain charge but alter side-chain properties. All substitutions rendered YdbL non-functional with the exception of N72A, which partially affected function, and E60A, E60D and R90K which did not cause detectable defects. Of the residues that were essential for function, all except L68 and R90 were highly conserved, scoring between 8 and 9 by Consurf (35) (Fig. 3(d)). Substituting other highly conserved YdbL residues not predicted to be involved in the YnbE-YdbL interaction, namely E35A, E35D, Y40A, N58A, W104A and K107A, also resulted in non- or partially- functional YdbL variants compared to non-conserved residue control variants E51A and D77A. This group included W104A and K107A, predicted to form part of the YdbH-YdbL interaction site. Therefore, the functionality of YdbL is not solely dependent on its interaction with YnbE. To further characterize the non-functional variants of YdbL, we compared the growth curve of Δ*tamB* Δ*yhdP* double mutant carrying non-functional variants of YdbL encoded on pET23/42 plasmid with empty vector control. We found that cells producing the YdbL(N72A) variant, and to a lesser extent YdbL (E35D), (R90A), (K86A), (L87A), (E35A), (L68A) variants, lysed faster than those carrying the vector control, suggesting that residual YdbL activity was still present in these variants (Table 1, SI Fig. 8(a)). Immunoblot analysis of whole-cell lysate samples of KRX strains carrying pET23/42-derived plasmids encoding FLAG-YdbL variants revealed that most, but not all, non-functional or partially functional variants exhibited higher levels of YdbL than wild type, whereas functional variants showed levels comparable to those of wild-type YdbL (SI Fig. 8(b)). Together, these results suggest that YdbL function depends primarily on its ability to interact with YdbH or YnbE.

**Table 1.** Functional studies of YdbL variants based on their interaction interface with YnbE and evolutionary conservation score. Highlights in orange reveal when there was a significant predicted contact or a H-bond between YdbL and YdbH or YnbE, large amide proton shift in ppm above 0.2 or loss of function. Consurf scores (35) above 7 are bolded. Functionality of variants was assessed by their ability to confer lethality in the Δ*tamB* Δ*yhdP* double mutant when they were produced from the pET23/42 plasmid. Only plasmids encoding partially functional and non-functional *ydbL* alleles yielded viable transformants in Δ*tamB* Δ*yhdP* double mutant cells. Transformants producing YdbL (W104A) and (K107A) showed severe growth defects on LB ampicillin plates, while YdbL (N72A), and to a lesser extent YdbL (E35D), (R90A), (K86A), (L87A), (E35A), (L68A) variants, showed significant increases in lysis compared to pET23/42 vector control when grown in liquid cultures (see SI figure 8(a)) and were therefore categorized as partial-loss of function variants.

| pET23/42 | Predicted interaction (MultiFold2) | Significant experimental chemical shift? (N-H HSQC NMR) | Consurf score | Functional phenotype? |
| --- | --- | --- | --- | --- |
| Vector | - | - | - | N/A |
| YdbL | - | - | - | Yes |
| FLAG-YdbL | - | - | - | Yes |
| FLAG-YdbL(L24A) | YnbE-YdbL contact | (Y)<br>>0.2 ppm | **8** | No |
| FLAG-YdbL(R28A) | YnbE-YdbL H-bond | N<br><0.1 ppm | **9** | No |
| FLAG-YdbL(E35A) | - | N<br><0.1ppm | **9** | Partially |
| FLAG-YdbL(E35D) | - | N<br><0.1ppm | **9** | Partially |
| FLAG-YdbL(Y40A) | - | N<br><0.1 ppm | **8** | No |
| FLAG-YdbL(E51A) | - | N<br><0.1 ppm | 4 | Yes |
| FLAG-YdbL(N58A) | - | N<br><0.1 ppm | **9** | No |
| FLAG-YdbL(E60A) | YnbE-YdbL contact | Y<br>>0.3 ppm | 6 | Yes |
| FLAG-YdbL(E60D) | YnbE-YdbL contact | Y<br>>0.3 ppm | 6 | Yes |
| FLAG-YdbL(R61A) | YnbE-YdbL H-bond | Y<br>>0.4 ppm | **9** | No |
| FLAG-YdbL(R61K) | YnbE-YdbL<br>H-bond | Y<br>>0.4 ppm | 9 | No |
| FLAG-YdbL(Y65A) | YnbE-YdbL<br>contact | (Y)<br>>0.2 ppm | 9 | No |
| FLAG-YdbL(L68A) | YnbE-YdbL<br>contact | Y<br>>0.4 ppm | 7 | Partially |
| FLAG-YdbL(N72A) | YnbE-YdbL<br>H-bond | Y<br>>0.4 ppm | 8 | Partially |
| FLAG-YdbL(D77A) | - | N<br><0.1 ppm | 3 | Yes |
| FLAG-YdbL(I79A) | YnbE-YdbL<br>contact | Y<br>>0.2 ppm | 8 | No |
| FLAG-YdbL(K86A) | YnbE-YdbL<br>H-bond | N<br><0.1 ppm | 7 | Partially |
| FLAG-YdbL(L87A) | YnbE-YdbL<br>contact | Y<br>>0.4 ppm | 7 | Partially |
| FLAG-YdbL(R90A) | YnbE-YdbL<br>H-bond | (Y)<br>>0.2 ppm | 6 | Partially |
| FLAG-YdbL(R90K) | YnbE-YdbL<br>H-bond | (Y)<br>>0.2 ppm | 6 | Yes |
| FLAG-YdbL(W104A) | YdbH-YdbL<br>contact | N<br><0.1 ppm | 9 | Partially |
| FLAG-YdbL(K107A) | YdbH-YdbL<br>contact | N<br><0.1 ppm | 9 | Partially |

To test whether the loss of function caused by changes at the YnbE-YdbL interface was directly linked to its ability to bind YnbE, we used native mass spectrometry (native MS) to monitor binding events between YdbL mutant proteins and the C-terminal YnbE peptide (Fig. 3(e)). Although other E60 variants did not cause detectable defects in function, an E60K variant was introduced to swap charge at this site predicted to shift significantly by NMR. We also included variants with the W104A and K107A substitutions situated at the predicted YdbH-YdbL interface as negative controls. Not all mutant proteins could be stably produced to sufficient yields for native MS as shown in SI Fig. 9(a), namely those with the R28A, R61A, R61K and Y65A substitutions and were therefore excluded from the native MS analyses. For those YdbL variants that we could purify (L24A, E60A, E60K, L68A, N72A, I79A, K86A, L87A, R90A, R90K, W104A and K107A), we analysed them with equimolar amounts of YnbE peptide and acquired native MS data. As shown in Fig. 3(e) and SI Fig. 9(b), the wild type spectrum display two different charge state distributions with measured masses (9,599士0.98 Da and 12,342士0.71 Da) corresponding to wild-type YdbL and 1:1 ratio of YdbL-YnbE peptide. Upon mixing YdbL variants with the YnbE peptide, we observed a reduction in peptide binding most prominently for those with the I79A, K86A, L87A and R90A substitutions (Fig. 3(e), SI Fig. 9(b) and SI Table (S5)). Therefore, even single changes can affect the YdbL-YnbE binding interface. These YdbL residues are situated closer to the C-terminus of YnbE and thus may indicate a region particularly important for binding and sensitive to perturbation. Overall, our genetic and biochemical analyses reveal that YnbE-YdbL interaction is required for YdbL function.

### YdbL is degraded by periplasmic protease DegP in vitro

Thus far, our data has revealed that excess intracellular levels of YdbL relative to YdbH and YnbE are toxic in cells relying on YdbH-YnbE for growth, with this dominant-negative phenotype specific to YdbH-YnbE function. Although we cannot rule out other types of regulation of YdbL, it may be important for cells to rapidly alter YdbL levels post-translationally by protease-mediated degradation.

Of the periplasmic proteases researched so far, DegP is the best characterized, operating as a dual chaperone-protease primarily associated with degrading misfolded OMPs (36–40). To degrade its targets, DegP forms a modular assembly that recognizes the C-terminal residues of target proteins as degrons (41–43). Although no clear consensus motif has been identified that encompasses all DegP substrates, hydrophobic C-terminal residues and significant domains of hydrophobicity are considered to be well-recognized by DegP for degradation (41).

Thus, we analyzed the hydrophobicity of YdbL, observing two significant regions of hydrophobicity (Fig. 4(a(ii))). The first region situated close to the YnbE-binding site is mediated predominantly via the two α-helices A48-N72 and V76-A89. In addition, the combination of the V33-L44 loop with the C-terminal residues W104-F108 generated another significant area of hydrophobicity. The presence of these hydrophobic regions, especially the hydrophobic C-terminal residues, made YdbL a reasonable candidate for DegP degradation. Furthermore, as described above, changes in this region increased levels of YdbL (SI Fig. 8b).

**Figure 4.**
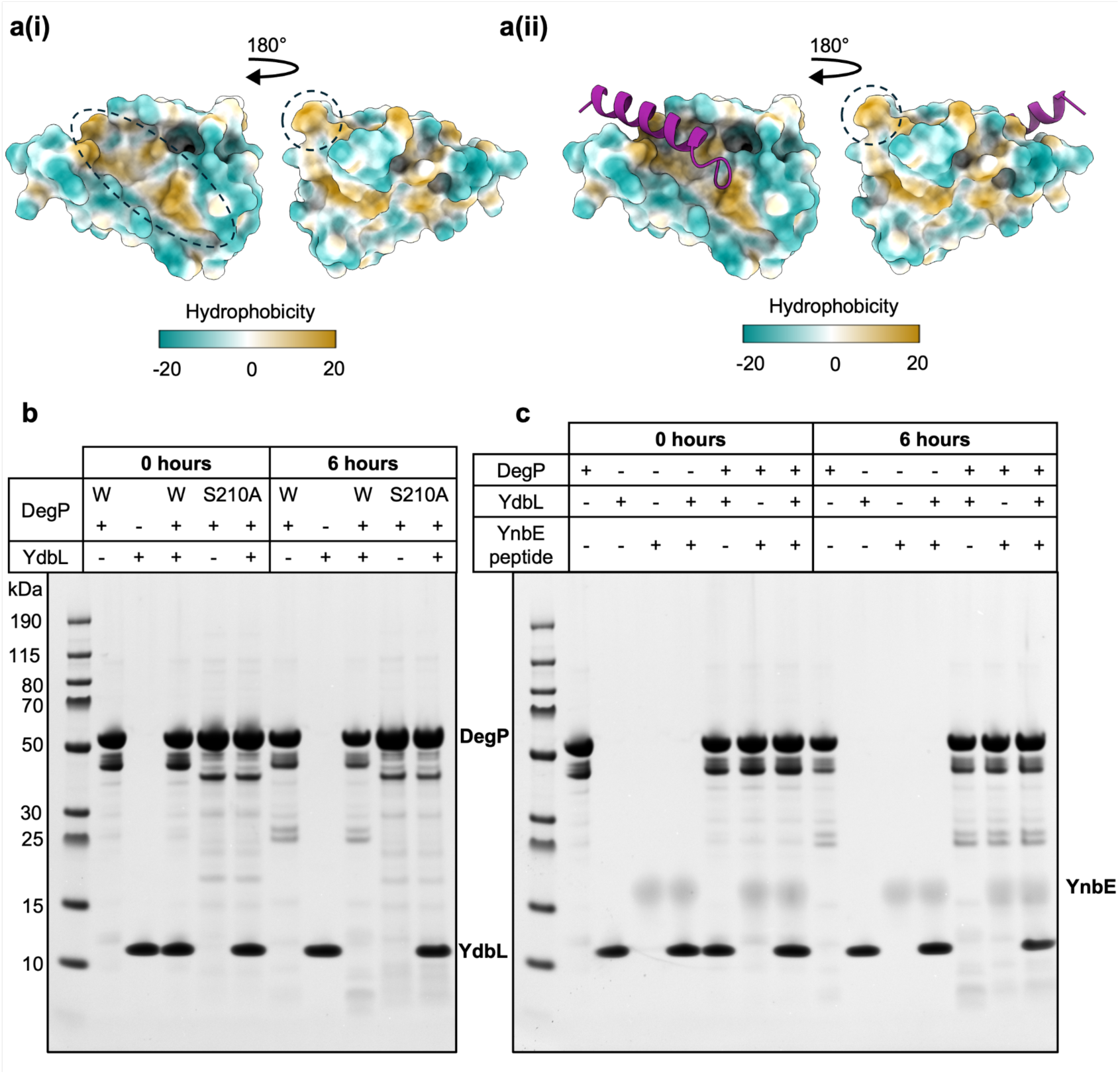
DegP degrades YdbL *in vitro* but addition of the YnbE peptide protects YdbL from degradation. **a. (i)** Hydrophobicity of YdbL calculated using ChimeraX v1.7.1 is shown in the top panel. Patches of hydrophobicity are found around the protein, including the C-terminus visualised with the dashed circle**. (ii)** The hydrophobic patch on YdbL depicted in (**i**) becomes obscured upon YnbE binding as shown in the top panel although the C-terminal patch remains exposed. **b.** The degradation assay reveals YdbL is degraded by DegP. 5 µM of purified YdbL was incubated with 5µM purified DegP at 37 °C over the course of 6 hours, with samples taken at 0 and 6 hour time-points before loading in SDS buffer to quench the reaction. A control sample using catalytically dead mutant DegP S210A was also used to indicate DegP-specific degradation of the target. **c.** In the degradation assay, YdbL is protected from degradation by DegP in the presence of the YnbE peptide. Both (**b**) and (**c**) gels are representative of biological repeats, N = 3.

We tested this hypothesis *in vitro* by incubating an equimolar ratio of purified YdbL and DegP together at 37 ℃ for 6 hours, using both wild-type DegP, and its catalytically inactive mutant S210A as a negative control (37). We observed complete degradation of YdbL in a DegP-specific manner (Fig. 4(b)). To ensure that YdbL was not unstable or aggregated during the assay, we performed a series of biophysical validation tests. SEC of YdbL before and after a 6-hour incubation at 37 ℃ revealed a distinctive lack of aggregation indicating that YdbL itself was not misfolding or aggregating under these assay conditions (SI Fig. 10(a)). Additionally, the melting temperature of YdbL was identified as 53.4 ℃ by circular dichroism, indicative of high thermal stability (SI Fig. 10(b)). These data therefore pointed towards a model where folded YdbL is degraded by DegP, perhaps via recognition of the aforementioned hydrophobic regions.

### YnbE protects YdbL from degradation by DegP *in vitro*

We next tested the effect of YnbE on YdbL degradation by DegP *in vitro*. Following incubation with the YnbE peptide, we observed drastically reduced degradation of YdbL, suggesting the peptide protected YdbL from degradation by DegP (Fig. 4(c)). Further analysis of the hydrophobic surface of the YdbL-YnbE peptide complex revealed that one hydrophobic patch of YdbL at the YnbE-binding interface was covered completely but the C-terminal hydrophobic region remained exposed (Fig. 4(a(ii))). To understand whether peptide binding may have a stabilizing effect on YdbL to inhibit degradation, we performed a thermal melt on the complex, identifying a melting temperature of 65.6 ℃, which is higher than that of the YdbL monomer (SI Fig. 10(b)). This stabilizing effect of YnbE when complexed with YdbL may obscure YdbL complexes from proteolysis by DegP and thus facilitate the clearance of apo-YdbL which could contribute to aberrant sequestration of YnbE. This would also avoid the cell degrading YdbL complexes required for YdbH-YnbE function. Together, these data suggest cells may respond rapidly to relative changes in protein levels by DegP recognizing increasing levels of YdbL that would harm YdbH-YnbE function. We could not test this model under physiologically relevant conditions since we cannot detect cellular levels of YdbL unless it is produced to high levels in engineered strains in a T7 polymerase-dependent manner.

## Discussion

Current models propose that phospholipid transport between the IM and the OM is mediated by the bridge-forming AsmA-like proteins YhdP, TamB and YdbH (7–9, 11, 15, 18, 44, 45). YdbH interacts with YnbE to form a functional intermembrane bridge (19, 20), and YdbL has been proposed to affect the structure and function of the YdbH-YnbE complex (19). Here, we show that YdbL specifically controls YdbH-YnbE complex formation by directly interacting with YnbE and likely preventing its uncontrolled multimerization. Our biochemical data demonstrating a direct interaction between YdbL and YnbE further support the chaperone-like function of YdbL and explain the dominant-negative effect of YdbL overproduction, which causes lethality in the absence of YhdP and TamB. As the YdbH-YnbE complex is essential for the growth of the Δ*yhdP* Δ*tamB* mutant, increased levels of YdbL relative to YnbE could titrate YnbE and inhibit its association with YdbH. Accordingly, YdbL-dependent lethality can be overcome by increasing YdbH-YnbE levels *in trans*.

Our structural and functional characterization of YdbL reveals the importance of conserved residues at the YnbE-YdbL interaction interface. The majority of YdbL residues at this interface, when changed, impaired both YdbL function and its binding to YnbE. Interestingly, our mutational analysis also identified variants at the predicted YdbH-YdbL interface that partially impaired YdbL function, suggesting that YdbL may also interact with YdbH, as predicted by MultiFold2 and the fact that YdbH affects the cellular levels of YdbL *in trans*. Since YdbL is required to fully rescue YdbH-YnbE function and may have roles beyond controlling YnbE, we propose that YdbL contributes to mediating the YdbH-YnbE interface by providing a docking mechanism for YnbE at the C-terminus of YdbH.

Given the effects of YdbL on YnbE and YdbH, the relative levels of these proteins are clearly critical for proper function. Unlike YdbH and YnbE, YdbL could not be detected in cells unless it was overexpressed in the KRX strain in a T7-dependent manner. Furthermore, we identified YdbL as a target for DegP-mediated degradation *in vitro*, even when thermostable and correctly folded. Since when complexed with YnbE, there was little to no degradation of YdbL, we hypothesize that cells may be able to fine-tune YdbL levels by selectively targeting apo-YdbL not engaged in complexes with YnbE for degradation. However, the mechanistic details underlying this process remain to be determined.

Our results provide a key step into understanding the YdbH-YnbE-YdbL complex, underpinning the direct interaction of YdbL with YnbE as crucial for modulating YdbH-YnbE function as one of the major putative phospholipid transporters between the IM and OM. We hypothesize that YdbL prevents runaway self-multimerization of YnbE via a steric hindrance model (Fig. 5). Given the relatively subtle structural changes that occur at the YdbL-YnbE interaction site, it is plausible that the globular region of YdbL appended to the C-terminal α-helix of YnbE may inhibit β-strand associations between YnbE proteins by spatially inhibiting hydrogen bonding networks. Docking of YdbL-YnbE onto YdbH would allow formation of the YdbH-YnbE complex. Fine-tuning relative levels of these proteins is critical for proper YdbH-YnbE function and our finding that YdbL is targeted for degradation by periplasmic protease DegP provides a possible mechanism for rapid cellular control of YdbL.

**Figure 5.**
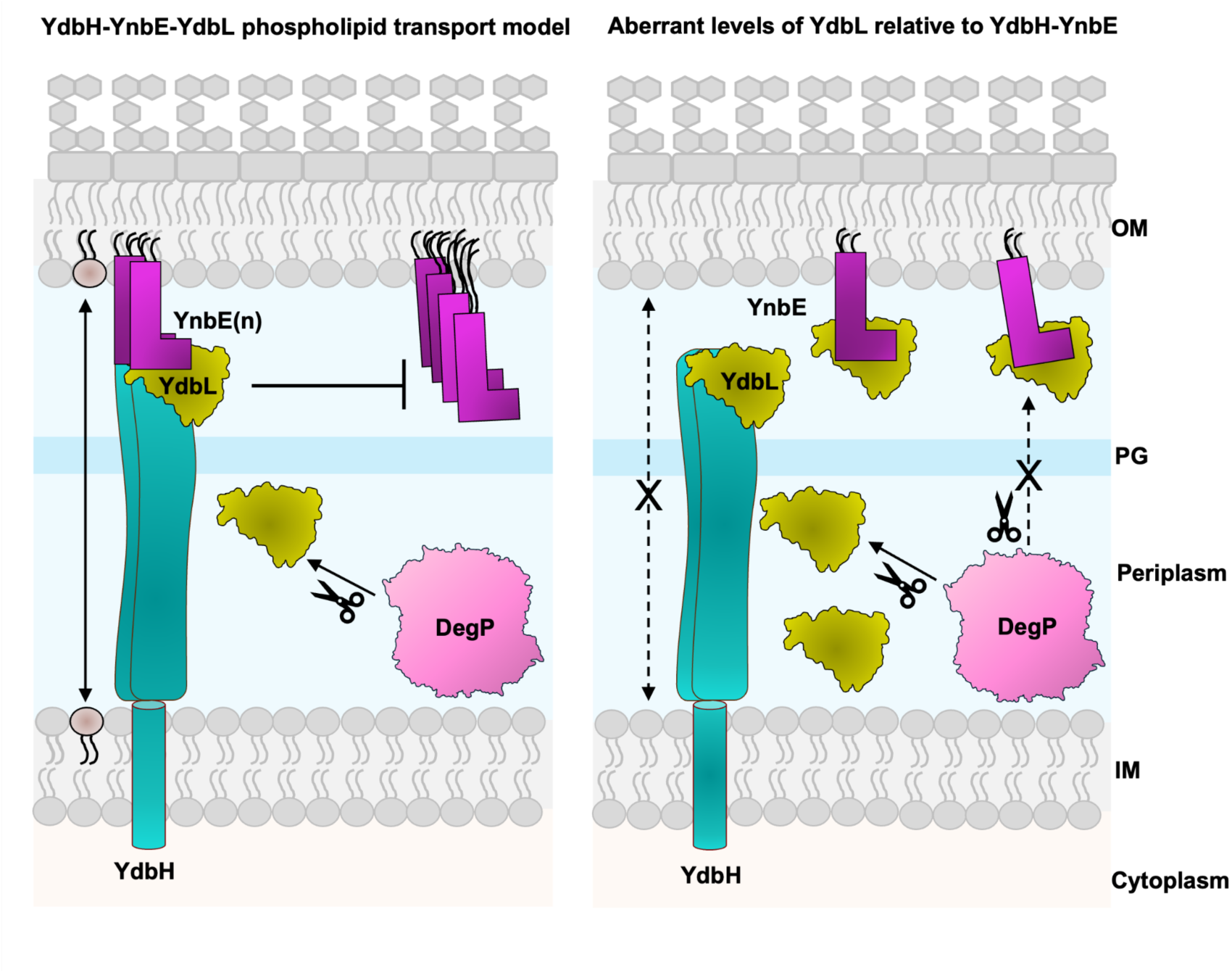
Model for function of the YdbH-YnbE-YdbL complex. The left panel shows the expected phospholipid transport model for YdbH-YnbE-YdbL. Here YdbL aids complex formation of the YdbH-YnbE intermembrane bridge and prevents runaway multimerisation of YnbE by steric hindrance which can affect efficiency of YdbH-YnbE complex formation. We speculate that cellular control systems such as DegP degradation can selectively remove apo-YdbL not engaged in complexes with YnbE. The right panel shows the model for aberrant levels of YdbL affecting YdbH-YnbE phospholipid transport function. Here YdbL binds to YdbH but also forms complexes with YnbE that are titrated from YdbH such that an intermembrane bridge for continuous transport no longer forms. This model explains why increasing the levels of both YdbH and YnbE are needed to overcome the negative effects of YdbL.

## Materials and Methods

### Bacterial strains and culture conditions

Strains used in this study were derived from the wild-type *E. coli* K-12 strain MG1655 and are listed in Table S1. Gene deletions from the Keio collection (46) were introduced into target strains via generalized P1_vir_ transduction, with kanamycin resistance serving as the selection marker. When necessary, the kanamycin-resistance cassette was removed using Flp recombinase (47). Unless indicated otherwise, bacterial cultures were grown in lysogeny broth (LB Lennox; Fisher Scientific) at 37 °C with shaking, and growth was monitored by measuring optical density at 600 nm (OD_600_). Solid media consisted of LB agar plates (15 g/L agar; BD Difco) or MacConkey agar (BD Difco). Where appropriate, antibiotics and supplements were added at the following concentrations: ampicillin (125 μg/mL), bacitracin (100 μg/mL), chloramphenicol (20 μg/mL), kanamycin (30 μg/mL), spectinomycin (100 μg/mL), L-rhamnose (0.2% w/v) and L-arabinose (0.02% or 0.2% w/v) and *p*-benzoylphenylalanine (pBPA, 0.48 mM in 1N NaOH).

### Site-directed mutagenesis (SDM)

Primers and details of plasmids generated using SDM are listed in Table (S2 and S3), respectively. Phusion High-Fidelity DNA Polymerase (New England Biolabs; NEB) was used to perform SDM PCR. The resulting PCR products were digested with DpnI to remove the template and purified using the GeneJET PCR purification kit (Thermo Fisher Scientific). The purified DpnI-digested PCR products were electroporated into DH5α (Table S1). Transformants carrying pBAD33-SpecR- and pET23/42-derived plasmids were selected on LB agar with 100 µg/mL spectinomycin and 125 µg/mL ampicillin, respectively. For deletion and insertion SDM, T4 DNA ligase (NEB) was used to ligate the DpnI-digested PCR products after the treatment with T4 polynucleotide kinase (NEB). Plasmids were isolated using GeneJET Plasmid miniprep kit (Thermo Fisher Scientific) and confirmed by Sanger sequencing.

### Construction of pBAD33-SpecR-RBS-YdbH/YnbE/YdbL plasmid

To construct pBAD33-Native-YdbH/YnbE/YdbL plasmid, the chromosomal locus of ydbH operon (*ydbH-ynbE-ydbL*) including 20 bases upstream containing ribosome binding site (RBS), was amplified from wild-type genomic DNA using primers 5YdbH-YdbL_SacI and 3YdbH-YdbL_HindIII (Table S2). The resulting PCR product and pBAD33 plasmid(21) were digested with restriction endonuclease SacI (NEB) and HindIII-HF (NEB). After purification, the fragments were ligated together using T4 DNA ligase (NEB) following the manufacturer’s protocol. The ligation mixture was repurified and electroporated into DH5α, then selected on the LB plate containing chloramphenicol. Plasmids were isolated and confirmed by Sanger sequencing.

The chloramphenicol resistance cassette on both the pBAD33 vector and pBAD33-YdbH/YnbE/YdbL was replaced with spectinomycin cassette using recombineering (48). Briefly, the spectinomycin cassette from pCL-His6-LptB (49) was amplified using primers 5pBAD33specP1 and 3pBAD33specP2 (Table S2). The purified DpnI-digested PCR product was electroporated into recombineering strain DY378 (48) carrying either pBAD33 vector or pBAD33-YdbH/YnbE/YdbL. Spectinomycin-resistant recombinants carrying pBAD33SpecR or pBAD33SpecR-YdbH/YnbE/YdbL were selected at 30 °C. Plasmids were confirmed by Sanger sequencing.

To generate pBAD33SpecR-RBS-YdbH/YnbE/YdbL, a strong RBS sequence (TTTAAGAAGGAG ATATACAT; RBS underlined) was inserted upstream of *ydbH* start codon to replace the native RBS present on pBAD33-SpecR-YdbH/YnbE/YdbL.

### Construction of pET23/42 GST-His_2_-YdbH/YnbE/RBS-FLAG-YdbL plasmid

The pET23/42GST-His_2_-YdbH/YnbE/YdbL plasmid (19) was modified by inserting a FLAG tag-encoding sequence immediately downstream of the sequence encoding the YdbL signal sequence, generating the pET23/42GST-His_2_-YdbH/YnbE/FLAG-YdbL plasmid. A strong RBS sequence was then introduced upstream of the *ydbL* start codon by replacing the DNA sequence <u>TGA</u>GGTGATG (stop codon underlined) which contains the *ynbE* stop codon and seven nucleotides upstream of the *ydbL* start codon, with <u>TAA</u>TTTAAG<u>AAGGAG</u>ATATACAT (stop codon and RBS are underlined) to generate pET23/42GST-His_2_-YdbH/YnbE/RBS-FLAG-YdbL.

### Construction and *In Vivo* Photocrosslinking of strains

For *in vivo* photocross-linking experiments, we used a previously described method (19, 22) with the following modification. To construct plasmid-encoded *ynbE* amber alleles, SDM was used to convert specific codons into amber stop codons (5′-TAG). These plasmids were then used to transform NR7359 (19) carrying the pSUP-BpaRS-6TRN (50) (Table S1). To generate the required cross-linking strains, either pBAD33-SpecR or pBAD33-SpecR-RBS-YdbL was introduced, and transformants were selected on LB agar supplemented with ampicillin, chloramphenicol, and spectinomycin. Overnight cultures of these cross-linking strains were diluted into 5 mL of LB supplemented with 125 µg/mL ampicillin, 20 µg/mL chloramphenicol, 100 µg/mL spectinomycin, 0.48 mM pBPA and 0.2% (w/v) arabinose, and grown at 37 ⁰C until reaching an OD_600_ ∼ 0.8 to 1.0. In a 24-well flat-bottom cell culture plate (Costar; Corning Inc.), half of each culture was exposed to UV light (365 nm; Spectroline E series) for 30 minutes at room temperature, while the remaining half was left untreated. Following treatment, cell pellets were collected by centrifugation at 16,873 × g for 1 minute at room temperature and resuspended in 100 μL of 1× Laemmli sample buffer containing 12.5 units of Benzonase (Novagen). Equal volumes of unboiled samples were separated on 15% SDS-polyacrylamide gels and analyzed by immunoblotting as described below.

### Efficiency of plating assay

We used a previously described protocol (8) using overnight cultures that were serially diluted in LB with the appropriate antibiotics at a ratio of 1:10 (v/v). Dilutions were spotted on plates using a 48-pin replicator and plates were incubated at 37°C overnight before imaging.

### Functional characterization of pET23/42 FLAG-YdbL variants

Functionality of plasmid-encoded flag-*ydbL* mutant alleles was tested based on the fact that plasmid-encoded *ydbL* causes lethality in the Δ*tamB* Δ*yhdP* double mutant (NR5161) (Table S1), but not in the wild type or the Δ*tamB* or Δ*yhdP* single mutants (19). Strain NR5161 was electroporated with pET23/42FLAG-YdbL-derived plasmids, and functionality was evaluated by checking colony formation on LB agar supplemented with ampicillin. Plasmids encoding non-functional flag-*ydbL* alleles produced stable colonies, whereas those encoding functional flag-*ydbL* alleles produced none. The non-functional flag-*ydbL* alleles were further examined for growth defects by performing a 24-hr growth curve in comparison with NR5161 carrying the pET23/42 vector.

### Construction and preparation of KRX strains for detection of FLAG-YdbL

To check the protein levels of FLAG-YdbL variants, we generated strains by electroporating the pET23/42FLAG-YdbL-derived plasmids into KRX strain (Promega). Additionally, we constructed the KRX Δ*ydbH-ydbL*::kan (NR7980) strain through recombineering using KRX carrying the pKD46 plasmid (47) (NR5530) (Table S1). Briefly, the Δ*ydbH-ydbL*::kan region from MG1655 Δ*ydbH-ydbL*::kan (NR6436) (Table S1) was amplified using colony PCR with primer sets listed in Table S2 and described in Table S3. The PCR product was purified and electroporated into recombineering strain NR5530. Kanamycin-resistant recombinants were selected on LB agar supplemented with kanamycin at 30 °C to obtain KRX Δ*ydbH-ydbL*::kan (NR7980), which was confirmed by PCR.

To prepare whole-cell lysate samples, overnight culture of KRX strains carrying pET23/42FLAG-YdbL-derived plasmids were diluted into 5 mL of LB with appropriate antibiotics and grown at 37 ⁰C until mid-log phase (OD_600_ ∼ 0.6 - 0.7). Cultures were then induced with 0.2% (w/v) L-rhamnose and incubated for 3 h at 37 °C. Samples were normalized by pipetting a culture volume (in μL) equivalent to 800 divided by the OD_600_ of the culture. Cells were pelleted by centrifugation at 16,873 x g for 1 min at room temperature and resuspended in 50 μL of 1× Laemmli sample buffer, achieving a final concentration of cells equivalent to OD_600_ ∼ 16. The whole-cell lysate samples were boiled for 10 min and loaded onto 15% SDS-polyacrylamide gels for electrophoresis, followed by immunoblotting as described below.

### Immunoblot analysis of YnbE-FLAG and FLAG-YdbL

Immunodetection of YnbE-FLAG and FLAG-YdbL was done using whole-cell lysate samples prepared as described above. After gel electrophoresis, proteins were then transferred to polyvinylidene difluoride (PVDF) membranes using a semidry transfer cell (Bio-Rad) at 10 V for 2 hrs. Membranes were blocked in TBST milk [1X Tris-buffered saline (pH 7.6), 0.1% Tween-20 with 5% (w/v) nonfat dry milk] on a shaker for 30 min at room temperature and probed with Anti-FLAG M2 (1:10,000; Millipore Sigma) and anti-mouse horseradish peroxidase (1:10,000; GE Healthcare). Signal was developed using Clarity Max Western ECL substrate (Bio-Rad) and detected with a ChemiDoc XRS+ system (Bio-Rad).

### Purification of YdbL

DNA encoding mature YdbL (residues 22-108) was amplified by PCR from *E. coli* BW25113 (46) and cloned into a custom pET vector (pBEL1092) by Gibson assembly to obtain YdbL variants with an N-terminal 6xHistag followed by a TEV site with (pGLI190) and without the native signal sequence (pGLI171) Table S4. Primers and plasmid construction details can be found in Tables S2 and S3. SignalP 6.0 (51) was used to identify the native signal sequence from residues 1 to 21. Resulting plasmids were transformed into T7 Express cells (NEB) for expression and purification.

For expression, overnight cultures in LB (Miller) containing 1% glucose and 100 µg/mL carbenicillin were diluted 1:50 and grown at 37 °C, 180 rpm to an OD_600_ of 0.8. Cultures were induced overnight at 15 °C, 180 rpm with 1 mM IPTG. Cells were harvested by centrifugation and resuspended in 50 mM Tris pH 8, 300 mM NaCl, 10 mM imidazole, 10 % glycerol with an EDTA-free cOmplete Protease Inhibitor Cocktail tablet (Roche). Cells were lysed by 3-4 passes through an Emulsiflex C3 cell disruptor (Avestin). Lysate was clarified by centrifugation at 30,000 g for 30 min at 4 °C. Supernatant was incubated with Ni Sepharose 6 Fast Flow for at least 1 h at 4 °C. Resin was washed in 50 mM Tris pH 8, 300 mM NaCl, 30 mM imidazole before eluting in 50 mM Tris pH 8, 300 mM NaCl, 250 mM imidazole at 4 °C. For TEV cleavage, YdbL was incubated with TEV (1:10 TEV:protein) for 1 h at room temperature before dialysis overnight at 4 °C in 50 mM Tris pH 8, 300 mM NaCl using SnakeSkin 3.5 kDa MWCO dialysis tubing. A second Ni Sepharose 6 Fast Flow step was performed to isolate the cleaved protein. Finally, protein-fractions were pooled and concentrated using a 10 kDa MWCO centrifugal filter (Amicon) and polished by size exclusion chromatography (SEC) on a Superdex 75 Increase 10/300 GL (S75 10/300I) column equilibrated with 50 mM Tris pH 8, 150 mM NaCl. Samples were analyzed by SDS-PAGE Coomassie staining throughout purification.

### Crystallization and structure determination of YdbL

The Versatile Macromolecular Xtallography in-situ (VMXi) beamline (23, 24), Diamond Light Source was used to crystallize and collect room temperature X-ray diffraction data for YdbL. YdbL was concentrated between 15 - 45 mg/mL for crystallization experiments using a 10 kDa MWCO centrifugal filter (Amicon). Vapor diffusion experiments were set-up with a Mosquito® crystal robot using MiTeGen In-Situ-1™ 96-well plates at a range of protein concentrations, crystallization screens and protein:reservoir ratios, with 100 nL or 50 nL reservoir added to 100 nL protein. Samples were stored at 20 °C and imaged using a Rock Imager (Formulatrix). The best diffracting crystals were grown with 100 nL 26 mg/mL YdbL (pGLI171 construct) added to 50 nL reservoir composed of 0.1 M HEPES pH 7, 30 % (w/v) Jeffamine ED-200130 pH 7 from the Hampton Index screen.

Samples were handled by the robot and crystals of interest were collected *in-situ* using the VMXi-beamline with a Dectris Eiger2 × 4M detector. Data processing was triggered automatically upon data collection. Several data-processing pipelines were used to index and merge diffraction data as per other Diamond MX beamlines (52, 53). Successful DIALS via xia2 triggered the secondary xia2.multiplex pipeline (54) to sort and merge individual datasets into a consistent isomorphous dataset. The number of copies in the asymmetric unit (ASU) was estimated using Matthew’s coefficient (55, 56) in CCP4. The indexed Scaled.mtz file was imported into Phenix. The structure was phased via molecular replacement using Phaser (57) with the AlphaFold-predicted structure (P76076, residues 22-108 with signal-sequence truncated). The resulting model was refined using Phenix.refine (58), with manual inspection and adjustment in Coot (59) between refinements. The final structure consisted of one copy of YdbL in the ASU with a final R_work_/R_free_ of 0.1958/0.2404.

### Structure prediction, analysis and interpretation

A full-length structural prediction for YdbL was downloaded from the AlphaFold protein structure database (https://alphafold.ebi.ac.uk/) (60, 61) using its UniProt accession code P76076. The MultiFold2 server (32, 33) (https://www.reading.ac.uk/bioinf/MultiFOLD/) was used to simulate proteins of interest together using protein sequences available from UniProt with signal sequences removed if applicable (YdbH: P52645, YnbE: P64448 and YdbL:P76076). The ModFoldDock2 server (32, 62) (https://www.reading.ac.uk/bioinf/ModFOLD/) was used to assess the quality of the predicted interaction interface by MultiFold2. UCSF ChimeraX (v1.7.1) was used for all structural visualization and used to analyze molecular lipophilic potential (mlp command), coulombic electrostatic potential (coulombic command), structural alignment (matchmaker mm command) to get rmsd between atom pairs, contact sites (by buried solvent accessible area more than or equal to 15 Å^2^) and hydrogen bonding. For the crystal structure of YdbL, the ConSurf server (35, 63–67) (https://consurf.tau.ac.il/consurf_index.php) was used to determine the evolutionary conservation of residues using phylogenetic relationships between homologous sequences. The Dali server (68–70) (http://ekhidna2.biocenter.helsinki.fi/dali/) was used to evaluate structural homology between other 3D structures and YdbL. PDBePISA (26) (https://www.ebi.ac.uk/pdbe/pisa/pistart.html) was used to evaluate crystal-contact interfaces.

### SEC-SAXS

Purified YdbL expressed from pGLI171 or pGLI190 constructs were concentrated between 4.8 - 9.9 mg/mL with a final buffer composition 50 mM Tris pH 8, 150 mM NaCl, 1% Glycerol. SEC-SAXS was conducted on the B21 beamline, Diamond Light Source. A 45 µL injection volume was used for SEC-SAXS data collection on a S75 3.2/300I precision column, with BSA run as control to check the column was working effectively. Further experimental details can be found in (71). Data were reduced and analyzed using ScatterIV (https://bl1231.als.lbl.gov/scatter/). Following data reduction, experimental .dat files were inputted into the Fast open-source X-ray Scattering (FoXS) server (https://modbase.compbio.ucsf.edu/foxs/) (72, 73) to compare their fit with the YdbL crystal structure.

### Solution-state Nuclear Magnetic Resonance (NMR) Spectroscopy

Single- (^15^N-) and double- (^13^C-^15^N-) labelled YdbL were prepared by condensation induction. In brief, overnight cultures of T7 Express cells transformed with pGLI171 grown in LB with 100 ng/µL carbenicillin and 1% glucose were diluted 1:15 in LB with carbenicillin before growing to OD_600_ between 0.800 to 0.850. Cells were harvested at 4,000 g for 15 min at room temperature before removing supernatant and resuspending gently in 1x M9 minimal media to wash cells of excess LB media using a second 4,000 g, 15-min centrifugation at room temperature. After removing supernatant, cells were resuspended gently in 2x M9 minimal media supplemented with 1 g/L ^15^N-NH_4_Cl and 2.5 g/L D-glucose (for ^15^N-YdbL) or 2 g/L ^13^C-D-glucose (for ^13^C-^15^N-YdbL) as the sole nitrogen and carbon sources. Cells were concentrated from 2 L to 1 L for isotope labelling. Cells were incubated statically at room temperature for 30 min before inducing with 2 mM IPTG overnight at 18 ℃, 180 rpm. Labelled YdbL was harvested and purified as described previously.

The C-terminal YnbE peptide (H-EHEIIIKADKDVEELLETRSDLF-OH) was synthesized to >98% purity by GL Biochem. The lyophilized peptide was reconstituted in 10 mM Na_2_H_2_PO_4_, 20 mM NaCl, pH 7.4 buffer used for NMR data collection which required small pH correction with NaOH.

Final ^15^N- or ^13^C-^15^N-labelled YdbL samples were buffer exchanged via SEC for NMR experiments on a Superdex 75 10/300I column equilibrated in 10 mM Na_2_H_2_PO_4_, 20 mM NaCl, pH 7.4 and concentrated to 180 µM for ^15^N-YdbL and 60 µM for ^13^C-^15^N-YdbL. Peptide was added to be equimolar to the concentration of protein. Samples were supplemented with 5% D_2_O and added to 5 mm Shigemi tubes to a final volume of 350 µL. All NMR experiments were performed at 20 ℃ using the 750 MHz and 950 MHz spectrometers equipped with Oxford Instruments Company magnets, Bruker Avance III HD consoles and 5 mm TCI CryoProbes according to standard triple-resonance protocols (74).

Resonance assignments for apo-YdbL were obtained using both ^15^N- and ^13^C-^15^N-labelled protein with 2D experiments including ^1^H-^15^N HSQC, ^1^H-^13^C HSQC and ^1^H-^15^N BEST-TROSY and3D experiments including ^15^N-edited NOESY-HSQC, ^15^N-edited TOCSY-HSQC, (H)CC(CO)NH and BEST-TROSY versions of HNCA, HNCACB and HNCO (75, 76). Experiments with YnbE peptide-bound YdbL were conducted as per apo-YdbL with the exception of 3D HNCACB which was not collected and 3D HCCH-TOCSY which was collected. All 3D NMR data were collected with 25% non-uniform sampling in the two indirect dimensions using standard Bruker sampling schedules. 2D NMR were processed using NMRPipe (77) and 3D NUS data were processed with the hmsIST software (78) and NMRPipe.

NMR spectra were analyzed and assignments recorded using CcpNmr Analysis v2.5.2 (79). Sodium 3-(trimethylsilyl)-1-propanesulfonate (DSS) was used to reference ^1^H and ^13^C chemical shifts whilst ^15^N chemical shifts were referenced indirectly. Details of specific experiments and sample conditions can be found in the BMRB deposition files.

### Expression test and purification of YdbL variants

YdbL variants were encoded from pGLI190 as per Tables S3 and S4. Mutant proteins were produced purified as described previously for wild-type YdbL.

To detect levels of His-tagged YdbL mutants after expression, T7 express cells transformed with each mutant plasmid combination were diluted 1:50 in LB supplemented with Carbenicillin (100 µg/mL) and grown at 37 ℃, 180 rpm shaking to an OD_600_ of 0.8-0.9. Cells were induced with 1 mM IPTG and expressed overnight at 15 ℃, 180 rpm. Cells were measured for OD_600_ and an equivalent of 1 mL cells at OD_600_ = 1.0 were pelleted and resuspended in 4x SDS-loading buffer with PBS. The mixtures were then heated at 95℃ for 10 minutes before spinning by centrifugation. 10 µL of supernatant were separated on a Novex 4-12% Tris-Glycine gel in 1x MES by SDS-PAGE and transferred to a nitrocellulose membrane. Membranes were blocked in PBST containing 5% (w/v) skimmed milk for 1 hour. Membranes were then incubated with rabbit polyclonal anti-RecA (Abcam ab63797) at a dilution of 1:3000 or mouse monoclonal anti-His (Abcam ab18184) at a dilution of 1:1000 in PBST + 5% (w/v) skimmed milk overnight at 4 ℃. The membranes were then washed three times with PBST and were incubated with IRDye 800CW goat anti-rabbit IgG polyclonal antibody (LICOR) at a dilution of 1:20000 or IRDye 800CW goat anti-mouse IgG polyclonal antibody (LICOR) at a dilution of 1:20000 in PBST + 5% (w/v) skimmed milk for 1 hour room temperature. The membranes were then washed five times with PBST before a final rinse with PBS to remove excess Tween-20. Blots were imaged using the Odyssey CLx Imager (LICOR).

### Native mass spectrometry

Before MS analysis, YdbL and mutants were buffer exchanged into 200 mM ammonium acetate pH 8 using Biospin-6 (BioRad) columns. Reconstituted peptide in water was diluted to a final concentration of 100 µM in 200 mM ammonium acetate pH 8. Peptide was added to protein in an equimolar ratio such that final peptide and protein concentrations were both 3.5 µM.

Samples were introduced directly into the mass spectrometer using gold-coated capillary needles (prepared in-house). Data were collected on a Q-Exactive UHMR mass spectrometer (Thermo Fisher Scientific). The instrument parameters were as follows: capillary voltage 0.8 kV, instrument operation in negative mode, S-lens RF 100%, quadrupole selection 600 m/z to 10,000 m/z, neither collision activation in the HCD cell nor in-source dissociation applied, noise level 3, trapping gas setting 7.5, temperature 200 ℃, orbitrap resolution 12,500.

Data were analyzed using Xcalibur 4.1 Qual Browser (Thermo Scientific) and UniDec (Marty et al., 2015) software packages. The ratio of peptide-bound vs apo YdbL was determined by numerical integration. Measurements were repeated in triplicates, and the errors are reported as the standard deviation between measurements. All calculated vs expected masses can be found in Table S5.

### Expression and purification of DegP

Wild-type DegP and S210A DegP expression constructs were kindly provided by Tim Clausen (Research Institute of Molecular Pathology, Vienna, Austria).

Constructs were transformed into T7 express. For expression, overnight cultures containing 1% glucose and 100 ng/uL Carbenicillin were diluted 1:50 and grown at 37 °C, 180 rpm to an OD_600_ of 0.9. Cultures were induced overnight 15 °C, 180 with 1 mM IPTG. Cells were harvested by centrifugation and resuspended in 50 mM Tris pH 8, 300 mM NaCl, 10 mM Imidazole, 10 % Glycerol. Cells were lysed by 3-4 passes through an Emulsiflex C3 cell disruptor (Avestin). Lysate was clarified by centrifugation at 30,000 g for 30 minutes. Supernatant was incubated with Ni Sepharose 6 Fast Flow at 4 °C overnight. Resin was washed in 50 mM Tris pH 8, 300 mM NaCl, 40 mM Imidazole before eluting in 50 mM Tris pH 8, 300 mM NaCl, 250 mM Imidazole. Finally, fractions of interest were pooled and concentrated using a 30 kDa MWCO centrifugal filter (Amicon) and polished by size exclusion chromatography (SEC) on a Superdex 200 Increase 10/300 GL (S200 10/300I) column equilibrated in 20 mM Tris pH 8, 150 mM NaCl. Samples were analyzed by SDS-PAGE Coomassie staining throughout purification.

### DegP degradation assay

5 µM of purified YdbL and YnbE peptide were incubated with equimolar DegP in 20 mM Tris pH 8, 150 mM NaCl at 37 ℃ for 6 hours, with samples taken at 0- and 6-hour timepoints before adding into SDS loading buffer to quench the reaction. A control sample with mutant S210A DegP was used to indicate DegP-specific degradation of the target.

### Biophysical analysis of YdbL by SEC and circular dichroism

To monitor aggregation propensity during the timeframe of the DegP degradation assay, 10 µM YdbL was incubated in 20 mM Tris pH 8, 150 mM NaCl at 37 ℃ for 6 hours before conducting SEC on an S75 10/300I column equilibrated in 20 mM Tris pH 8, 150 mM NaCl. A control sample at 0 hours was also loaded for comparison. Peak fractions were monitored for loss of sample and formation of an aggregate peak in the void volume.

For circular dichroism, purified YdbL was buffer exchanged by SEC using an S75 10/300I column into 10 mM NaH_2_PO_4_ (pH 7.4), 20 mM NaCl. Samples were diluted to a final concentration of 0.1 mg/mL. Circular dichroism spectra were obtained using a Jasco J-815 Spectropolarimeter over a wavelength range of 260-190 nm, a digital integration time of 1 s, and a 1 mm bandwidth. HT voltage signal was collected in parallel. Protein melting temperature was determined by collecting 200 - 250 nm scans as a function of temperature from 22 to 90 ℃ in 2 ℃ intervals. For peptide-bound YdbL experiments, reconstituted peptide was added in a 2:1 excess of YdbL.

## Author Contributions

Conceptualization: C.D., S.K., N.R., G.L.I.; Phenotypic assays: S.K., S.C.; *In vivo* photocrosslinking experiments: S.K.; Strain construction: S.K., S.C.; Plasmid cloning: S.K., S.C., C.D., B.F.C.; X-ray crystallography: C.D., B.F.C.; NMR experiments: C.D., C.R.; Native MS experiment: C.D., J.R.B.; DegP degradation assays: C.D., E.K.T., E.J., T.L.H.; Protein purification: C.D., E.K.T., E.J.; Biophysics: C.D., E.J.; Interpretation of results: all authors; Supervision: N.R., G.L.I.; Writing - original draft: C.D., S.K., N.R., G.L.I; Writing - review & editing: all authors.

## Acknowledgments

Firstly, we would like to thank our funding sources: 228310/Z/23/Z (Wellcome Trust Grant awarded to C.D.), R35GM153349 (National Institute of General Medical Sciences Grant awarded to N.R.), European Research Council (ERC) under the European Union’s Horizon Europe research and innovation programme (grant agreement No. 101162143 awarded to G.L.I.), MR/W016672/1 (Medical Research Council career development award awarded to G.L.I), URF\R1\211567 (Royal Society grant awarded to J.R.B.), 317713/Z/24/Z (Wellcome Trust Grant awarded to T.L.H), 218514/Z/19/Z (Wellcome Trust Grant awarded to E.K.T.). Additionally, the 950 MHz NMR spectrometer was upgraded with funding from the University of Oxford Wellcome Institutional Strategic Support Fund, the John Fell Fund, and the Edward Penley Abraham Cephalosporin Fund, and from the Engineering and Physical Sciences Research Council (EP/R029849/1). We thank Diamond Light Source for beamtime (proposal mx31353) and the staff of beamlines VMXi (in particular Halina Mikolajek and Michael Hough) for assistance with crystal generation, testing and data collection, and B21 (in particular Nathan Cowieson) for assistance with data collection and processing. We thank David Staunton (Department of Biochemistry, University of Oxford) for assistance with CD experiments. We would like to thank Jacob Hong (Department of Microbiology, The Ohio State University) for his helpful assistance in the construction of a few *flag-ydbL* alleles. Expression plasmids for wild-type and S210A DegP, were kindly provided by Tim Clausen (Research Institute of Molecular Pathology, Vienna, Austria). We would also like to thank Matthew Hankins and Qiaoyu Tian (Sir William Dunn School of Pathology, University of Oxford) for critical reading of the manuscript.

## Supplemental Information

**Fig. S1.**
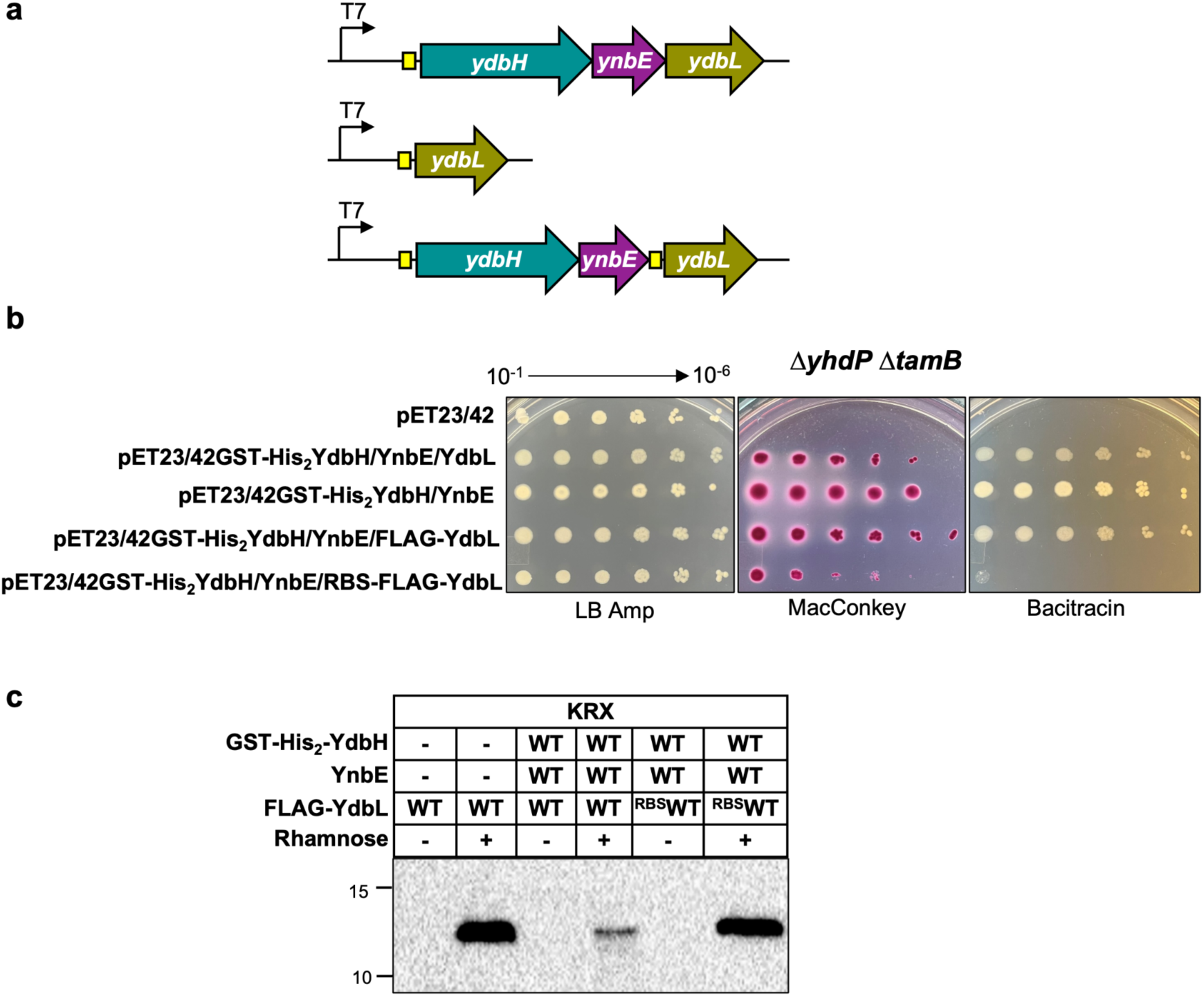
In the absence of YhdP and TamB, YdbL is required for suppression of mucoidy by plasmid-encoded YdbH-YnbE and causes lethality in excess. a. The genetic arrangement of T7 promoter, the ribosome binding site (RBS; yellow box), and *ydbH* (teal), *ynbE* (plum) and *ydbL* (gold) encoded on pET23/42-derived plasmids. b. The effect of higher levels of YdbL on the growth of Δ*yhdP* Δ*tamB* double mutant was tested by spotting serially diluted cultures of Δ*yhdP* Δ*tamB* mutant strains carrying pET23/42-derived plasmids on LB agar containing ampicillin, MacConkey agar, and LB agar containing bacitracin. Increasing production of YdbL through introduction of an RBS upstream of the ydbL start codon on a pET23/42-derived plasmid impaired growth of the Δ*yhdP* Δ*tamB* double mutant on MacConkey agar and LB agar with bacitracin. c. Immunoblot analysis showed that introducing an RBS upstream of *ydbL* on pET23/42-derived plasmids increased the cellular level of YdbL. KRX strains carrying FLAG-YdbL encoded on pET23/42-derived plasmid were grown to log phase, after which they were either left uninduced or induced with 0.2% rhamnose. They were then incubated for an additional 3 hours at 37 ⁰C before sample preparation. Whole-cell lysate samples were prepared as described in the Materials and Methods section, and proteins were separated on 15% SDS-polyacrylamide gel and probed with α-FLAG antibodies by immunoblotting. Molecular mass markers (in kDa) are shown on the left. All data are representative of three independent experiments.

**Fig. S2.**
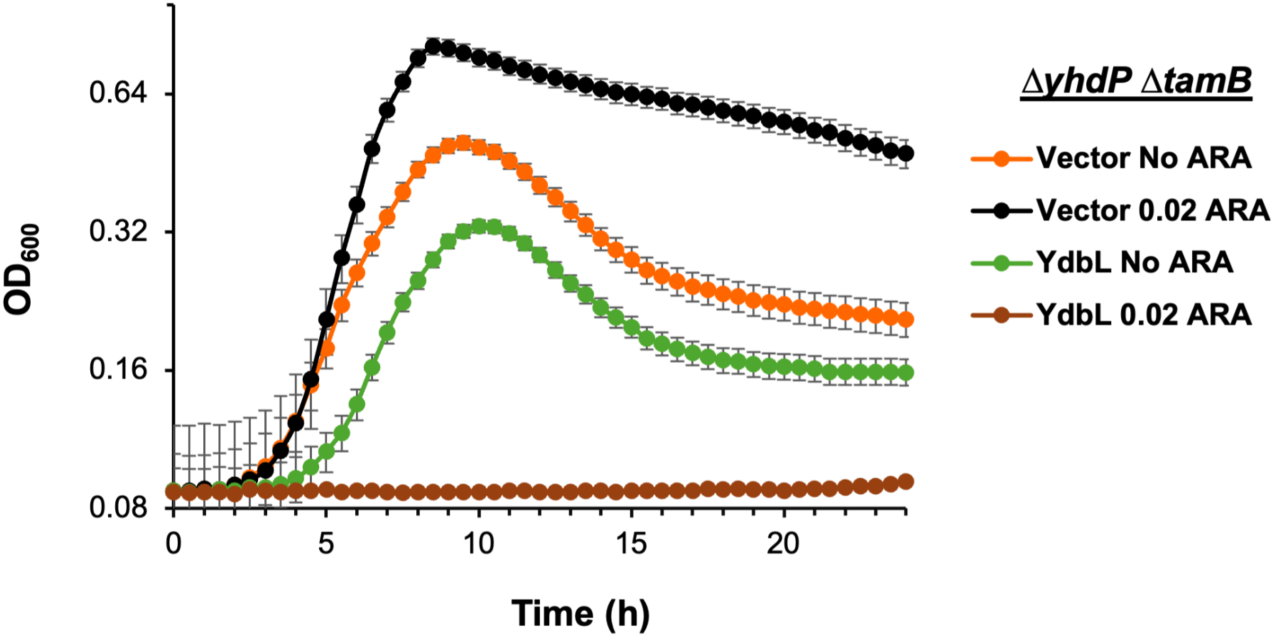
Overexpression of YdbL from pBAD33 causes lethality in the Δ*yhdP*::frt Δ*tamB*::frt mutant. Growth curves of cultures grown in LB containing spectinomycin with or without 0.02% arabinose at 37 °C. The metabolic effect of arabinose consumption allowed the Δ*yhdP* Δ*tamB* double mutant carrying the pBAD33-SpecR vector to grow better under 0.02% arabinose condition than in its absence. The Δ*yhdP* Δ*tamB* double mutant carrying pBAD33-SpecR-RBS-YdbL failed to grow in the presence of 0.02% arabinose, and the lysis of this strain compared to Δ*yhdP* Δ*tamB* double mutant carrying pBAD33-SpecR vector was faster even without arabinose induction due to leaky expression of *ydbL*. OD600 values are represented on a logarithmic scale. Data represent the average and standard deviation from three biological replicates.

**Fig. S3.**
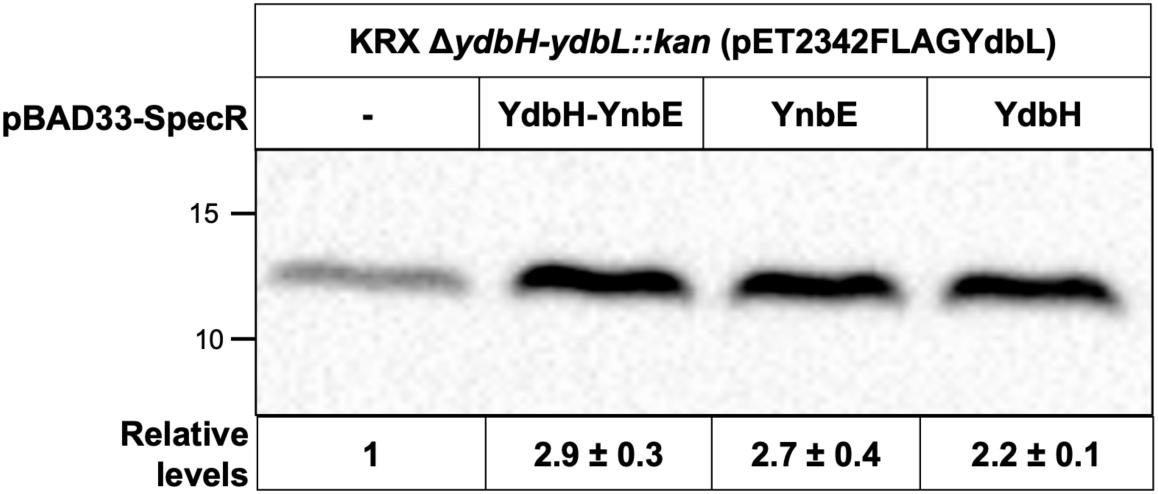
Producing YdbH and/or YnbE *in trans* increases the levels of YdbL. KRX Δ*ydbH-ydbL*::*kan* strains carrying pET23/42FLAG-YdbL and pBAD33-SpecR-derived plasmids were grown to log phase. Production of YdbH and/or YnbE was induced for three hours at 37 ⁰C with 0.2% rhamnose. Whole-cell lysate samples were prepared as described in Materials and Methods, and proteins were separated on 15% SDS-polyacrylamide gel and probed with α-FLAG antibodies by immunoblotting. When YdbH, YnbE, or both were produced from the pBAD33 plasmid, increased levels of FLAG-YdbL produced from the pET23/42 plasmid were detected. For calculating relative levels, the signal from samples derived from cells with pBAD33 vector plasmid was set to 1. Molecular mass markers (in kDa) are shown on the left. All data are representative of three independent experiments.

**Fig. S4.**
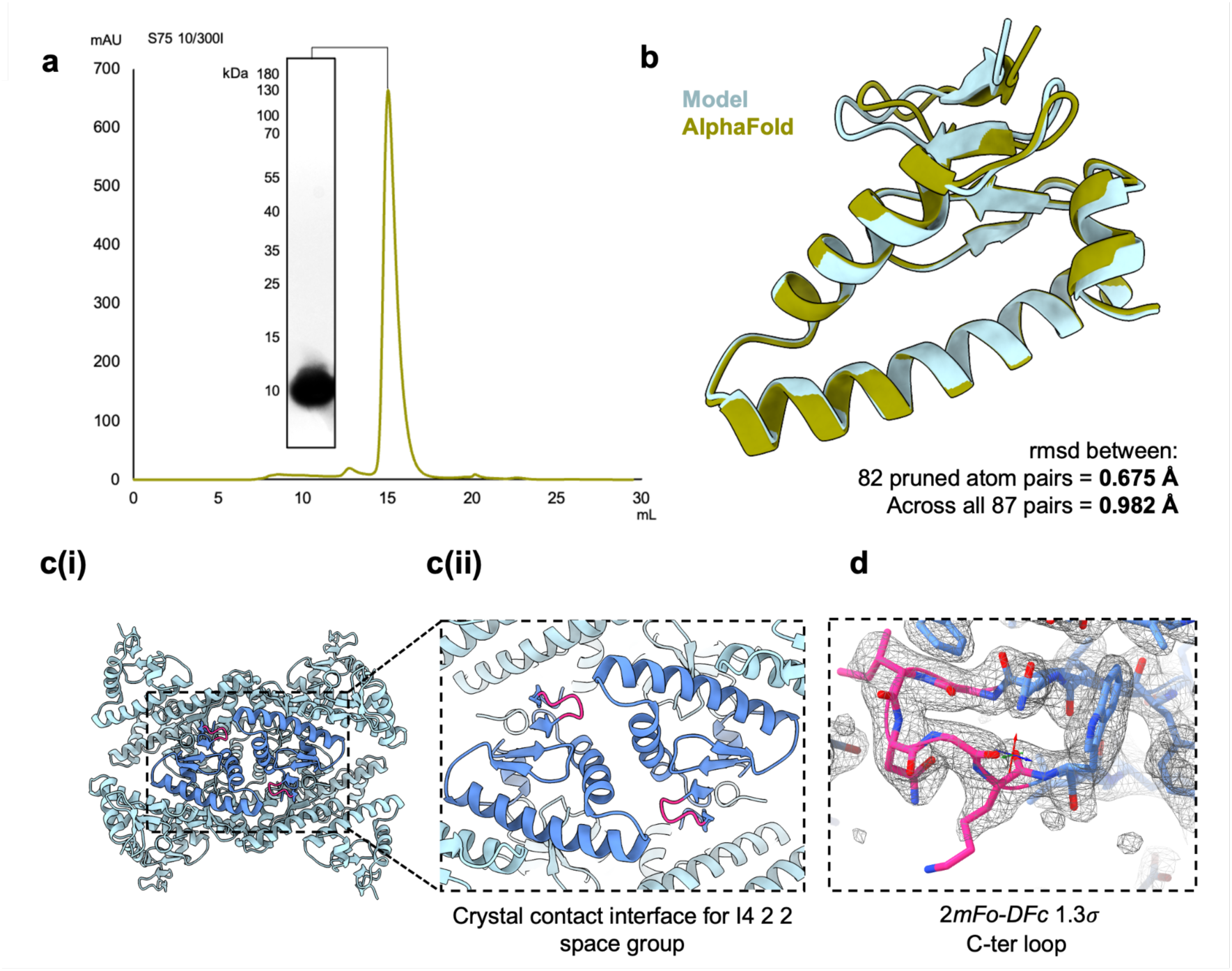
X-ray crystallography (XRC) YdbL model is concurrent with AlphaFold. **a.** Purification of YdbL for XRC. S75 10/300I SEC trace reveals a homogenous protein peak. Coomassie-stained SDS-PAGE shows full-length protein without any major contaminants or signs of degradation/aggregation. **b.** Overlay of AlphaFold2-YdbL (P76076) with 2.12 Å model reveals highly comparable secondary structure with an rmsd score below 1.0 Å across all atom pairs. **c. (i)** YdbL crystal contact interface with I4 2 2 space group for YdbL crystal lattice shown in sky blue with two representative YdbL monomers highlighted in cornflower blue. **(ii)** Magnified region showing the crystal-contact interfaces as corroborated by PDBePISA reveals the C-terminal loop (magenta) may be mobile and subject to an alternative conformation to the AlphaFold prediction in this packing lattice. d. The C-terminal loop is well represented by the electron density map for modelling in this conformation, with colors matching what is used in (c).

**Fig. S5.**
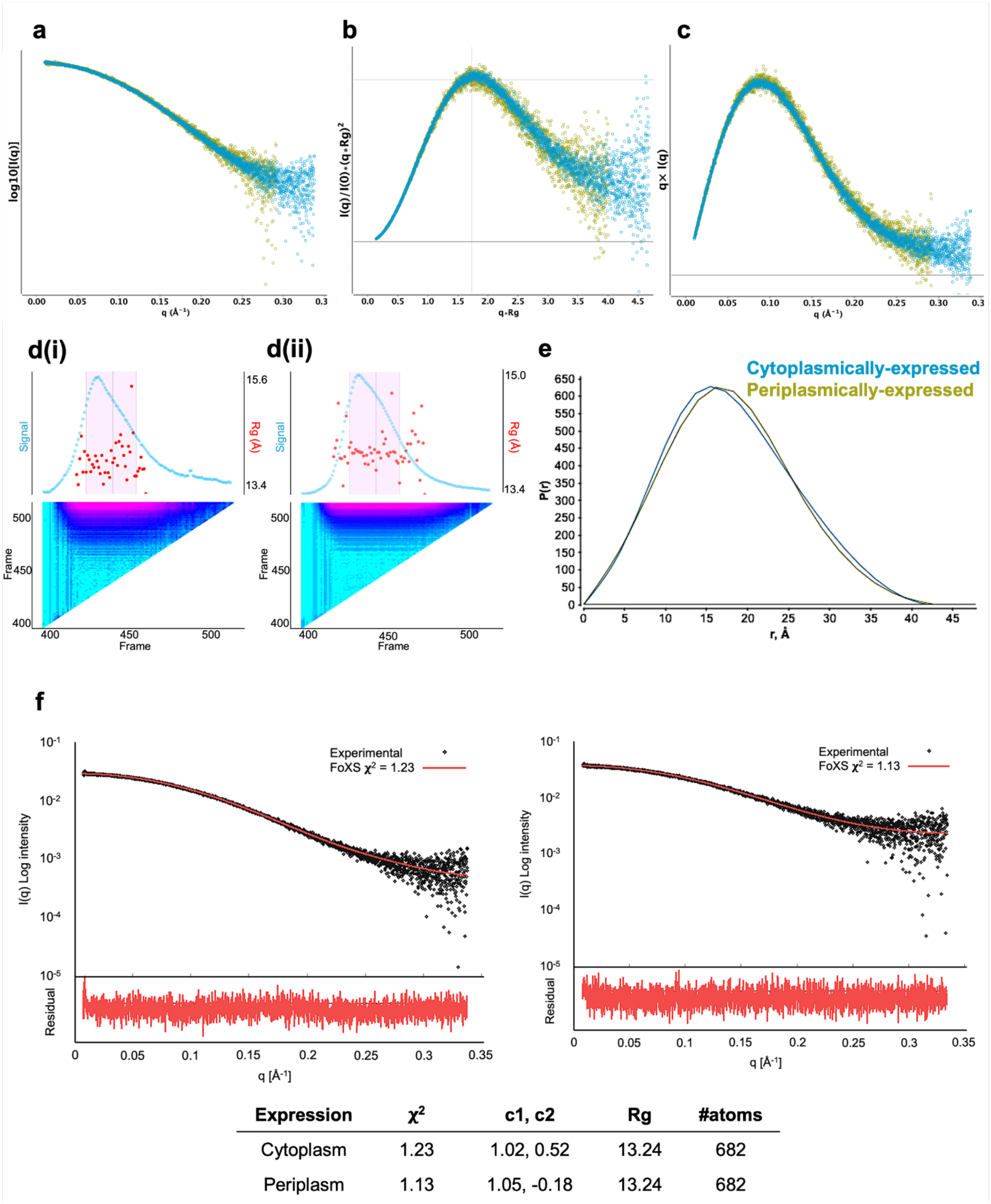
XRC structure matches solution structure via SEC-SAXS. **a.** Log10 SAXS intensity versus scattering vector, *q*. Plotted range represents the positive only data within the specified *q*-range. **b.** Dimensionless Kratky plot. Cross-hair marks the Guinier-Kratky point (1.732, 1.1), the main peak position for globular particles. **c.** Total scattered intensity plot. Plot readily demonstrates negative intensities at high-*q*. Over-subtraction of background leads to significant negative intensities. Likewise, under-subtraction can be observed as an elevated baseline at high-*q*. The horizontal line is drawn at y=0. **d.** SEC signal intensity plots for **(i)** cytoplasmically-expressed and **(ii)** periplasmically-expressed YdbL depict how Rg varies across frames, with the frame-by-frame panel below depicting their relative agreement, whereby cyan indicates high similarity and pink indicates low similarity. e. The plotted P(r) model function. Cytoplasmically- and periplasmically-expressed constructs are represented in blue and gold respectively. f. SEC-SAXS experimental data was analysed in comparison to the refined .pdb model using the FoXS server with cytoplasmically- and periplasmically-expressed data fitted to the crystal structure from left to right. Statistical fit of Chi-squared (**χ**2) and the free parameters - c1 (scaling of atomic radius) within limits of 0.99 ≤ c1 ≤ 1.05 and c2 (adjustment of hydration layer and bulk water densities) within limits of -2.0 ≤ c2 ≤ 4.0 are also shown below for both experimental datasets. Rg and number of atoms in the crystal structure are also recorded.

**Fig. S6.**
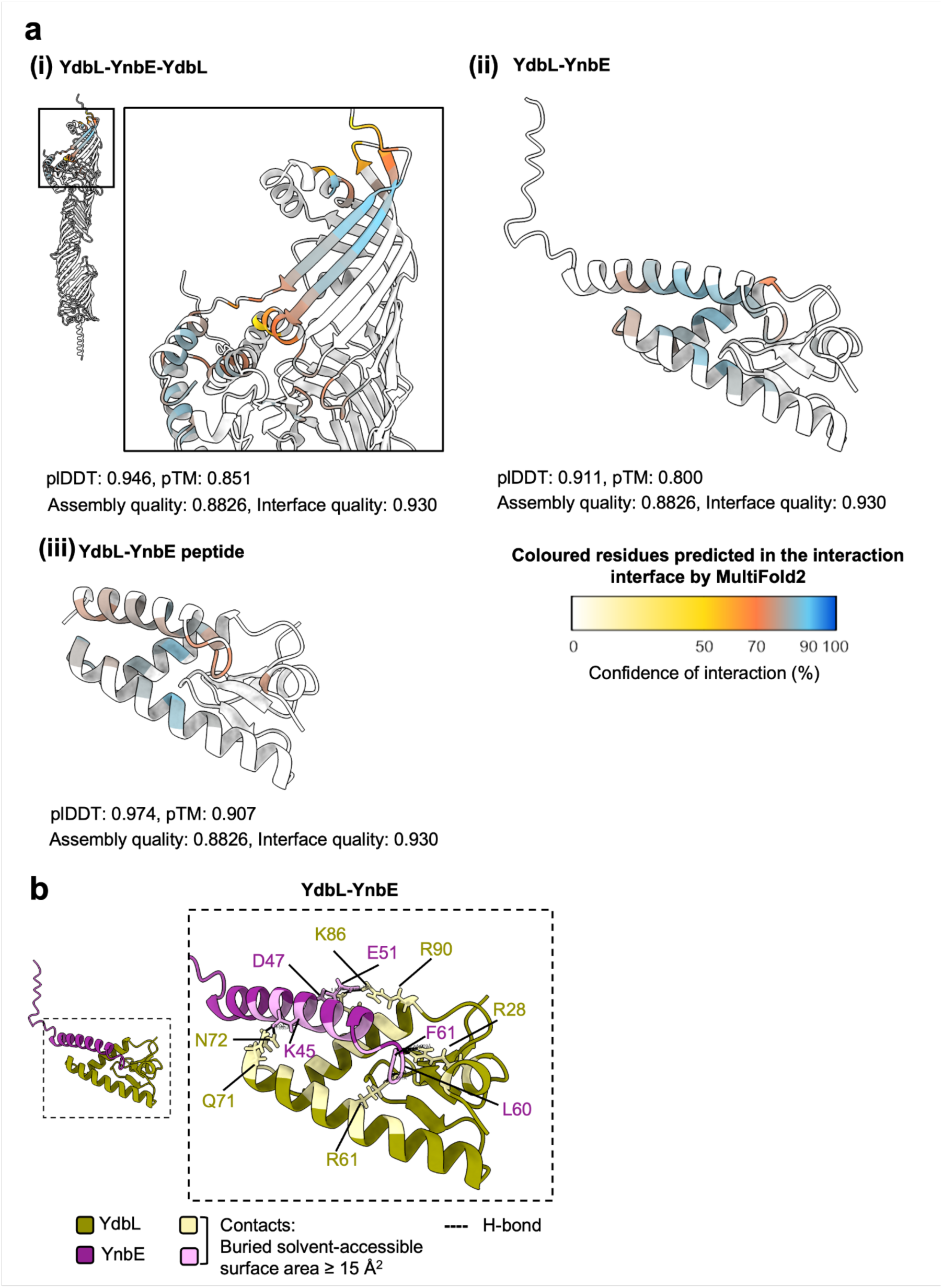
MultiFold2- and ModFolddock2-predicted interaction interfaces between YdbH, YnbE and YdbL. **a.** Confidence of MultiFold2-predicted interaction interfaces by ModFolddock2 for **(i)** YdbH-YnbE-YdbL, **(ii)** YdbL-YnbE and **(iii)** YdbL-YnbE peptide. Coloured residues are predicted to form the interaction interfaces between proteins. Their colour is scaled by the confidence of interaction with 90-100% being highly likely versus lower confidence predictions around 50%. b. Predicted MultiFold2 interface for YdbL-YnbE coloured by protein (YdbL in gold, YnbE in plum). A magnified view of the interaction interface reveals the contact region and predicted hydrogen bonds. Residues predicted to mediate direct interactions are labelled.

**Fig. S7.**
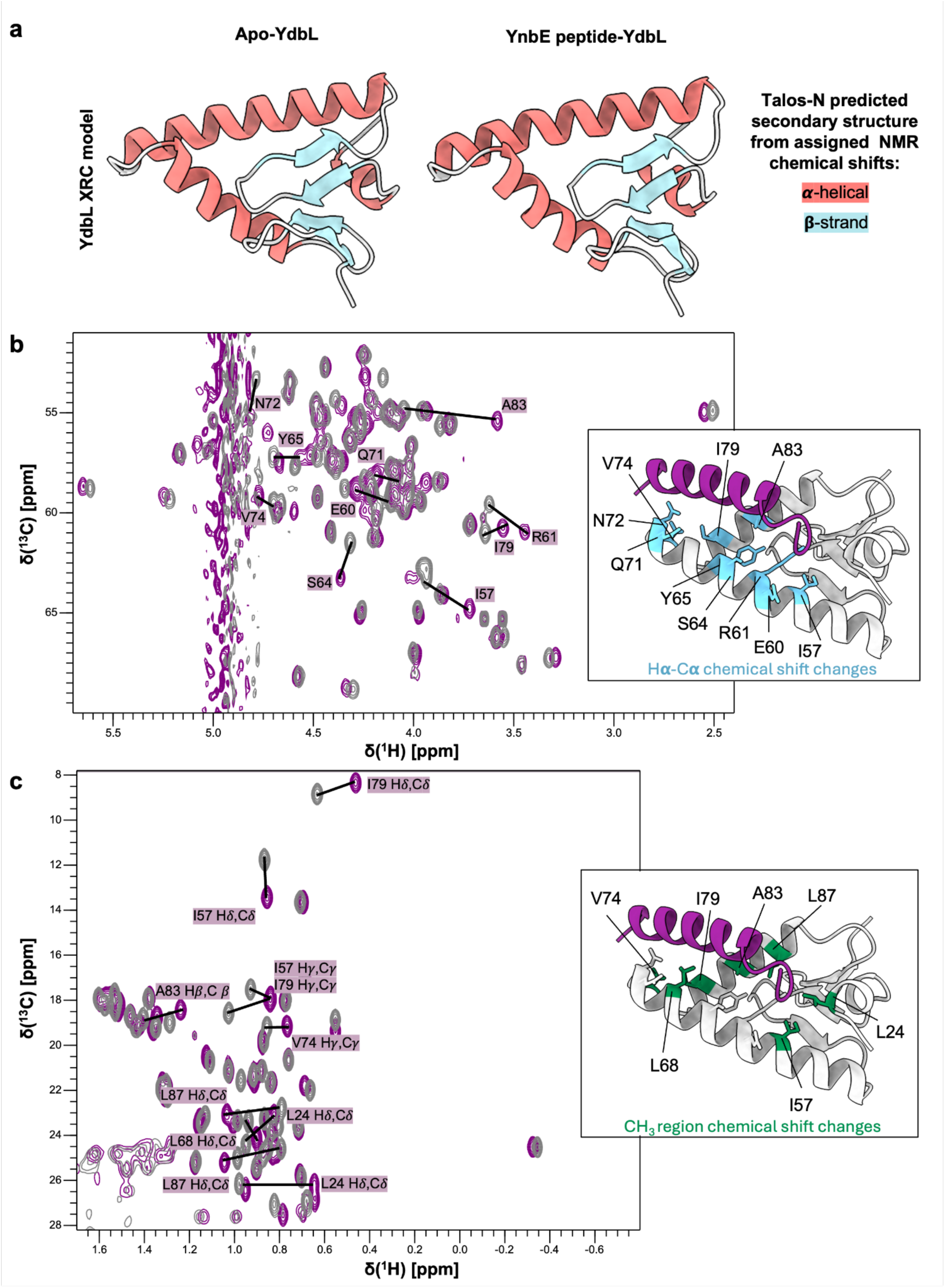
Analysis of secondary structure and chemical shift changes between apo- and peptide-bound YdbL did not reveal any significant conformational changes. **a.** Talos-N secondary structure prediction for apo- (left) vs peptide-bound (right) YdbL mapped onto the crystal model of YdbL. Red depicts **α**-helical structure, blue depicts **β**-strand structure and white depicts loops or unstructured regions. Residues are identified as **α**-helical or **β**-strand if the Talos-N analysis of backbone chemical shifts assigns a helix or strand probability of >0.5, respectively. **b.** Overlaid 2D ^1^H-^13^C HSQC spectra showing the H**α**-C**α** region for apo- (grey) and peptide-bound (purple) YdbL. The H**α**-C**α** peaks that are labelled show some of the most significant chemical shift changes that occur upon peptide binding. Most of these residues also show large amide shift changes in Figure 3(c). Interestingly, most of the residues highlighted in this spectrum show an increase in the C**α** chemical shift upon peptide binding. The YdbL-YnbE peptide MultiFold2 model shown to the right of the spectra is labelled with residues in blue indicating chemical shift differences for H**α**-C**α** at 0.1 ppm or more. Since they are located in helical regions, this increase in C**α** chemical shift may indicate a stabilization of the helical structure rather than a significant conformational change; this observation is in agreement with the MultiFold2 predictions (Figure 2(d)). **c.** Overlaid 2D ^1^H-^13^C HSQC spectra showing the methyl (CH_3_) region for apo- (grey) and peptide-bound (purple) YdbL. Labelled peaks focus on the most significant chemical shift changes. As shown by the model to the right of the spectra, the methyl groups showing large chemical shift changes are all located in the identified peptide binding site (colored green), with many of these residues also showing the most significant amide shift changes shown in Figure 3(c). The observed changes in methyl group chemical shifts are likely to arise from small side chain conformational rearrangement due to direct contact between these groups and residues in the YnbE peptide and/or from movements of nearby aromatic amino acid side chains which lead to through-space changes in chemical shifts.

**Fig. S8.**
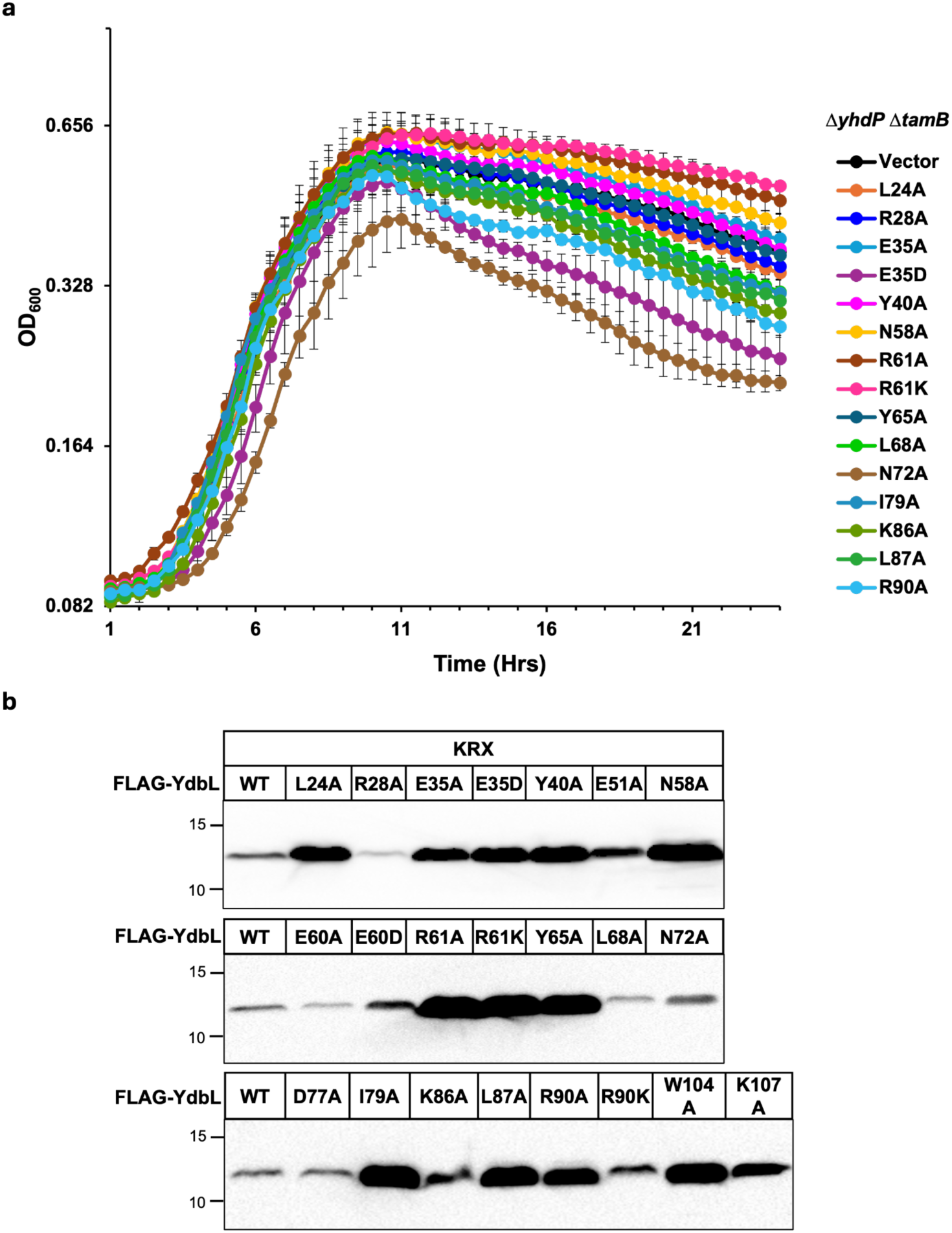
Growth curves and immunoblots of YdbL variants. **a.** Growth curves for non-functional YdbL variants. Overnight cultures of the Δ*tamB* Δ*yhdP* double mutant carrying FLAG-YdbL variants encoded on pET23/42 plasmid were diluted (1:1000) in LB containing ampicillin and grown at 37°C for 24 hrs. OD_600_ values are represented on a logarithmic scale. Data represent the average and standard deviation from three biological replicates. **b.** Immunoblots showing the cellular levels of YdbL variants. KRX strains carrying FLAG-YdbL variants encoded on the pET23/42-derived plasmid were grown to log phase and then for three additional hours at 37 ⁰C after induction with 0.2% rhamnose. Whole-cell lysate samples were prepared as described in the Materials and Methods section, and proteins were separated on 15% SDS-polyacrylamide gel and probed with α-FLAG antibodies by immunoblotting. Most, but not all, non-functional YdbL variantsshowed increased levels compared to wild-type YdbL (WT). Molecular mass markers (in kDa) are shown on the left. All data are representative of three independent experiments.

**Fig. S9.**
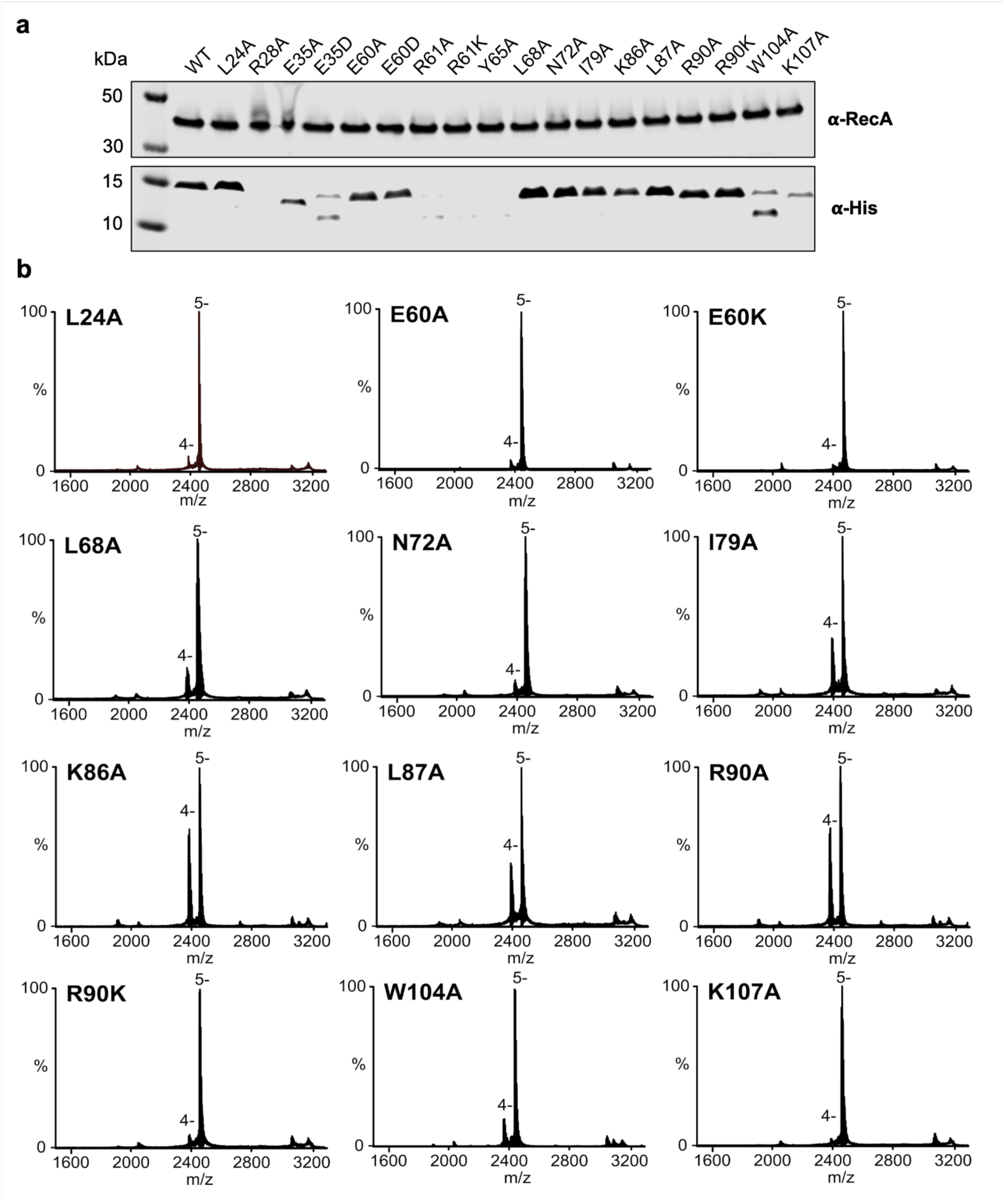
nMS data for all mutants. **a.** Expression test for all YdbL variant proteins with substitutions at the YnbE-binding interface. Recombinantly expressed protein variants were analyzed via immunoblot against His-YdbL (*α*-His) from whole- cell lysates. An *α*-RecA immunoblot was included for a loading control. The expression test is representative across three biological replicates. **b.** Native mass spectra of each mutant in the presence of 3.5 µM peptide. The highest intensity charge state series are labelled: apo(4-) and peptide-bound (5-) Spectra are representative across three technical replicates.

**Fig. S10.**
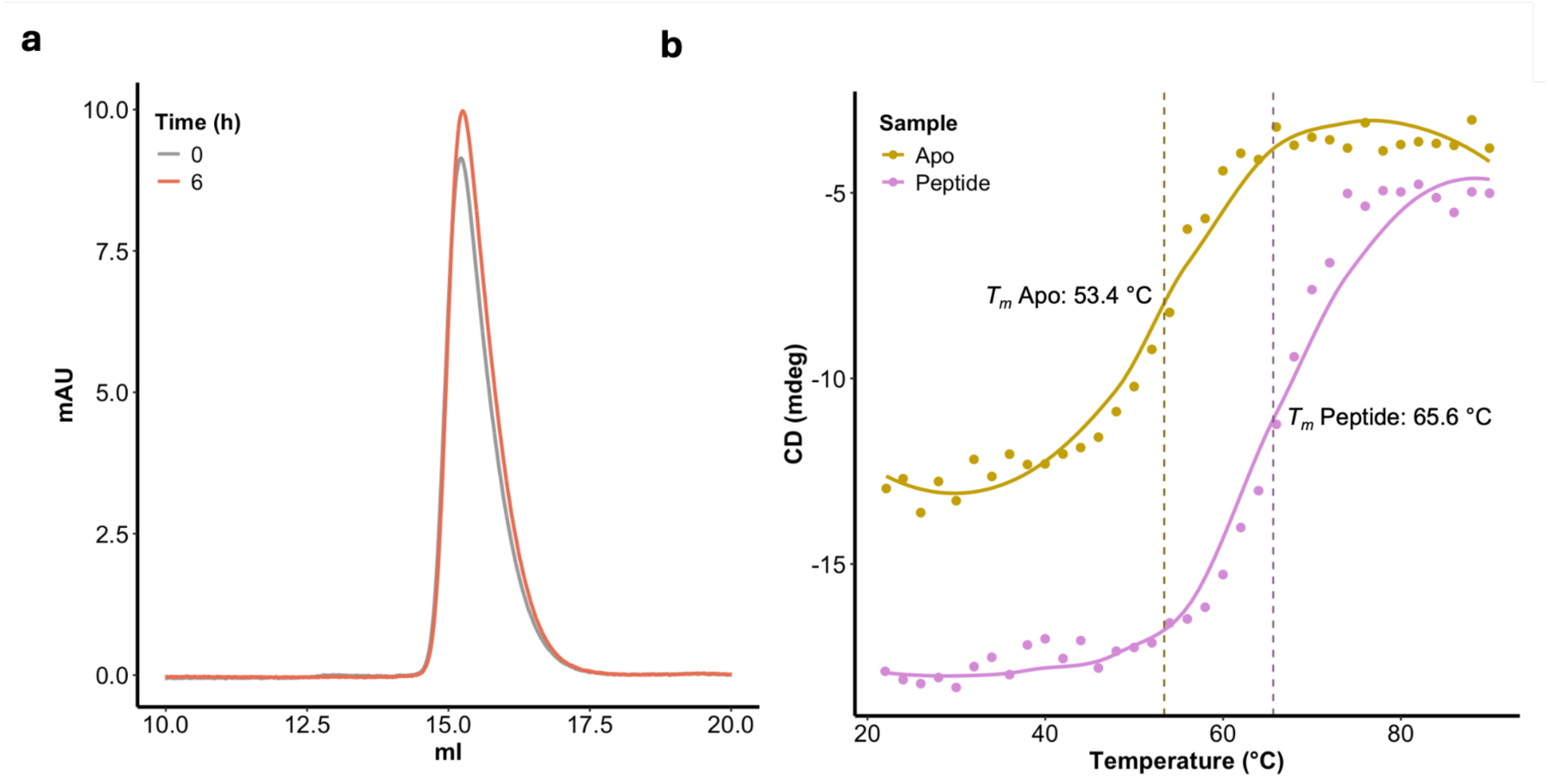
Biophysics for YdbL reveals it is thermostable, folded and does not aggregate in the DegP degradation assay conditions. **a.** SEC aggregation assay. YdbL was incubated at 37 °C for 0 and 6 hours in standard assay buffer before loading onto an S75 10/300I column. At both timepoints, YdbL eluted as a monodisperse peak with no visible aggregate peak or loss of folded sample. **b.** Thermal denaturation analysis shows peptide-bound YdbL has a higher thermal stability than apo-YdbL. The CD melt curves for apo-YdbL (gold) and peptide-bound YdbL (purple) are plotted as the circular dichroism millidegrees against temperature. A wavelength of 222 nm was selected to monitor loss of *α*-helical secondary structure with dots used to show raw data with scans from 22-90 °C in 2 °C step intervals. The solid lines depict the smoothened melting curve. The midpoint of sharp melting transition occurs at 53.4 °C for apo-YdbL and 65.6 °C for peptide-bound YdbL.

## Tables

**Table S1.** Strains used in this study.

| Strain | Genotype | Source |
| --- | --- | --- |
| DH5α | F<sup>-</sup> ϕ80/*lacZ*ΔM15 $\Delta$(*lacZYA-argF*)U169 *recA1 endA1 hsdR17* (*rK*<sup>-</sup>, *mK*<sup>+</sup>) *phoA supE44* $\lambda$ *thi-1* | Life Technologies |
| T7 Express | *fhuA2 lacZ::T7 gene1 [lon] ompT gal sulA11 R(mcr-73::miniTn10--Tet<sup>S</sup>)2 [dcm] R(zgb-210::Tn10--Tet<sup>S</sup>) endA1 $\Delta$(*mcrC-mrr*)114::IS10* | NEB |
| MG1655 | Wild type | (1) |
| NR5161 | Δ*yhdP::frt* Δ*tamB::frt* | (2) |
| KRX | [F', traD36, Δ*ompP*, *proA+B*<sup>+</sup>, *lacIq*, $\Delta$(*lacZ*)M15] Δ*ompT*, *endA1*, *recA1*, *gyrA96* (*Nal*<sup>r</sup>), *thi-1*, *hsdR17* (*rK*<sup>-</sup>, *mK*<sup>+</sup>), *e14*<sup>-</sup> (*McrA*<sup>-</sup>), *relA1*, *supE44*, $\Delta$(*lac-proAB*), $\Delta$(*rhaBAD*)::T7 RNA polymerase. | Promega |
| NR5530 | KRX (pKD46) | This study |
| NR6436 | MG1655 Δ*ydbH-ydbL::kan* | (3) |
| NR6658 | Δ*yhdP::frt* Δ*tamB::frt* (pET23/42) | (3) |
| NR7056 | NR5161 (pET23/42GST-His<sub>2</sub>-YdbH/YnbE/YdbL) | (3) |
| NR7254 | NR5161 (pET23/42GST-His<sub>2</sub>-YdbH) | (3) |
| NR7258 | MG1655 (pET23/42GST-His<sub>2</sub>-YdbH/ΔYnbE/YdbL) | (3) |
| NR7352 | NR5161 (pET23/42 GST-His<sub>2</sub>-YdbH /YnbE/FLAG-YdbL) | (3) |
| NR7359 | MG1655 Δ*ydbH-ydbL::kan* (pSup-BpaRS-6TRN) | (3) |
| NR7817 | NR5161 (pET23/42 pET23/42GST-His<sub>2</sub>-YdbH/YnbE-FLAG<sub>STOP</sub>) | (3) |
| NR7818 | NNR5161 (pET23/42YnbE-FLAG<sub>STOP</sub>) | (3) |
| NR7859 | KRX (pET23/42FLAGYdbL) | This study |
| NR7867 | KRX (pET23/42GST-His<sub>2</sub>-YdbH /YnbE/FLAG-YdbL) | This study |
| NR7868 | MG1655 (pET23/42YdbL) | (3) |
| NR7885 | MG1655 $\Delta yhdP::frt$ (pET23/42YdbL) | (3) |
| NR7890 | MG1655 $\Delta tamB::frt$ (pET23/42YdbL) | (3) |
| NR7902 | NR5161 (pET23/42GST-His<sub>2</sub>-YdbH/YnbE<sub>STOP</sub>) | (3) |
| NR7980 | KRX $\Delta ydbH-ydbL::kan$ | This study |
| NR8194 | KRX (pET23/42FLAG/YdbL/E35A) | This study |
| NR8195 | KRX (pET23/42FLAG/YdbL/R61A) | This study |
| NR8196 | KRX (pET23/42FLAG/YdbL/Y65A) | This study |
| NR8197 | KRX (pET23/42FLAG/YdbL/W104A) | This study |
| NR8198 | KRX (pET23/42FLAG/YdbL/K107A) | This study |
| NR8202 | NR5161 (pET23/42FLAG/YdbL/E35A) | This study |
| NR8203 | NR5161 (pET23/42FLAG/YdbL/R61A) | This study |
| NR8204 | NR5161 (pET23/42FLAG/YdbL/Y65A) | This study |
| NR8212 | NR5161 (pET23/42FLAG/YdbL/R28A) | This study |
| NR8213 | NR5161 (pET23/42FLAG/YdbL/Y40A) | This study |
| NR8214 | KRX (pET23/42FLAG/YdbL/R28A) | This study |
| NR8215 | KRX (pET23/42FLAG/YdbL/Y40A) | This study |
| NR8224 | NR5161 (pBAD33SpecR) | This study |
| NR8225 | NR5161 (pBAD33-SpecR-NativeYdbH/YnbE/YdbL) | This study |
| NR8226 | NR5161 [pBAD33-SpecR-RBS-YdbH/YnbE/YdbL] | This study |
| NR8227 | NR5161 [pBAD33-SpecR-RBS-YdbL] | This study |
| NR8278 | NR5161 [pBAD33SpecR-RBS-YdbL;pET23/42] | This study |
| NR8279 | NR5161 [pBAD33SpecR-RBS-YdbL; pET23/42GST-His<sub>2</sub>-YdbH/YnbE/YdbL] | This study |
| NR8280 | NR5161 [pBAD33SpecR-RBS-YdbL; pET23/42GST-His<sub>2</sub>-YdbH/YnbE-FLAG/YdbL] | This study |
| NR8281 | NR5161 [pBAD33SpecR-RBS-YdbL; pET23/42GST-His<sub>2</sub>-YdbH/YnbE<sub>STOP</sub>] | This study |
| NR8282 | NR5161 [pBAD33SpecR-RBS-YdbL; pET23/42GST-His<sub>2</sub>-YdbH/YnbEFLAG<sub>STOP</sub>] | This study |
| NR8283 | NR5161 [pBAD33SpecR-RBS-YdbL; pET23/42GST-His<sub>2</sub>-YdbH] | This study |
| NR8284 | NR5161 [pBAD33SpecR-RBS-YdbL; pET23/42YnbEFLAG<sub>STOP</sub>] | This study |
| NR8335 | NR5161 [pBAD33SpecR-RBS-YdbL; pET23/42YnbE<sub>STOP</sub>] | This study |
| NR8357 | MG1655 $\Delta ydbH\text{-}\Delta ydbL::kan$ [pSUP-BPARS-6TRN; GST-His<sub>2</sub>-YdbH/YnbE-FLAG(K27Am)<sub>STOP</sub>; pBAD33-SpecR] | This study |
| NR8358 | MG1655 $\Delta ydbH\text{-}\Delta ydbL::kan$ [pSUP-BPARS-6TRN; GST-His<sub>2</sub>-YdbH/YnbE(K27Am)-FLAG<sub>STOP</sub>; pBAD33-SpecR-RBS-YdbL] | This study |
| NR8359 | MG1655 $\Delta ydbH\text{-}ydbL::kan$ [pSUP-BPARS-6TRN; GST-His<sub>2</sub>-YdbH/YnbE(D47Am)-FLAG<sub>STOP</sub>; pBAD33-SpecR] | This study |
| NR8360 | MG1655 $\Delta ydbH\text{-}ydbL::kan$ [pSUP-BPARS-6TRN; GST-His<sub>2</sub>-YdbH/YnbE(D47Am)-FLAG<sub>STOP</sub>; pBAD33-SpecR-RBS-YdbL] | This study |
| NR8383 | KRX (pET23/42FLAG-YdbL/R61K) | This study |
| NR8384 | KRX (pET23/42FLAG-YdbL/N58A) | This study |
| NR8385 | KRX (pET23/42FLAG-YdbL/E35D) | This study |
| NR8391 | MG1655 $\Delta ydbH\text{-}ydbL::kan$ [pSUP-BPARS-6TRN; GST-His<sub>2</sub>-YdbH/YnbE(E41Am)-FLAG<sub>STOP</sub>; pBAD33SpecR] | This study |
| NR8392 | MG1655 $\Delta ydbH\text{-}ydbL::kan$ [pSUP-BPARS-6TRN; pET23/42 GST-His<sub>2</sub>-YdbH/YnbE(E41Am)-FLAG<sub>STOP</sub>; pBAD33SpecR-RBS-YdbL] | This study |
| NR8395 | MG1655 $\Delta ydbH\text{-}ydbL::kan$ [pSUP-BPARS-6TRN; GST-His<sub>2</sub>-YdbH/YnbE(F61Am)-FLAG<sub>STOP</sub>; pBAD33SpecR] | This study |
| NR8396 | MG1655 $\Delta ydbH\text{-}ydbL::kan$ [pSUP-BPARS-6TRN; pET23/42GST-His<sub>2</sub>-YdbH/YnbE(F61Am)-FLAG<sub>STOP</sub>; pBAD33SpecR-RBS-YdbL] | This study |
| NR8455 | KRX $\Delta ydbH\text{-}ydbL::kan$ (pET23/42FLAG-YdbL) | This study |
| NR8456 | KRX $\Delta ydbH\text{-}ydbL::kan$ (pET23/42GST-His<sub>2</sub>-YdbH/YnbE/FLAG-YdbL) | This study |
| NR8460 | NR5161 (pET23/42FLAG-YdbL/E35D) | This study |
| NR8461 | NR5161 (pET23/42FLAG-YdbL/N58A) | This study |
| NR8462 | NR5161 (pET23/42FLAG-YdbL/R61K) | This study |
| NR8484 | KRX $\Delta ydbH-ydbL::kan$ (pET23/42FLAG-YdbL; pBAD33SpecR) | This study |
| NR8485 | KRX $\Delta ydbH-ydbL::kan$ [pET23/42FLAG-YdbL; pBAD33SpecRBS-YdbH/YnbE] | This study |
| NR8501 | KRX (pET23/42FLAGYdbL/E51A) | This study |
| NR8502 | KRX (pET23/42FLAGYdbL/N72A) | This study |
| NR8503 | KRX (pET23/42FLAGYdbL/D77A) | This study |
| NR8504 | KRX (pET23/42FLAGYdbL/K86A) | This study |
| NR8598 | KRX $\Delta ydbH-ydbL::kan$ [pET23/42FLAG-YdbL; pBAD33Spec-RBS-YnbE] | This study |
| NR8609 | KRX $\Delta ydbH-ydbL::kan$ [pET23/42FLAG-YdbL; pBAD33SpecRBS-YdbH] | This study |
| NR8642 | NR5161 (pET23/42FLAG-YdbL/L68A) | This study |
| NR8643 | NR5161 (pET23/42FLAG-YdbL/I79A) | This study |
| NR8644 | NR5161 (pET23/42FLAG-YdbL/L87A) | This study |
| NR8645 | NR5161 (pET23/42FLAG-YdbL/R90A) | This study |
| NR8650 | KRX (pET23/42FLAG-YdbL/E60A) | This study |
| NR8651 | KRX (pET23/42FLAG-YdbL/E60D) | This study |
| NR8652 | KRX (pET23/42FLAG-YdbL/L68A) | This study |
| NR8653 | KRX (pET23/42FLAG-YdbL/I79A) | This study |
| NR8654 | KRX (pET23/42FLAG-YdbL/L87A) | This study |
| NR8655 | KRX (pET23/42FLAG-YdbL/R90A) | This study |
| NR8656 | KRX (pET23/42FLAG-YdbL/R90K) | This study |
| NR8673 | NR5161 (pET23/42FLAG-YdbL/L24A) | This study |
| NR8674 | NR5161 (pET23/42FLAG-YdbL/N72A) | This study |
| NR8675 | NR5161 (pET23/42FLAG-YdbL/K86A) | This study |

**Table S2:**
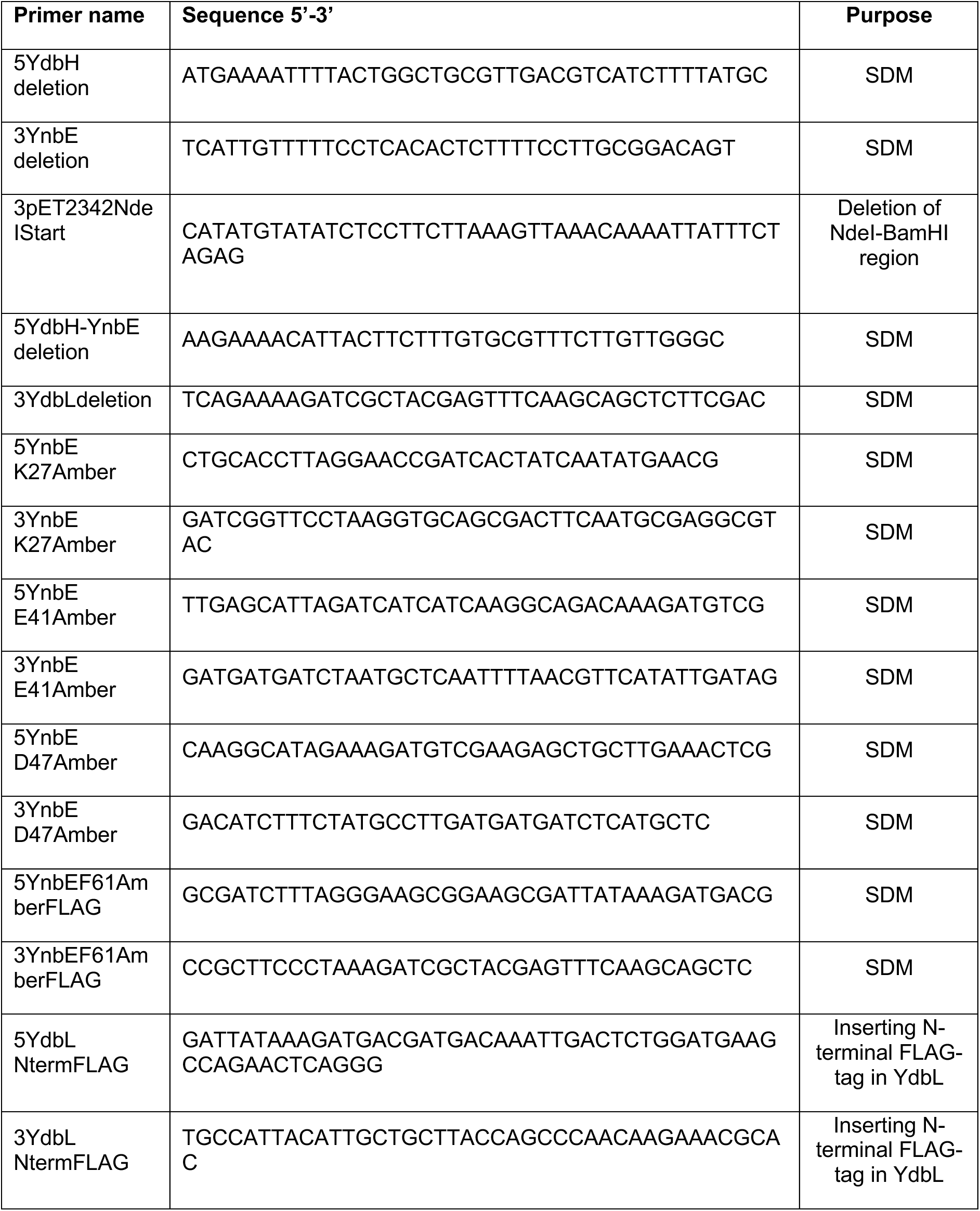
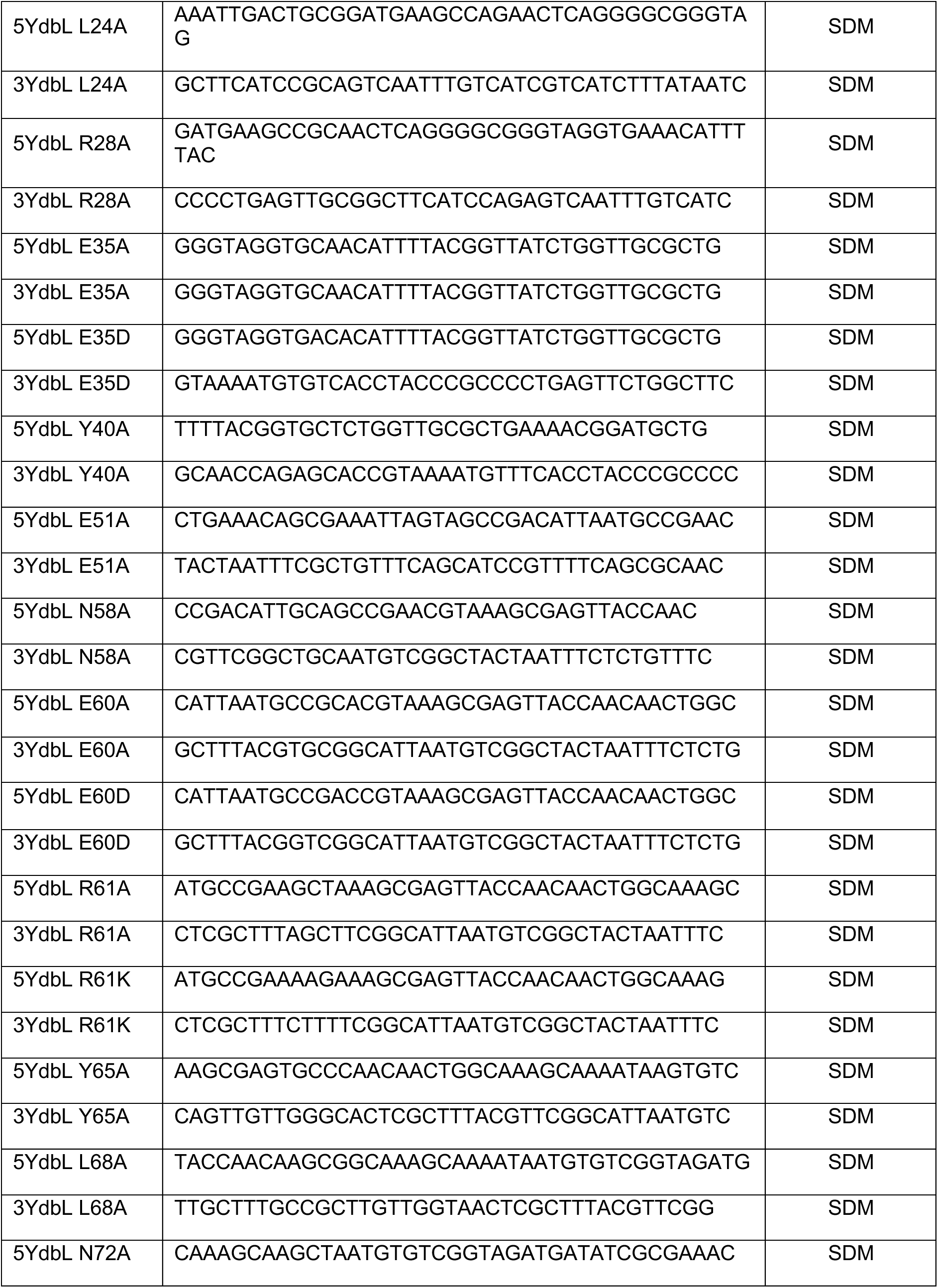
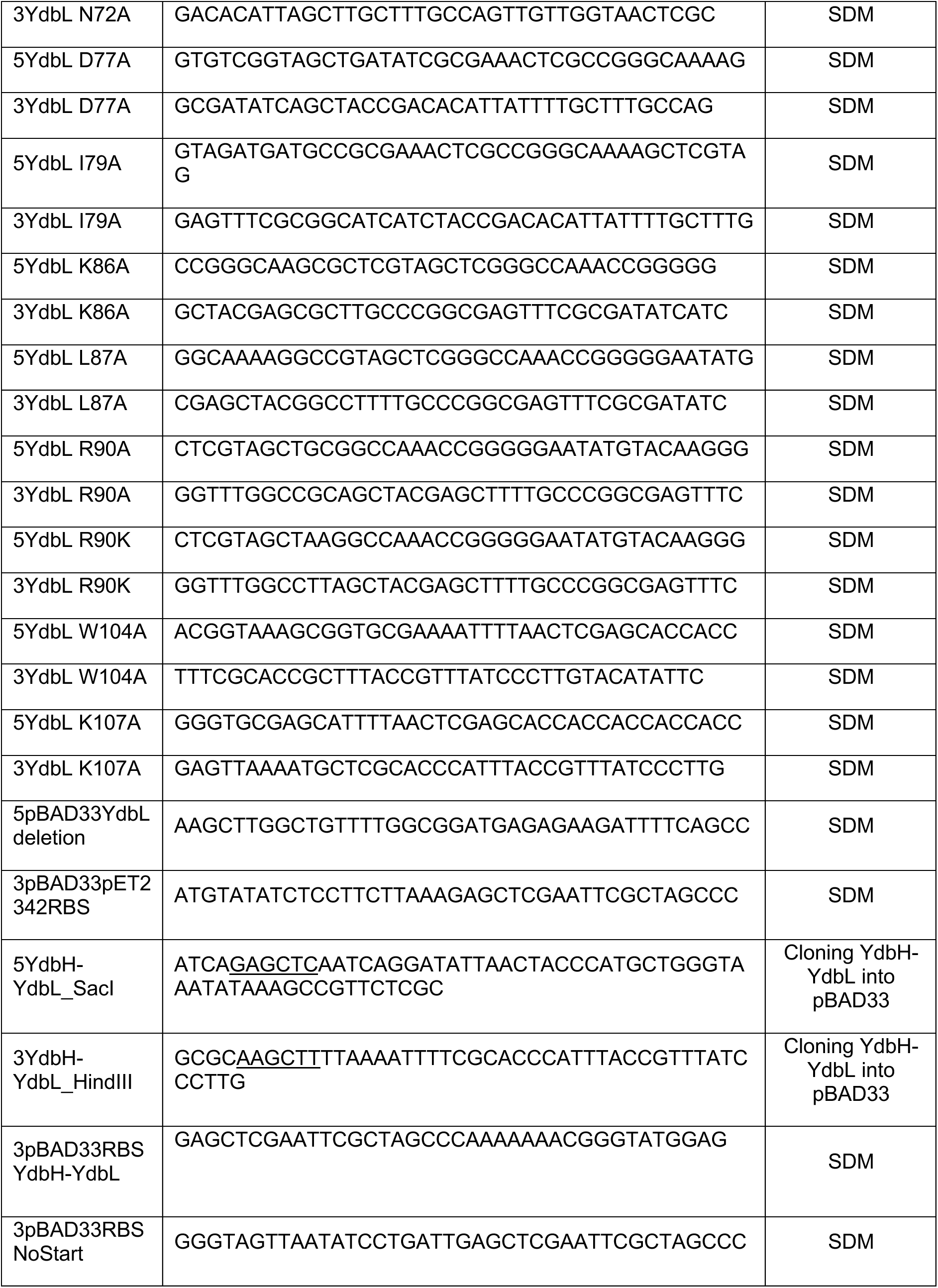
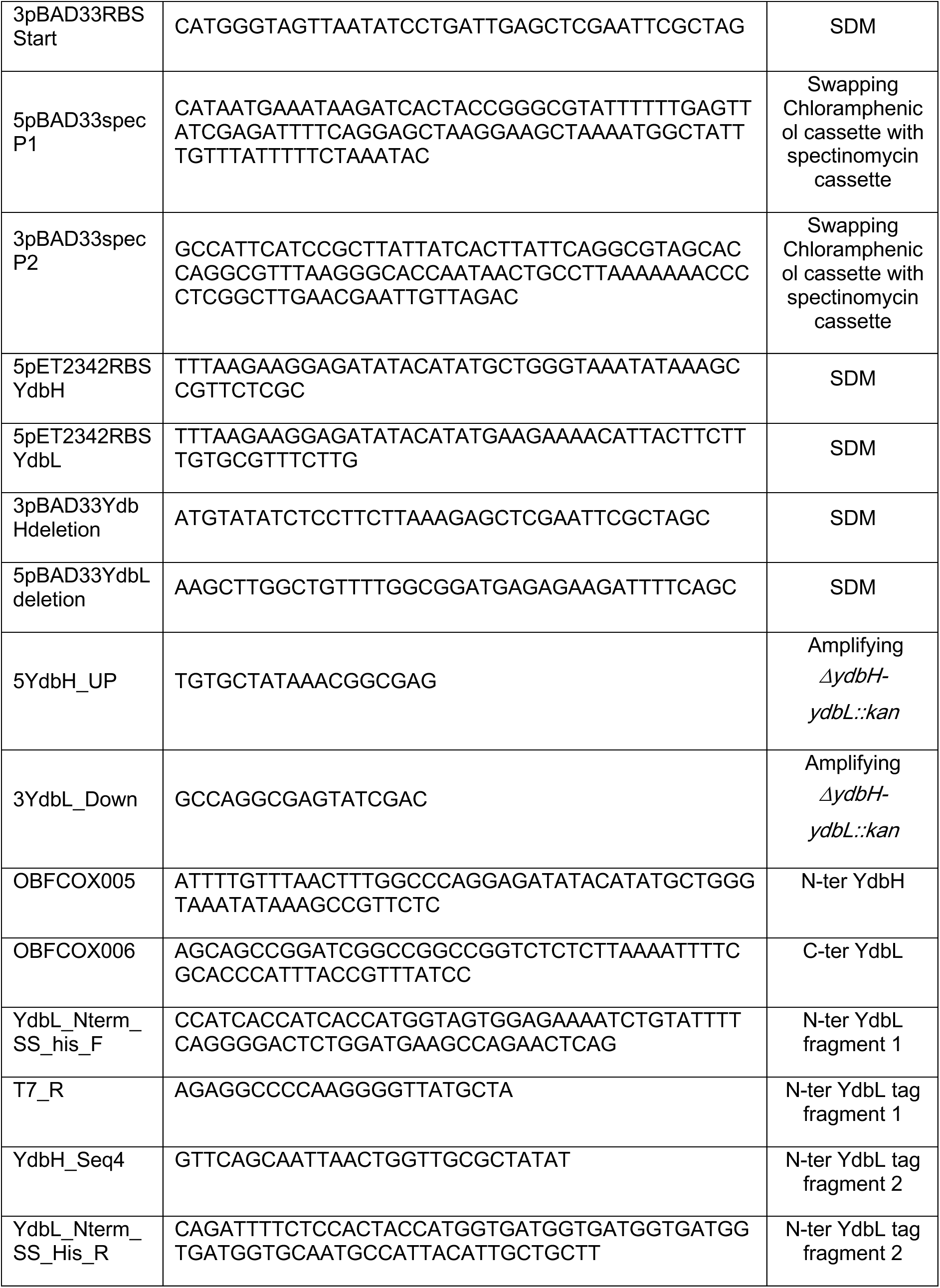
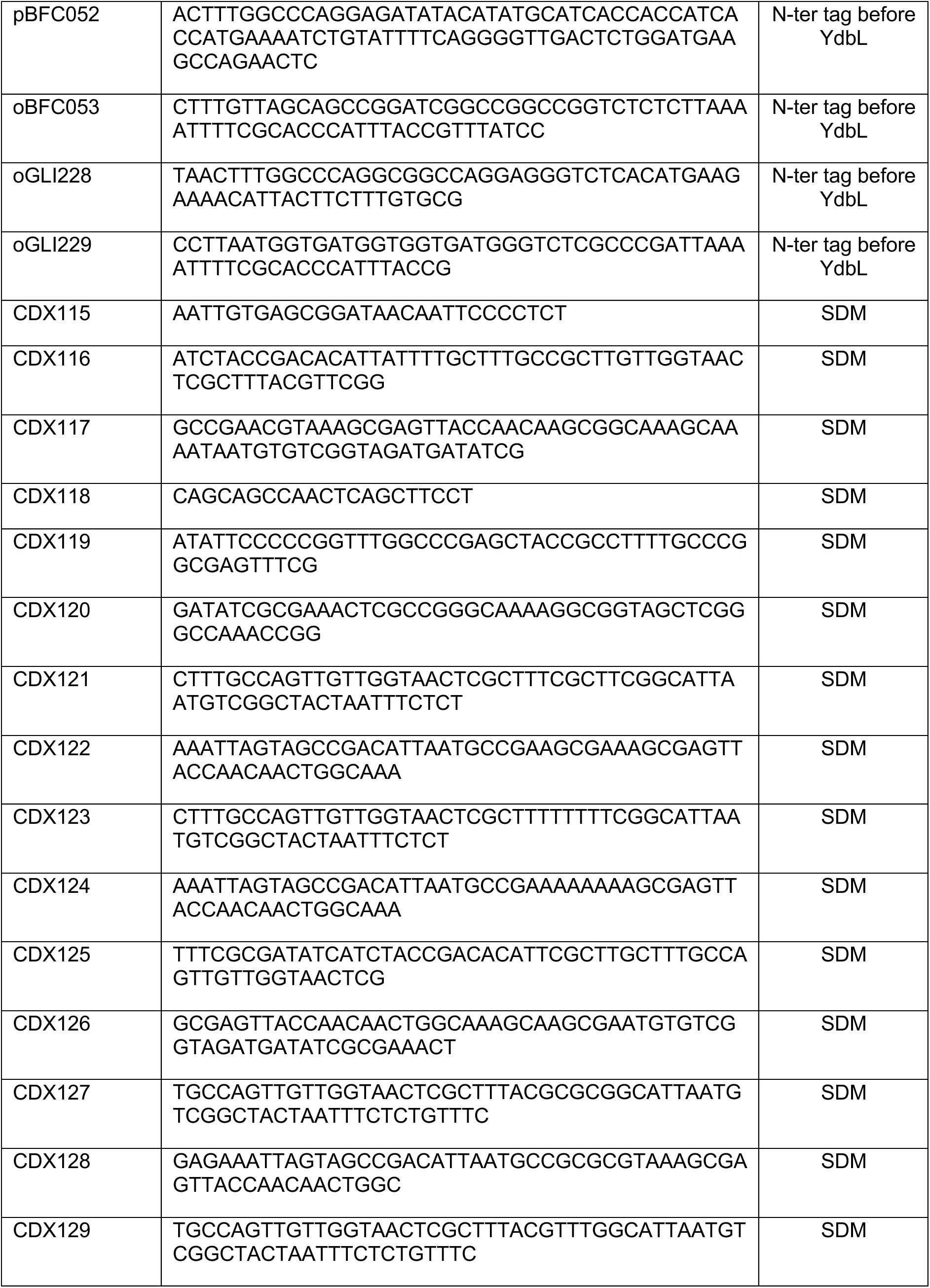
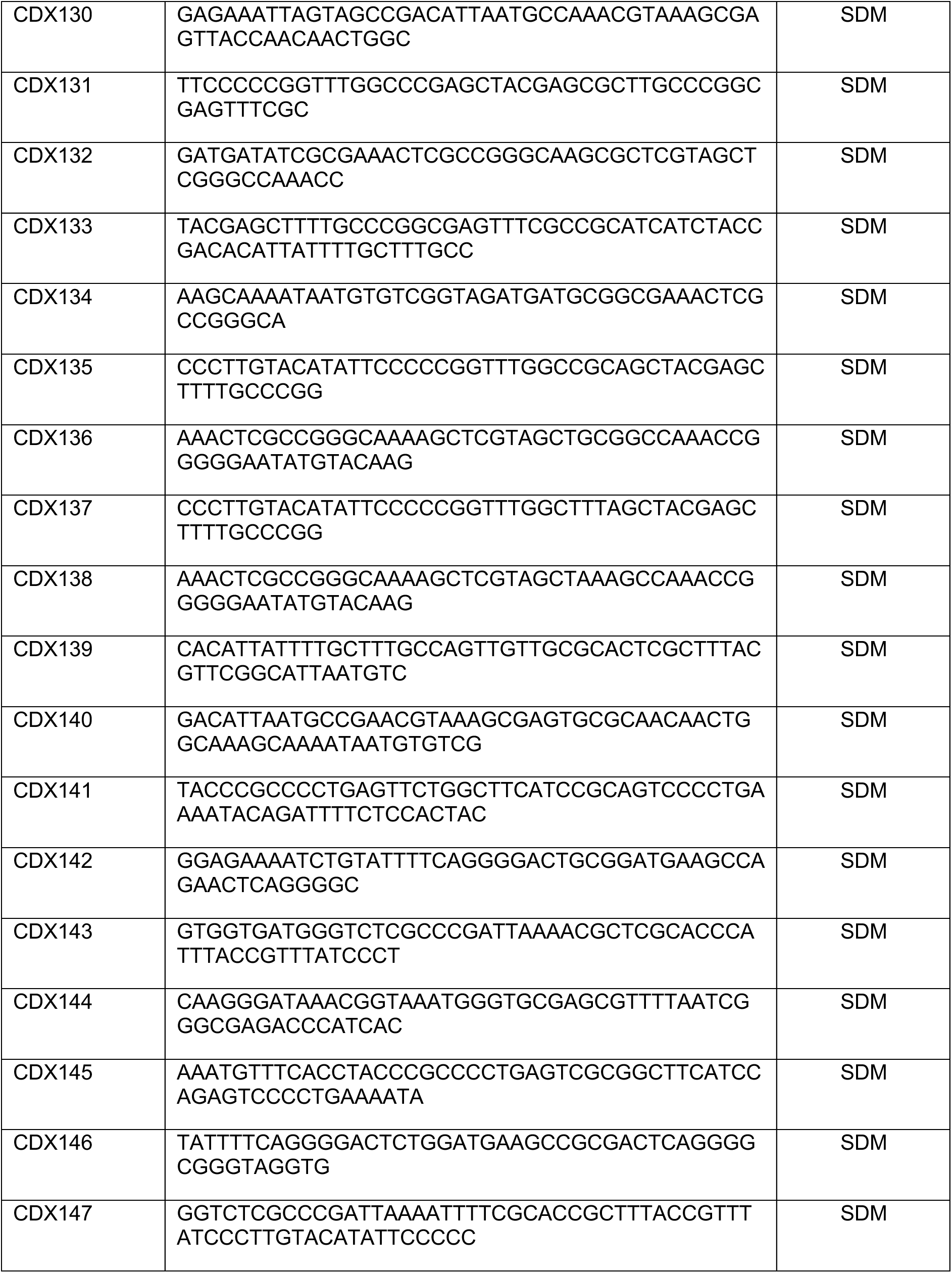
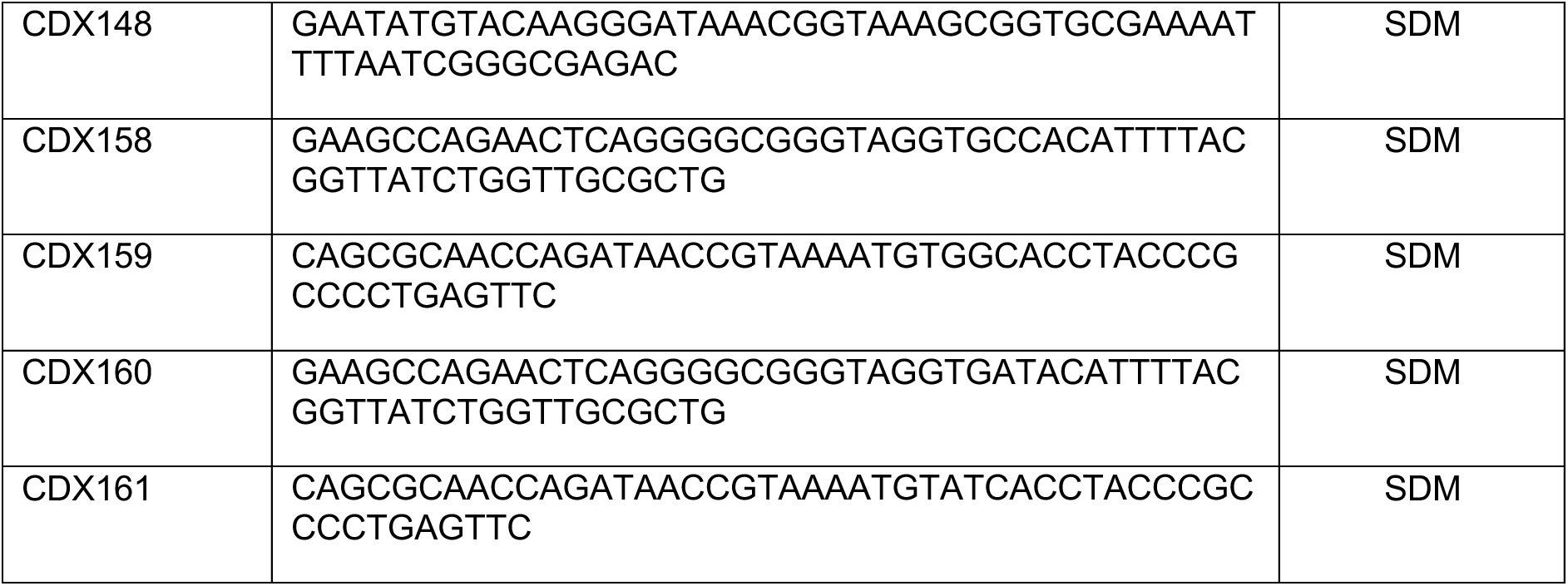
Primers used in the study.

**Table S3:**
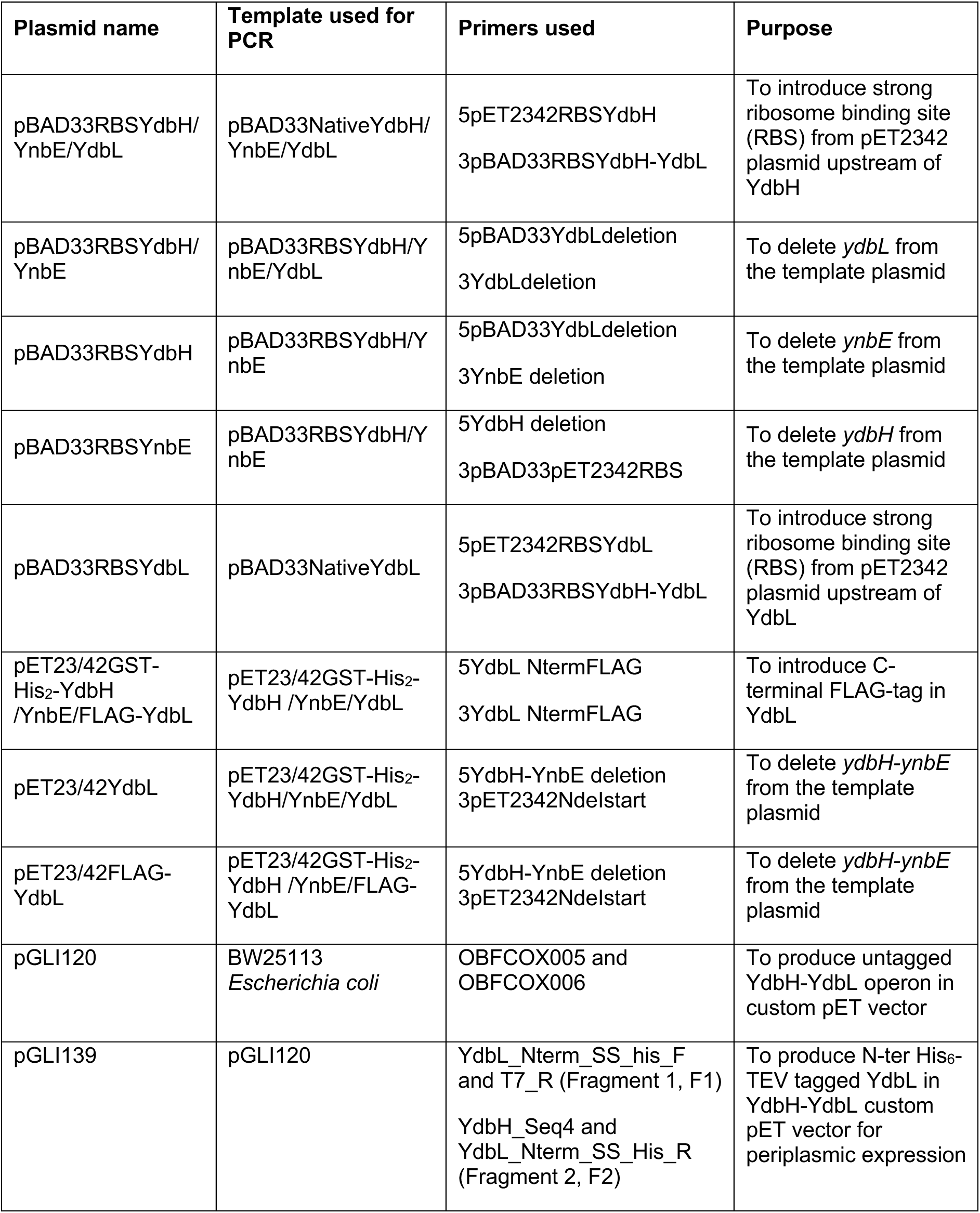
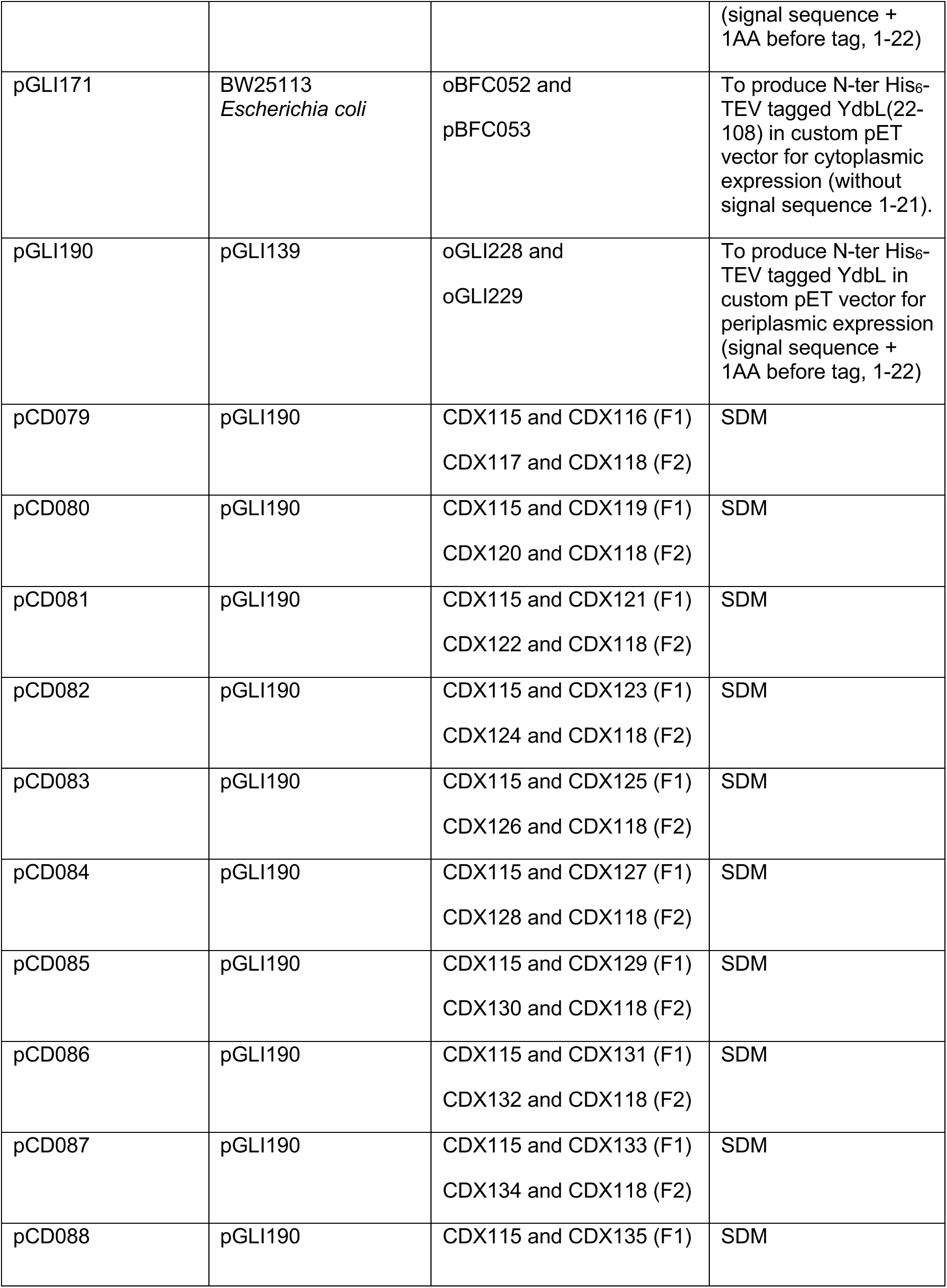
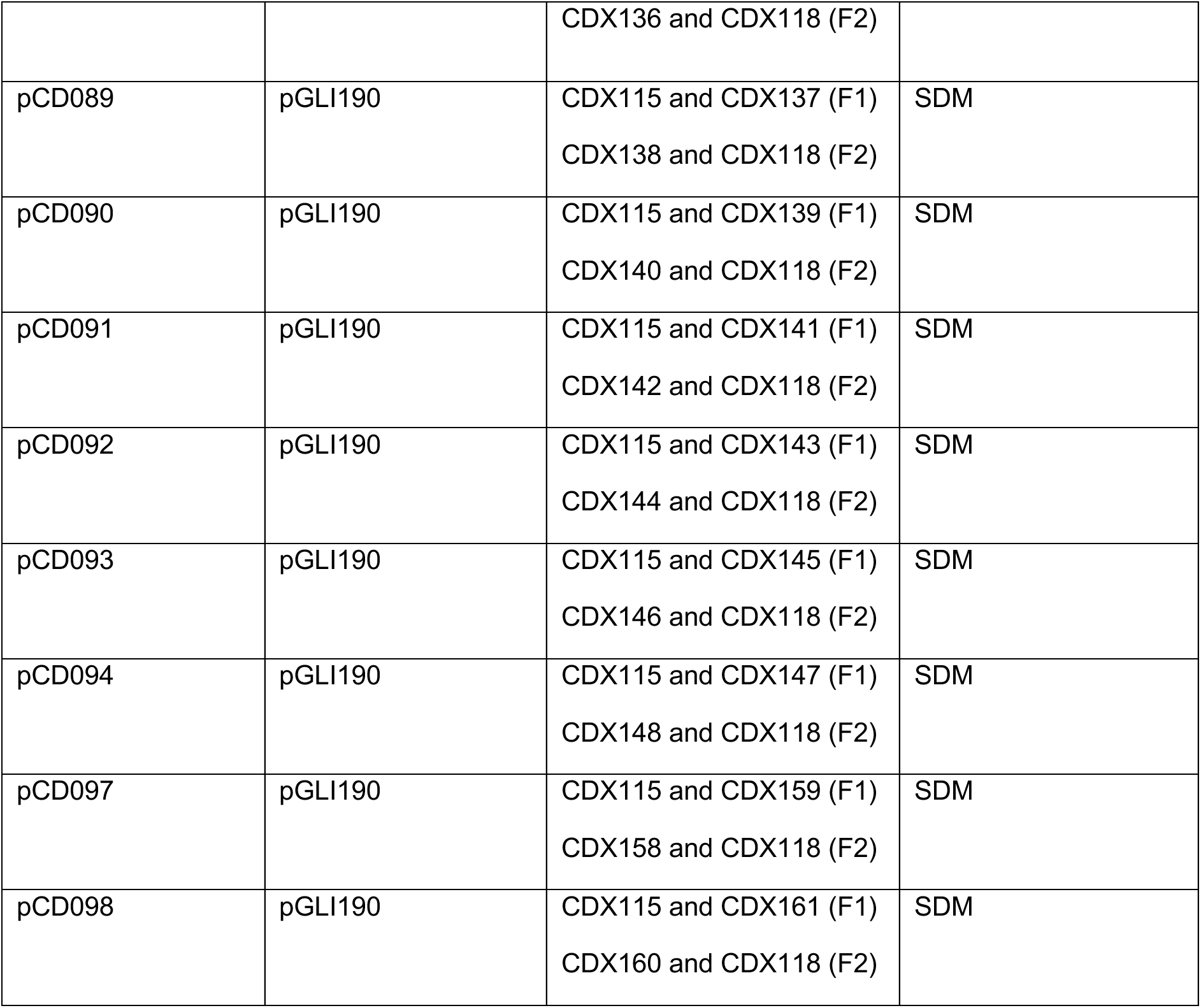
Details of plasmid construction.

**Table S4.** Plasmids used in this study.

| Plasmid ID | Details | Use of plasmid | Source |
| --- | --- | --- | --- |
| pBE1092 | Custom pET vector with N-ter His<sub>6</sub> tag | For cloning of expression plasmids | (4) |
| pDCE587 | Custom pET vector with C-ter His<sub>6</sub> tag | For cloning of expression plasmids | (4) |
| pGLI120 | pBE1092_YdbH-YdbL | Used as a template for pGLI139 construction | This study |
| pGLI139 | pBE1092_YdbH_YnbE_YdbL(1-22)_His<sub>6</sub>-TEV-YdbL(23-108) | Used as template for pGLI190 construction | This study |
| pGLI171 | pBE1092_His<sub>6</sub>-TEV-YdbL(22-108) | To express and purify full-length YdbL for structural studies (XRC and NMR). Signal sequence (1-21) removed for cytoplasmic expression before tag. | This study |
| pGLI190 | pDCE587_YdbL(1-22)_His<sub>6</sub>-TEV-YdbL(23-108) | To express and purify full-length YdbL for native MS and biochemical analyses. Signal sequence + 1AA before tag for periplasmic expression. | This study |
| pCD079 | pDCE587_YdbL(1-22)_His<sub>6</sub>-TEV_YdbL(23-108)_L68A | To express and purify mutant form of YdbL | This study |
| pCD080 | pDCE587_YdbL(1-22)_His<sub>6</sub>-TEV_YdbL(23-108)_L87A | To express and purify mutant form of YdbL | This study |
| pCD081 | pDCE587_YdbL(1-22)_His<sub>6</sub>-TEV_YdbL(23-108)_R61A | To express and purify mutant form of YdbL | This study |
| pCD082 | pDCE587_YdbL(1-22)_His<sub>6</sub>-TEV_YdbL(23-108)_R61K | To express and purify mutant form of YdbL | This study |
| pCD083 | pDCE587_YdbL(1-22)_His<sub>6</sub>-TEV_YdbL(23-108)_N72A | To express and purify mutant form of YdbL | This study |
| pCD084 | pDCE587_YdbL(1-22)_His<sub>6</sub>-TEV_YdbL(23-108)_E60A | To express and purify mutant form of YdbL | This study |
| pCD085 | pDCE587_YdbL(1-22)_His<sub>6</sub>-TEV_YdbL(23-108)_E60K | To express and purify mutant form of YdbL | This study |
| pCD086 | pDCE587_YdbL(1-22)_His<sub>6</sub>-TEV_YdbL(23-108)_K86A | To express and purify mutant form of YdbL | This study |
| pCD087 | pDCE587_YdbL(1-22)_His<sub>6</sub>-TEV_YdbL(23-108)_I79A | To express and purify mutant form of YdbL | This study |
| pCD088 | pDCE587_YdbL(1-22)_His<sub>6</sub>-TEV_YdbL(23-108)_R90A | To express and purify mutant form of YdbL | This study |
| pCD089 | pDCE587_YdbL(1-22)_His<sub>6</sub>-TEV_YdbL(23-108)_R90K | To express and purify mutant form of YdbL | This study |
| pCD090 | pDCE587_YdbL(1-22)_His<sub>6</sub>-TEV_YdbL(23-108)_Y65A | To express and purify mutant form of YdbL | This study |
| pCD091 | pDCE587_YdbL(1-22)_His<sub>6</sub>-TEV_YdbL(23-108)_L24A | To express and purify mutant form of YdbL | This study |
| pCD092 | pDCE587_YdbL(1-22)_His<sub>6</sub>-TEV_YdbL(23-108)_K107A | To express and purify mutant form of YdbL | This study |
| pCD093 | pDCE587_YdbL(1-22)_His<sub>6</sub>-TEV_YdbL(23-108)_R28A | To express and purify mutant form of YdbL | This study |
| pCD094 | pDCE587_YdbL(1-22)_His<sub>6</sub>-TEV_YdbL(23-108)_W104A | To express and purify mutant form of YdbL | This study |
| pCD097 | pDCE587_YdbL(1-22)_His<sub>6</sub>-TEV_YdbL(23-108)_E35A | To express and purify mutant form of YdbL | This study |
| pCD098 | pDCE587_YdbL(1-22)_His<sub>6</sub>-TEV_YdbL(23-108)_E35D | To express and purify mutant form of YdbL | This study |

**Table S5:**
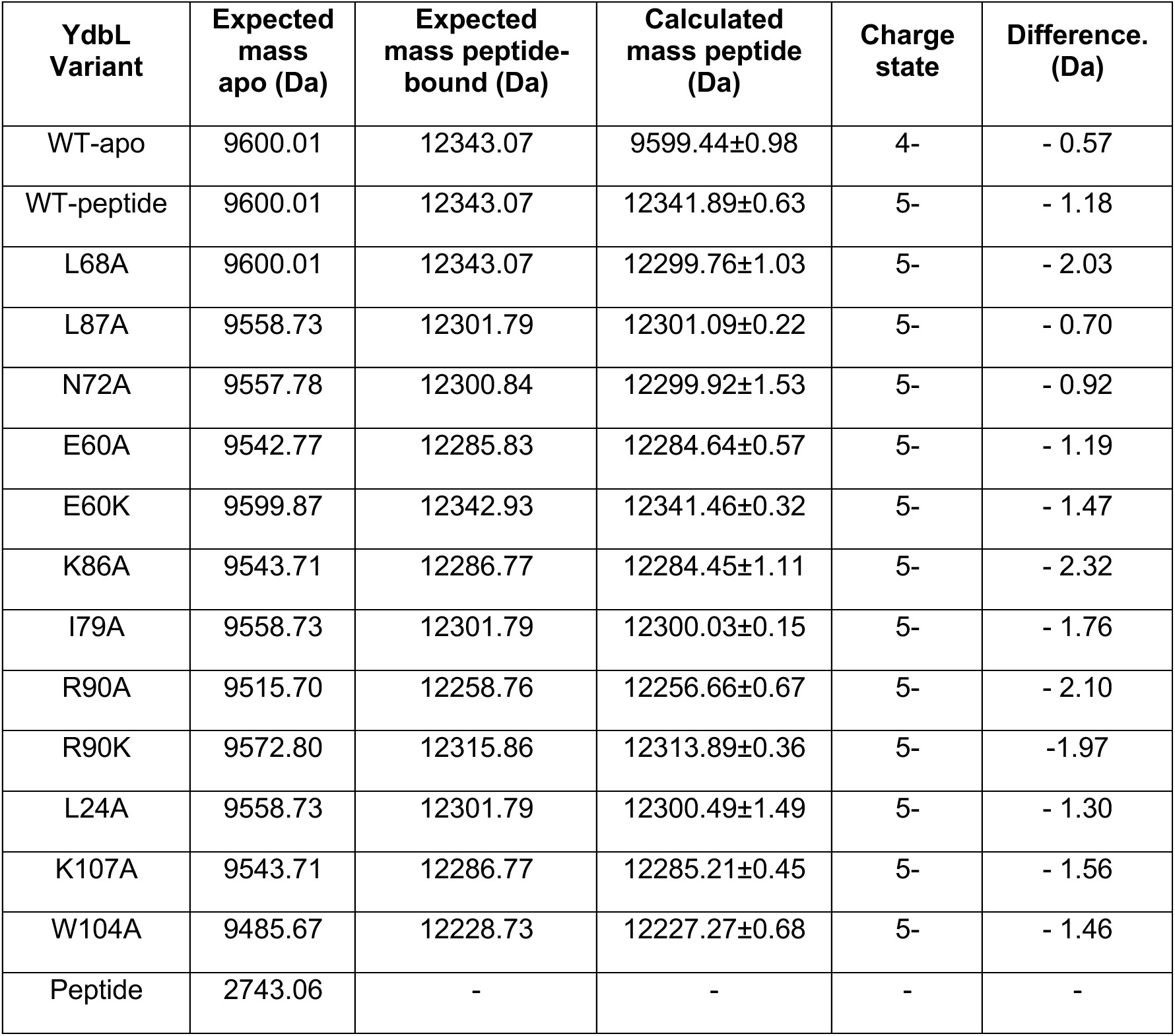
Expected versus calculated experimental masses for YdbL variants complexed with YnbE peptide by native MS.

**Table S6:** XRC data collection and refinement statistics.

|  | **YdbL** |
| --- | --- |
| **Wavelength** | 0.7749 |
| **Resolution range** | 53.22 - 2.12 (2.33 - 2.12) |
| **Space group** | I 4 2 2 |
| **Unit cell** | 63.994 63.994 95.851 90 90 90 |
| **Total reflections** | 62429 (3179) |
| **Unique reflections** | 5872 (1374) |
| **Multiplicity** | 10.5 (10.9) |
| **Completeness (%)** | 98.61 (95.09) |
| **Mean I/sigma(I)** | 4.6 (1) |
| **Wilson B-factor** | 32.99 |
| **R-merge** | 3.711 (1.373) |
| **R-meas** | 3.902 (1.447) |
| **R-pim** | 1.189 (0.452) |
| **CC1/2** | 0.861 (0.514) |
| **Reflections used in refinement** | 5872 (1374) |
| **Reflections used for R-free** | 510 (125) |
| **R-work** | 0.1992 (0.2921) |
| **R-free** | 0.2412 (0.3398) |
| **Number of non-hydrogen atoms** | 748 |
| **macromolecules** | 687 |
| **ligands** | 0 |
| **solvent** | 61 |
| **Protein residues** | 87 |
| **RMS(bonds)** | 0.006 |
| **RMS(angles)** | 0.64 |
| **Ramachandran favored (%)** | 100.00 |
| **Ramachandran allowed (%)** | 0.00 |
| **Ramachandran outliers (%)** | 0.00 |
| **Rotamer outliers (%)** | 0.00 |
| **Clashscore** | 0.72 |
| **Average B-factor** | 39.07 |
| **macromolecules** | 38.91 |
| **solvent** | 40.91 |

Statistics for the highest-resolution shell are shown in parentheses.

**Table S7:** SEC-SAXS experimental data after data reduction.

| **Sample details** |  |  |
| --- | --- | --- |
| Sample | **Cytoplasm-expressed YdbL (XRC sample)** | **Periplasm-expressed YdbL** |
| Source organism | *Escherichia coli* BW25113 | *Escherichia coli* BW25113 |
| Expression system | *Escherichia coli* T7 express | *Escherichia coli* T7 express |
| Molecular weight (Da) | 9,714 | 9,601 |
| Extinction coefficient (theoretical, M<sup>-1</sup> cm<sup>-1</sup>) | 11,460 | 11,460 |
| Loading concentration (mg/mL) | 9.9 | 4.8 |
| Buffer composition | 50 mM Tris pH 8, 150 mM NaCl, 1% Glycerol | 50 mM Tris pH 8, 150 mM NaCl, 1% Glycerol |
| **SAXS data collection parameters** |  |  |
| Beamline/detector | B21 Diamond Light Source, DECTRIS EIGER 4M | B21 Diamond Light Source, DECTRIS EIGER 4M |
| Energy (keV) | 13 | 13 |
| Beam size (μm) | < 75 | < 75 |
| Detector distance (m) | 3.7 | 3.7 |
| *q*-measurement range (Å<sup>-1</sup>) | 0.0045 - 0.34 | 0.0045 - 0.34 |
| Normalization | Transmitted intensity | Transmitted intensity |
| Exposures | 20 x 3 s | 20 x 3 s |
| Configuration (batch/SEC) | SEC | SEC |
| Column (flow rate, ml/min) | S75 3.2/300l precision column 0.075 mL/min | S75 3.2/300l precision column 0.075 mL/min |
| Experimental temperature (°C) | 22 | 22 |
| **Data processing/structural parameters** |  |  |
| SAXS Data Reduction | DAWN (Diamond Light Source) | DAWN (Diamond Light Source) |
| SAXS Data Processing | ScÅtter IV | ScÅtter IV |
| Buffer range (frames) | 424 - 456 | 422 - 454 |
| Main component (frames) | 18 - 2520 | 2 - 2149 |
| **Guinier Analysis** |  |  |
| $I(0)$ (cm<sup>-1</sup>) | 2.95E-2 | 1.29E-2 |
| $R_g$ (Å) | 13.9 | 13.8 |
| $qR_g$ range | 0.1423 - 1.2990 | 0.1054 - 1.2987 |
| $d_{max}$ | 40 | 43 |
| $R_c$ | 0.0 | 0.0 |
| $P_x$ | 4.0 | 4.0 |
| **$P(r)$ analysis** |  |  |
| $I(0)$ | 2.86E-2 | 1.24E-2 |
| $R_g$ (Å) | 13.84 | 13.79 |
| $d_{max}$ (Å) | 46 | 43 |
| $q$-range ($\text{\AA}^{-1}$) | 0.0045 - 0.34 | 0.0045 - 0.34 |
| Score | 1.125 | 1.178 |
| $V_p$ ($\text{\AA}^3$) | 13011 | 12890 |

**Table S8:**
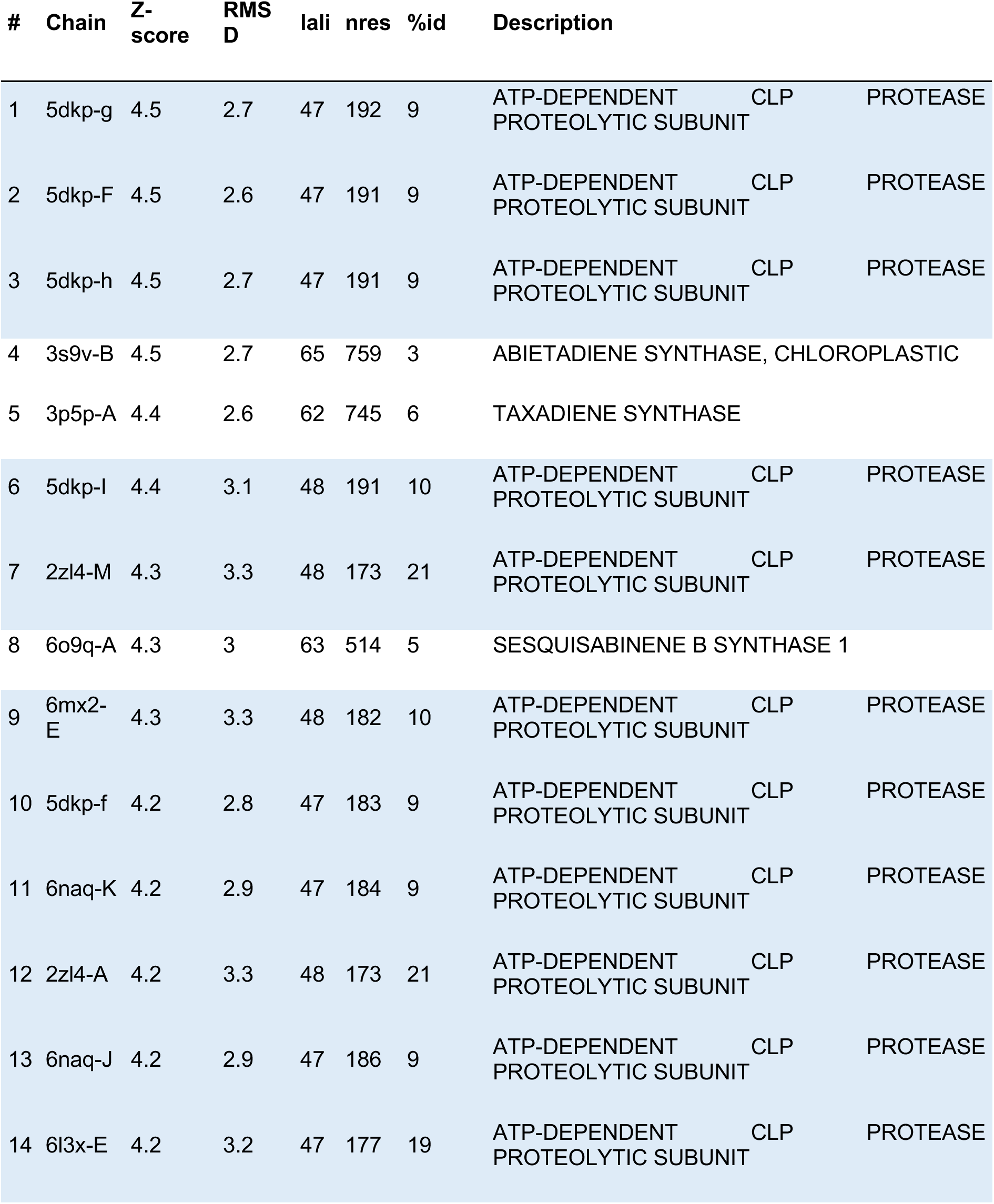
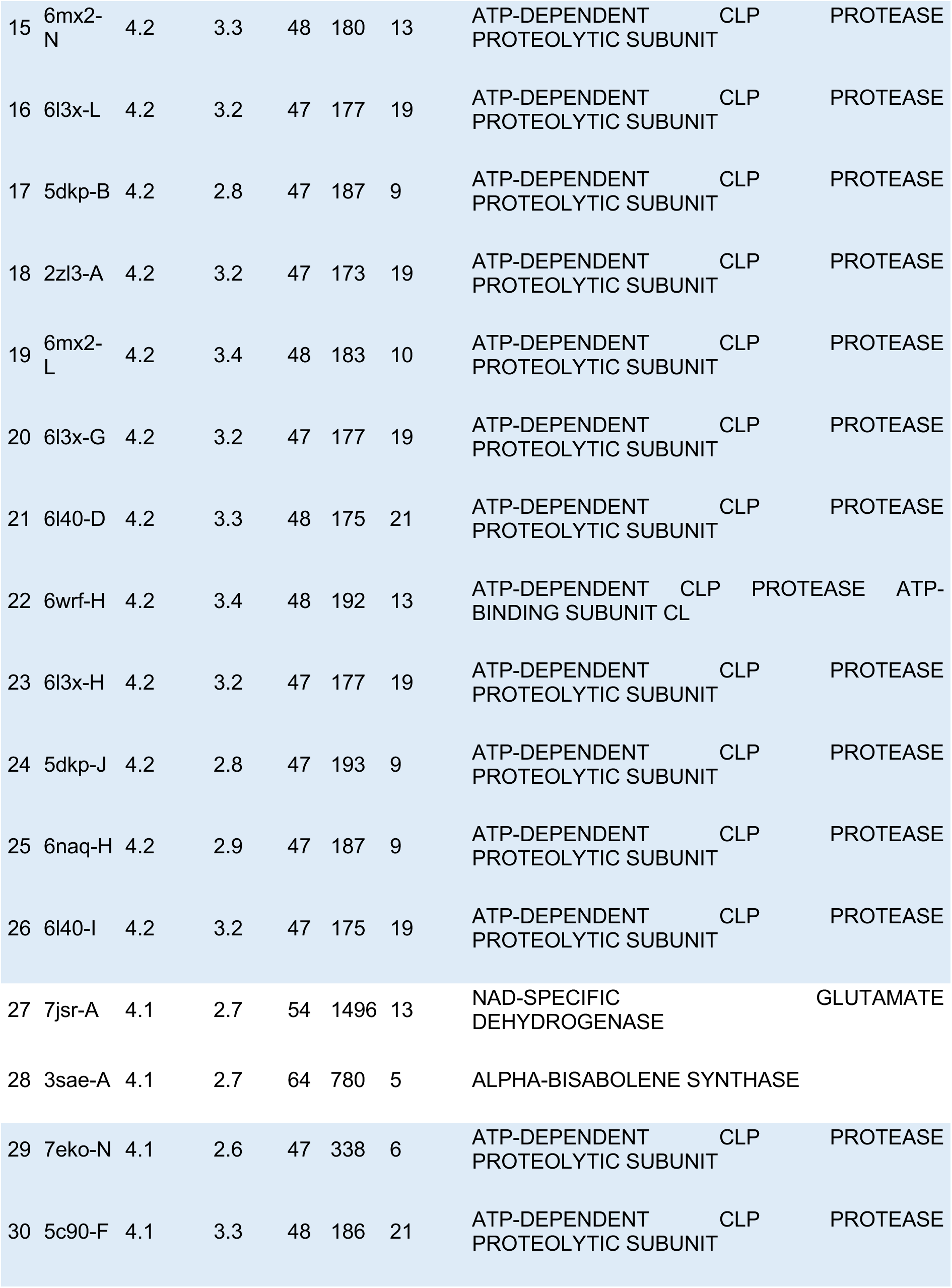
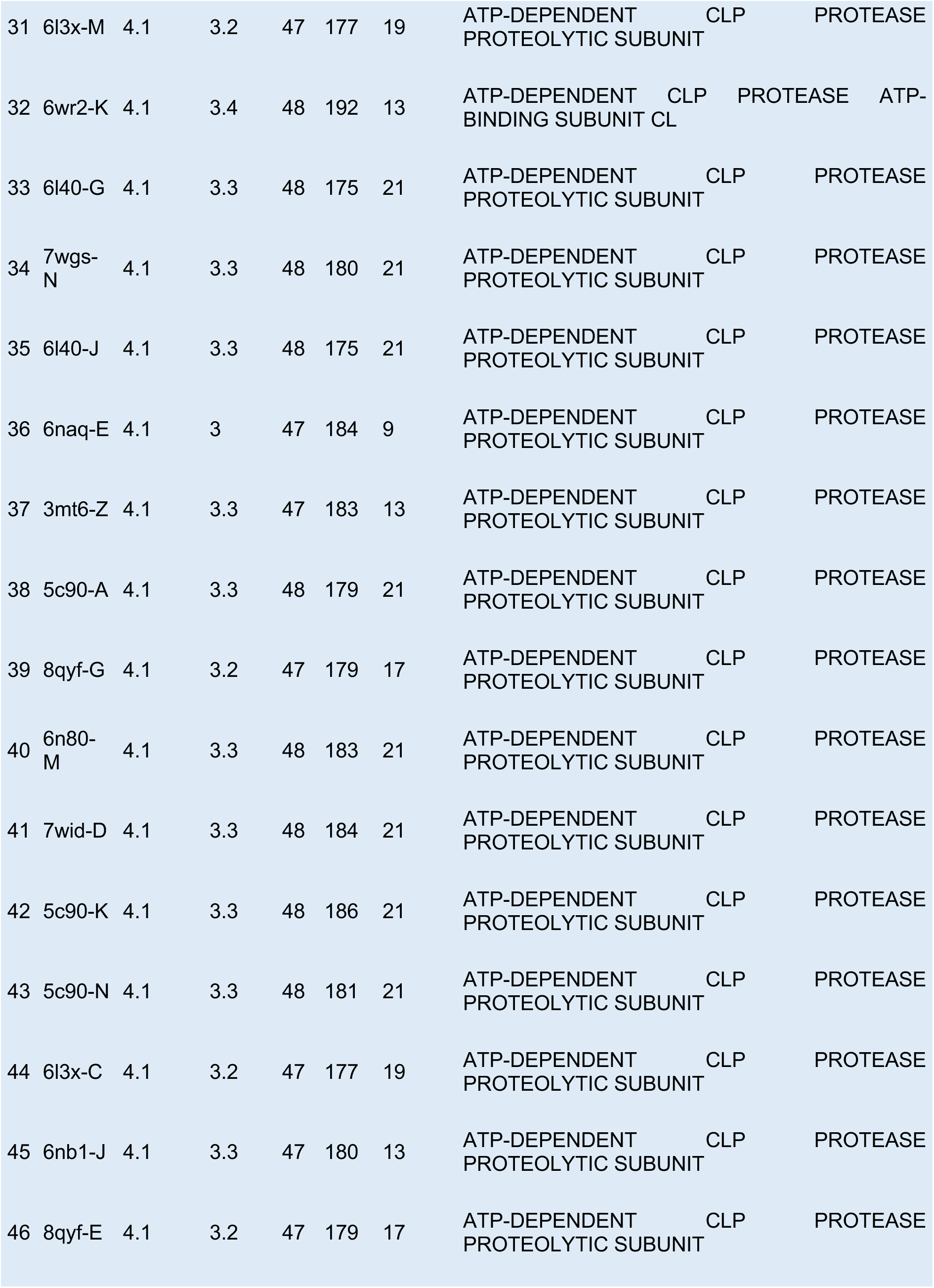
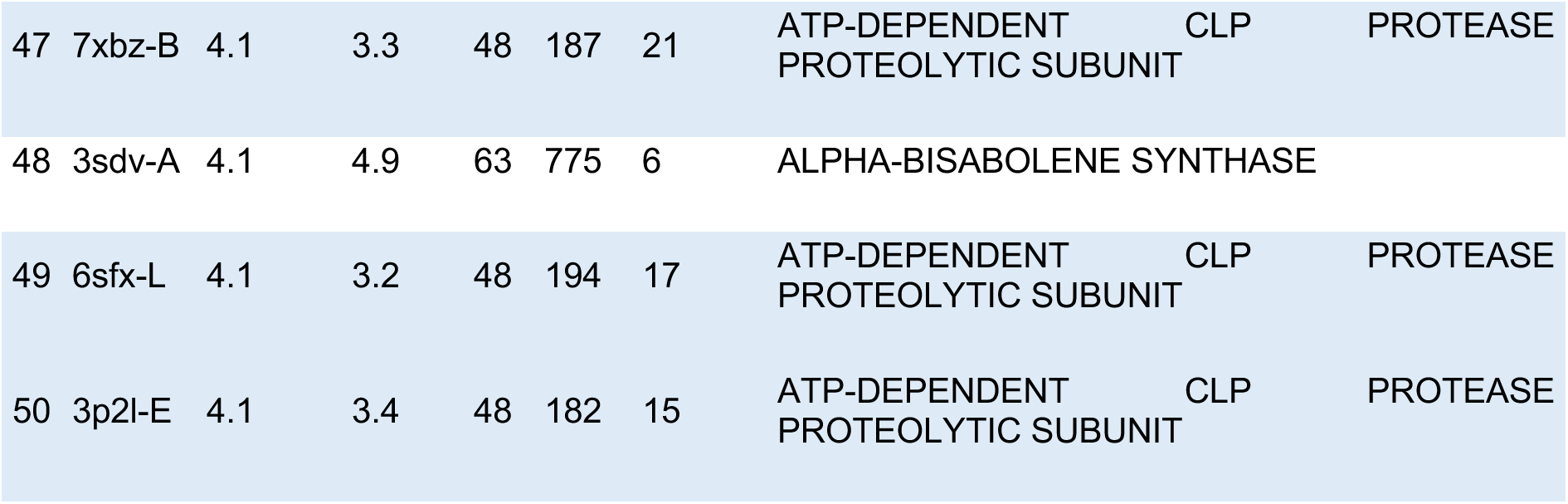
Dali v5 server results for YdbL model. Sorted by Z-score. Top 50 hits are shown of 1000. Blue shading depicts same significant hit with ATP-dependent Clp protease proteolytic subunit.

**Table S9:** Numerical amide backbone chemical shift changes ordered by distance of shift (ppm).

| **Residue** | **Amide shift between apo-YdbL and peptide-bound YdbL (ppm)** |
| --- | --- |
| L68 | 0.54654 |
| L87 | 0.48898 |
| R61 | 0.48451 |
| N72 | 0.4069 |
| E60 | 0.37511 |
| A83 | 0.34482 |
| K70 | 0.3023 |
| L24 | 0.28617 |
| A80 | 0.28498 |
| I79 | 0.28368 |
| Q66 | 0.27459 |
| G84 | 0.2717 |
| A69 | 0.26827 |
| S64 | 0.23965 |
| K62 | 0.22736 |
| Q71 | 0.22698 |
| V74 | 0.22613 |
| R90 | 0.22132 |
| Y65 | 0.21789 |
| T29 | 0.20932 |
| K81 | 0.19643 |
| A89 | 0.18748 |
| I57 | 0.18483 |
| A59 | 0.16876 |
| G34 | 0.16114 |
| L82 | 0.15266 |
| V97 | 0.14737 |
| L41 | 0.14318 |
| Y96 | 0.13412 |
| Q85 | 0.1216 |
| N73 | 0.11659 |
| V54 | 0.1132 |
| N58 | 0.09846 |
| S75 | 0.09804 |
| Y38 | 0.09524 |
| T36 | 0.09417 |
| E26 | 0.093 |
| E35 | 0.09283 |
| F37 | 0.09275 |
| K86 | 0.09002 |
| V33 | 0.0892 |
| V42 | 0.08761 |
| K92 | 0.0855 |
| V76 | 0.08472 |
| A63 | 0.08405 |
| G99 | 0.07971 |
| T23 | 0.07931 |
| Y40 | 0.07679 |
| D56 | 0.07551 |
| Q67 | 0.07475 |
| V88 | 0.07337 |
| D78 | 0.07218 |
| D25 | 0.06903 |
| R32 | 0.06719 |
| A91 | 0.0666 |
| F108 | 0.06479 |
| D77 | 0.05592 |
| L44 | 0.05531 |
| G39 | 0.05524 |
| A27 | 0.05345 |
| A43 | 0.05334 |
| G94 | 0.05285 |
| G31 | 0.04869 |
| A55 | 0.04702 |
| E51 | 0.04639 |
| K107 | 0.04445 |
| I100 | 0.04265 |
| G102 | 0.04184 |
| E95 | 0.04146 |
| T46 | 0.03985 |
| Q30 | 0.03175 |
| V105 | 0.03096 |
| N101 | 0.02656 |
| R28 | 0.02359 |
| T50 | 0.02176 |
| R106 | 0.02089 |
| K52 | 0.01752 |
| K103 | 0.01717 |
| W104 | 0.01554 |
| Q98 | 0.01434 |
| K45 | 0.01391 |
| A48 | 0.0134 |
| D47 | 0.01054 |
| L53 | 0.00803 |
| E49 | 0.00387 |

**Dataset S1 (separate file).** YdbL X-ray crystal structure is available on the PDB (accession code: 28OJ).

**Dataset S2 (separate file).** apo-YdbL assigned NMR chemical shifts can be found on the BMRB (accession code: 53587)

**Dataset S3 (separate file).** YnbE peptide-bound YdbL assigned NMR chemical shifts can be found on the BMRB (accession code: 53588).

## References

1. S. I. Miller, Antibiotic Resistance and Regulation of the Gram-Negative Bacterial Outer Membrane Barrier by Host Innate Immune Molecules. mBio 7, e01541–16 (2016).

2. T. J. Silhavy, D. Kahne, S. Walker, The Bacterial Cell Envelope. Cold Spring Harb Perspect Biol 2, a000414 (2010).

3. J. C. Henderson, et al., The Power of Asymmetry: Architecture and Assembly of the Gram-Negative Outer Membrane Lipid Bilayer. Annu Rev Microbiol 70, 255–278 (2016).

4. G. Benn, et al., Phase separation in the outer membrane of Escherichia coli. Proceedings of the National Academy of Sciences 118, e2112237118 (2021).

5. G. Benn, et al., OmpA controls order in the outer membrane and shares the mechanical load. Proceedings of the National Academy of Sciences 121, e2416426121 (2024).

6. M. N. Webby, et al., Lipids mediate supramolecular outer membrane protein assembly in bacteria. Science Advances 8, eadc9566 (2022).

7. J. Grimm, et al., The inner membrane protein YhdP modulates the rate of anterograde phospholipid flow in Escherichia coli. PNAS 117, 26907–26914 (2020).

8. N. Ruiz, R. M. Davis, S. Kumar, YhdP, TamB, and YdbH Are Redundant but Essential for Growth and Lipid Homeostasis of the Gram-Negative Outer Membrane. mBio 12, e02714–21 (2021).

9. M. V. Douglass, A. B. McLean, M. S. Trent, Absence of YhdP, TamB, and YdbH leads to defects in glycerophospholipid transport and cell morphology in Gram-negative bacteria. PLOS Genetics 18, e1010096 (2022).

10. S. D. Neuman, T. P. Levine, A. Bashirullah, A novel superfamily of bridge-like lipid transfer proteins. Trends in Cell Biology 32, 962–974 (2022).

11. S. Kumar, N. Ruiz, Bacterial AsmA-Like Proteins: Bridging the Gap in Intermembrane Phospholipid Transport. Contact 6, 25152564231185931 (2023).

12. Z. Wang, et al., An allosteric transport mechanism for the AcrAB-TolC multidrug efflux pump. eLife 6, e24905 (2017).

13. E. F. Semeraro, J. M. Devos, L. Porcar, V. T. Forsyth, T. Narayanan, In vivo analysis of the Escherichia coli ultrastructure by small-angle scattering. IUCrJ 4, 751–757 (2017).

14. V. R. F. Matias, A. Al-Amoudi, J. Dubochet, T. J. Beveridge, Cryo-Transmission Electron Microscopy of Frozen-Hydrated Sections of Escherichia coli and Pseudomonas aeruginosa. Journal of Bacteriology 185, 6112–6118 (2003).

15. B. F. Cooper, et al., Phospholipid Transport Across the Bacterial Periplasm Through the Envelope-spanning Bridge YhdP. Journal of Molecular Biology 437, 168891 (2025).

16. J. Selkrig, et al., Discovery of an archetypal protein transport system in bacterial outer membranes. Nat Struct Mol Biol 19, 506–510 (2012).

17. J. Selkrig, et al., Conserved features in TamA enable interaction with TamB to drive the activity of the translocation and assembly module. Sci Rep 5, 12905 (2015).

18. B. Yang, et al., Structural basis of outer membrane biogenesis by the TamAB translocase. Nat Commun 17, 437 (2026).

19. S. Kumar, R. M. Davis, N. Ruiz, YdbH and YnbE form an intermembrane bridge to maintain lipid homeostasis in the outer membrane of Escherichia coli. Proceedings of the National Academy of Sciences 121, e2321512121 (2024).

20. D. Sposato, et al., Structure–function relationship of the Pseudomonas aeruginosa AsmA-like proteins YhdP and YdbH involved in outer membrane biogenesis. Protein Science 34, e70227 (2025).

21. L. M. Guzman, D. Belin, M. J. Carson, J. Beckwith, Tight regulation, modulation, and high-level expression by vectors containing the arabinose PBAD promoter. J Bacteriol 177, 4121–4130 (1995).

22. J. W. Chin, A. B. Martin, D. S. King, L. Wang, P. G. Schultz, Addition of a photocrosslinking amino acid to the genetic code of Escherichia coli. Proceedings of the National Academy of Sciences 99, 11020–11024 (2002).

23. J. Sanchez-Weatherby, et al., VMXi: a fully automated, fully remote, high-flux in situ macromolecular crystallography beamline. J Synchrotron Rad 26, 291–301 (2019).

24. H. Mikolajek, et al., Protein-to-structure pipeline for ambient-temperature in situ crystallography at VMXi. IUCrJ 10, 420–429 (2023).

25. J. Sandy, H. Mikolajek, A. J. Thompson, J. Sanchez-Weatherby, M. A. Hough, Crystallization and In Situ Room Temperature Data Collection Using the Crystallization Facility at Harwell and Beamline VMXi, Diamond Light Source. J Vis Exp (2024). 10.3791/65964.

26. E. Krissinel, K. Henrick, Inference of Macromolecular Assemblies from Crystalline State. Journal of Molecular Biology 372, 774–797 (2007).

27. D. R. Armstrong, et al., PDBe: improved findability of macromolecular structure data in the PDB. Nucleic Acids Res 48, D335–D343 (2020).

28. L. Holm, A. Laiho, P. Törönen, M. Salgado, DALI shines a light on remote homologs: One hundred discoveries. Protein Science 32, e4519 (2023).

29. J. D. Goodreid, et al., Development and Characterization of Potent Cyclic Acyldepsipeptide Analogues with Increased Antimicrobial Activity. J. Med. Chem. 59, 624–646 (2016).

30. S. A. Mahmoud, P. Chien, Regulated Proteolysis in Bacteria. Annu Rev Biochem 87, 677–696 (2018).

31. M. F. Mabanglo, W. A. Houry, Recent structural insights into the mechanism of ClpP protease regulation by AAA+ chaperones and small molecules. J Biol Chem 298, 101781 (2022).

32. L. J. McGuffin, et al., Prediction and quality assessment of protein quaternary structure models using the MultiFOLD2 and ModFOLDdock2 servers. Nucleic Acids Res 53, W472–W477 (2025).

33. L. J. McGuffin, et al., Prediction of protein structures, functions and interactions using the IntFOLD7, MultiFOLD and ModFOLDdock servers. Nucleic Acids Res 51, W274–W280 (2023).

34. Y. Shen, A. Bax, Protein backbone and sidechain torsion angles predicted from NMR chemical shifts using artificial neural networks. J Biomol NMR 56, 227–241 (2013).

35. H. Ashkenazy, et al., ConSurf 2016: an improved methodology to estimate and visualize evolutionary conservation in macromolecules. Nucleic Acids Res 44, W344–W350 (2016).

36. M. J. Pallen, B. W. Wren, The HtrA family of serine proteases. Mol Microbiol 26, 209–221 (1997).

37. C. Spiess, A. Beil, M. Ehrmann, A Temperature-Dependent Switch from Chaperone to Protease in a Widely Conserved Heat Shock Protein. Cell 97, 339–347 (1999).

38. T. Clausen, C. Southan, M. Ehrmann, The HtrA family of proteases: implications for protein composition and cell fate. Mol Cell 10, 443–455 (2002).

39. D. Y. Kim, K. K. Kim, Structure and function of HtrA family proteins, the key players in protein quality control. J Biochem Mol Biol 38, 266–274 (2005).

40. X. Ge, et al., DegP primarily functions as a protease for the biogenesis of β-barrel outer membrane proteins in the Gram-negative bacterium Escherichia coli. The FEBS Journal 281, 1226–1240 (2014).

41. T. Krojer, et al., Interplay of PDZ and protease domain of DegP ensures efficient elimination of misfolded proteins. Proceedings of the National Academy of Sciences 105, 7702–7707 (2008).

42. T. Krojer, J. Sawa, R. Huber, T. Clausen, HtrA proteases have a conserved activation mechanism that can be triggered by distinct molecular cues. Nat Struct Mol Biol 17, 844–852 (2010).

43. S. Kim, R. T. Sauer, Cage assembly of DegP protease is not required for substrate-dependent regulation of proteolytic activity or high-temperature cell survival. Proceedings of the National Academy of Sciences 109, 7263–7268 (2012).

44. A. K. Rai, et al., Genetic evidence for functional diversification of gram-negative intermembrane phospholipid transporters. PLOS Genetics 20, e1011335 (2024).

45. D. Sposato, et al., Redundant essentiality of AsmA-like proteins in Pseudomonas aeruginosa. mSphere 0, e00677–23 (2024).

46. T. Baba, et al., Construction of Escherichia coli K-12 in-frame, single-gene knockout mutants: the Keio collection. Mol Syst Biol 2, MSB4100050 (2006).

47. K. A. Datsenko, B. L. Wanner, One-step inactivation of chromosomal genes in Escherichia coli K-12 using PCR products. Proceedings of the National Academy of Sciences 97, 6640–6645 (2000).

48. D. Yu, et al., An efficient recombination system for chromosome engineering in Escherichia coli. Proceedings of the National Academy of Sciences 97, 5978–5983 (2000).

49. B. W. Simpson, et al., Identification of Residues in the Lipopolysaccharide ABC Transporter That Coordinate ATPase Activity with Extractor Function. mBio 7, e01729–16 (2016).

50. Y. Ryu, P. G. Schultz, Efficient incorporation of unnatural amino acids into proteins in Escherichia coli. Nat Methods 3, 263–265 (2006).

51. F. Teufel, et al., SignalP 6.0 predicts all five types of signal peptides using protein language models. Nat Biotechnol 40, 1023–1025 (2022).

52. G. Winter, xia2: an expert system for macromolecular crystallography data reduction. J Appl Cryst 43, 186–190 (2010).

53. G. Winter, et al., DIALS as a toolkit. Protein Science 31, 232–250 (2022).

54. R. J. Gildea, et al., xia2.multiplex: a multi-crystal data-analysis pipeline. Acta Cryst D 78, 752–769 (2022).

55. B. W. Matthews, Solvent content of protein crystals. J Mol Biol 33, 491–497 (1968).

56. K. A. Kantardjieff, B. Rupp, Matthews coefficient probabilities: Improved estimates for unit cell contents of proteins, DNA, and protein–nucleic acid complex crystals. Protein Sci 12, 1865–1871 (2003).

57. A. J. McCoy, et al., Phaser crystallographic software. J Appl Cryst 40, 658–674 (2007).

58. P. V. Afonine, et al., Towards automated crystallographic structure refinement with phenix.refine. Acta Cryst D 68, 352–367 (2012).

59. P. Emsley, K. Cowtan, Coot: model-building tools for molecular graphics. Acta Cryst D 60, 2126–2132 (2004).

60. M. Varadi, et al., AlphaFold Protein Structure Database: massively expanding the structural coverage of protein-sequence space with high-accuracy models. Nucleic Acids Research 50, D439–D444 (2022).

61. M. Varadi, et al., AlphaFold Protein Structure Database in 2024: providing structure coverage for over 214 million protein sequences. Nucleic Acids Research 52, D368–D375 (2024).

62. A. Fadini, et al., Highlights of Model Quality Assessment in CASP16. Proteins: Structure, Function, and Bioinformatics **n/a** (2025).

63. F. Glaser, et al., ConSurf: identification of functional regions in proteins by surface-mapping of phylogenetic information. Bioinformatics 19, 163–164 (2003).

64. M. Landau, et al., ConSurf 2005: the projection of evolutionary conservation scores of residues on protein structures. Nucleic Acids Res 33, W299–302 (2005).

65. H. Ashkenazy, E. Erez, E. Martz, T. Pupko, N. Ben-Tal, ConSurf 2010: calculating evolutionary conservation in sequence and structure of proteins and nucleic acids. Nucleic Acids Res 38, W529–533 (2010).

66. G. Celniker, et al., ConSurf: Using evolutionary data to raise testable hypotheses about protein function. Israel Journal of Chemistry 53, 199–206 (2013).

67. B. Yariv, et al., Using evolutionary data to make sense of macromolecules with a “face-lifted” ConSurf. Protein Sci 32, e4582 (2023).

68. L. Holm, P. Rosenström, Dali server: conservation mapping in 3D. Nucleic Acids Res 38, W545–W549 (2010).

69. L. Holm, L. M. Laakso, Dali server update. Nucleic Acids Research 44, W351–W355 (2016).

70. L. Holm, Dali server: structural unification of protein families. Nucleic Acids Research 50, W210–W215 (2022).

71. N. P. Cowieson, et al., Beamline B21: high-throughput small-angle X-ray scattering at Diamond Light Source. J Synchrotron Radiat 27, 1438–1446 (2020).

72. D. Schneidman-Duhovny, M. Hammel, J. A. Tainer, A. Sali, Accurate SAXS Profile Computation and its Assessment by Contrast Variation Experiments. Biophysical Journal 105, 962–974 (2013).

73. D. Schneidman-Duhovny, M. Hammel, J. A. Tainer, A. Sali, FoXS, FoXSDock and MultiFoXS: Single-state and multi-state structural modeling of proteins and their complexes based on SAXS profiles. Nucleic Acids Research 44, W424–W429 (2016).

74. C. Redfield, “Assignment of Protein NMR Spectra Using Heteronuclear NMR—A Tutorial” in Protein NMR: Modern Techniques and Biomedical Applications, L. Berliner, Ed. (Springer US, 2015), pp. 1–42.

75. T. Schulte-Herbrüggen, O. W. Sørensen, Clean TROSY: Compensation for Relaxation-Induced Artifacts. Journal of Magnetic Resonance 144, 123–128 (2000).

76. E. Lescop, P. Schanda, B. Brutscher, A set of BEST triple-resonance experiments for time-optimized protein resonance assignment. Journal of Magnetic Resonance 187, 163–169 (2007).

77. F. Delaglio, et al., NMRPipe: a multidimensional spectral processing system based on UNIX pipes. J Biomol NMR 6, 277–293 (1995).

78. S. G. Hyberts, A. G. Milbradt, A. B. Wagner, H. Arthanari, G. Wagner, Application of iterative soft thresholding for fast reconstruction of NMR data non-uniformly sampled with multidimensional Poisson Gap scheduling. J Biomol NMR 52, 315–327 (2012).

79. W. F. Vranken, et al., The CCPN data model for NMR spectroscopy: development of a software pipeline. Proteins 59, 687–696 (2005).

## SI References

1. F. R. Blattner, et al., The complete genome sequence of Escherichia coli K-12. Science 277, 1453–1462 (1997).

2. N. Ruiz, R. M. Davis, S. Kumar, YhdP, TamB, and YdbH Are Redundant but Essential for Growth and Lipid Homeostasis of the Gram-Negative Outer Membrane. mBio 12, e02714–21 (2021).

3. S. Kumar, R. M. Davis, N. Ruiz, YdbH and YnbE form an intermembrane bridge to maintain lipid homeostasis in the outer membrane of Escherichia coli. Proceedings of the National Academy of Sciences 121, e2321512121 (2024).

4. D. C. Ekiert, et al., Architectures of Lipid Transport Systems for the Bacterial Outer Membrane. Cell 169, 273–285.e17 (2017).

